# Human eIF3 and eIF5 coordinate AUG recognition to sustain energy homeostasis

**DOI:** 10.64898/2026.11.21.768280

**Authors:** Filip Trčka, Terezie Prilepskaja, Anna M. Smirnova, Rosslyn Grosely, Laurelle Lee Young, Jana Vojtová, Mahabub Pasha Mohammad, Pragya Kamal, Solomon A. Haizel, Klára Frišová, Susan Wagner, Jan Jelínek, Joseph D. Puglisi, Christine Vogel, Anna Herrmannová, Leoš Shivaya Valášek

## Abstract

Precise translation initiation is essential for efficient production of proteins; missing or mis-selecting the canonical start site can have detrimental consequences. Past research identified several initiation factors governing start codon selection accuracy in yeast, but the increasing complexity of higher eukaryotes have limited similar research in mammalian cells. We present an *in cellulo* study of knock-in mutants of the eIF3c subunit of the multitasking eIF3 complex that disrupt eIF3’s interaction with eIF5, thereby preventing functional binding of eIF5 to pre-initiation complexes. This selectively relaxes start site selection stringency and reduces initiation sensitivity, leading to altered translation efficiency for nearly 600 proteins. Proteins with N-terminal targeting signals were among those especially affected, resulting in impaired mitochondrial function. These findings establish the first direct causal link between an initiation factor, stringent start site recognition, and impaired mitochondrial respiration and glycolytic activity, implicating the eIF3–eIF5 interaction as a key regulator of energy homeostasis.

## INTRODUCTION

The fidelity of transcription and translation ensures the faithful transmission of genetic information from life-perpetuating DNA into functional protein. The accuracy of each step requires precise biochemical discrimination between individual nucleotides or triplet codons. Translation is known to be the most error prone and energetically costly step of gene expression^1,2^. On the other hand, unlike mis-replication errors, mis-translation events are only transiently encoded in protein products and can be readily degraded unless they are intentional^3^. The situation changes dramatically with frequent mis-initiation events at the wrong translation initiation sites (TIS), which generate random and often toxic polypeptides. Therefore, the TIS selection, setting the reading frame for decoding, is the rate limiting step of the entire translational cycle^4^.

The center player in this process is the initiator methionyl-tRNA (Met-tRNA_i_^Met^) whose 3’ UAC 5’ anticodon is fully complementary to the most frequent TIS, AUG. Note that initiation also can start from near-cognate TISs (e.g., CUG, UUG, GUG, ACG, AAG, AGG, and others)^5^, but it is much less efficient with often regulatory roles^4,6^. For example, near-cognate TISs are the source of N-terminal variants of proteins with different targeting in the cell^7^.

During initiation from any TIS, Met-tRNA_i_^Met^ is delivered to the 40S subunit in the ternary complex (eIF2-TC) together with eIF2 bound to GTP in a reaction promoted by eIF1A, eIF1, eIF3, and at least in yeast also eIF5^8–10^. Thus, formed 43S pre-initiation complex (PIC) adopts an open, scanning conducive conformation for mRNA binding and subsequent scanning for TIS. eIF4G and eIF3 represent the major driving force of mRNA recruitment. The resulting 48S PIC mainly utilizes the N- and C-terminal tails (NTTs and CTTs) of eIF1A and eIF1 and the CTT of eIF2β to monitor partial (P^out^) *versus* complete (P^in^) accommodation of Met-tRNA_i_^Met^ in the ribosomal P site. This depends on the degree of complementarity between the codon currently scanned and the UAC anticodon^11–13^.

Upon establishment of a TIS/anticodon stable base-pairing, a series of rearrangements orchestrated by NTTs of eIF1A and eIF1 prompts the 48S PICs to reestablish a closed, non-processive conformation, thereby arresting scanning^14–21^. At the same time, eIF5 binds to the P site where it replaces eIF1 and activates GTP hydrolysis on eIF2-TC, thereby rendering the whole process irreversible^14^. Note that yeast eIF5 was shown to be part of the scanning complex right from its assembly and to stimulate reversible GTP hydrolysis on its way to TIS, with eIF1 preventing the free Pi release until the proper TIS has been recognized in the favorable Kozak sequence context^16,22,23^. However, our recent single molecule experiments suggested that human eIF5 stably joins the scanning complex not before its arrest at TIS^24^. Given that most of our current knowledge comes from yeast genetics, which have, for example, much less complex eIF3^9^, we can expect to find many more differences.

To complete the initiation phase, the closure of the 40S mRNA binding channel triggers ejection of most of eIFs except for at least eIF4G and eIF3^12,25–29^, allowing the 60S subunit joining step to produce an elongation-competent 80S ribosome.

eIF3 is the most complex initiation factor, which also promotes translation termination, ribosomal recycling, stop codon readthrough, re-initiation and, importantly, mRNA-specific translational control (reviewed in^8,30,31^). Human eIF3 is composed of altogether 12 subunits, 8 of which (namely eIF3a, c, e, f, h, k, l, and m) form the so-called PCI/MPN octamer, whereas eIF3b, g, i, and the C-terminal half of eIF3a assemble into the so-called Yeast-Like-Core (YLC)^32–36^ (Figure 1A). In contrast, the *S. cerevisiae* eIF3 complex has only 5 subunits^9^. Earlier yeast genetics experiments revealed that eIF3 interacts with eIF1 and eIF5 – the key eIFs ensuring accurate TIS recogniton^37–44^ – through its eIF3c subunits (Nip1p)^37,39,43,45–51^. In spite of that, its contribution to this intricate process remains unexplored.

**Figure 1.**
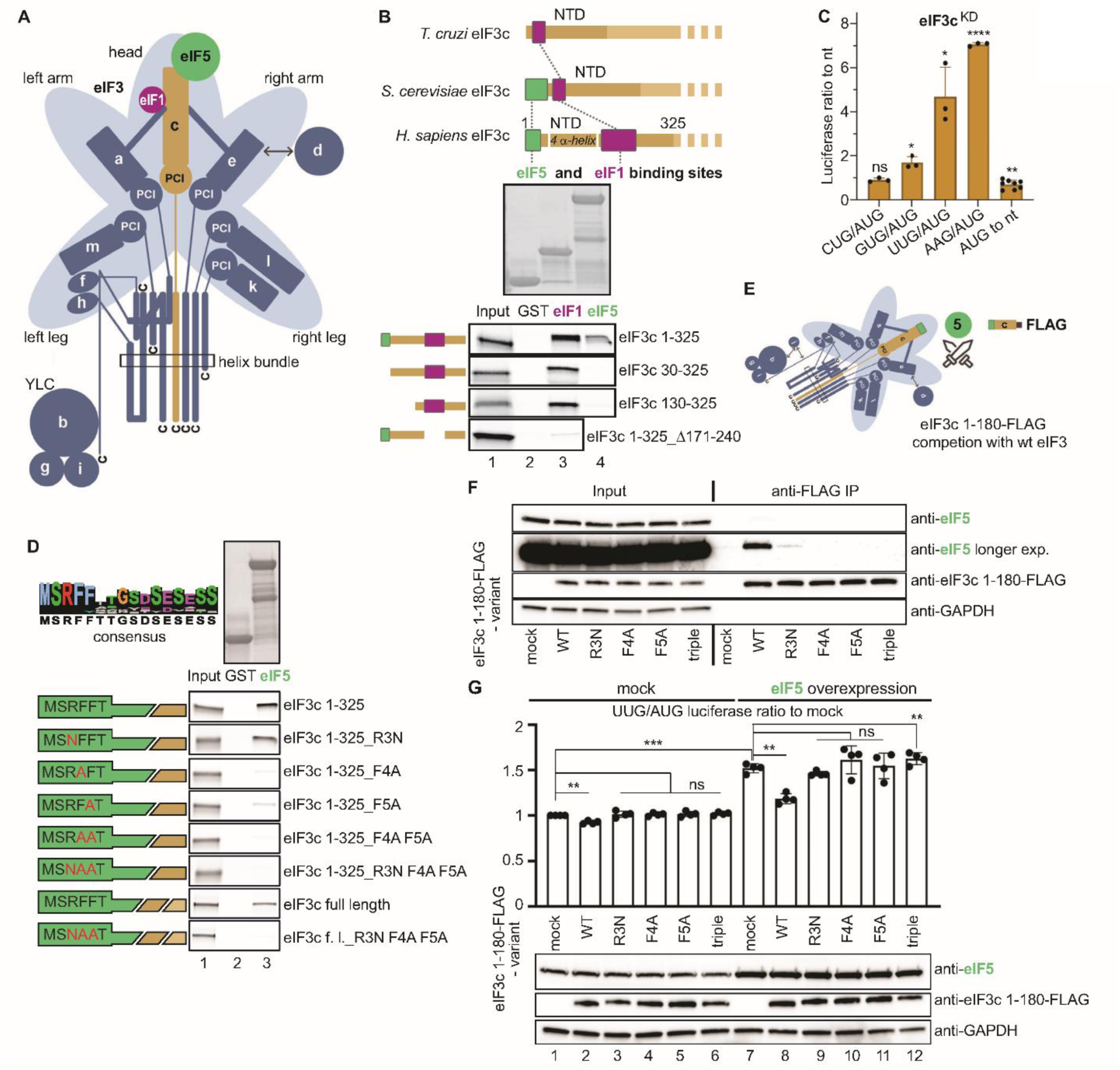
The eIF3c–eIF5 interaction is mediated by three N-terminal eIF3c residues. **(A)** Schematic model of the human eIF3 complex showing the PCI/MPN octamer on a light blue background, the yeast-like core (YLC), and the relative positions of eIF1 (magenta) and eIF5 (green) on the eIF3c N-terminal domain (ochre); adapted from^36^. **(B)** Mapping of the binding sites for eIF1 and eIF5 within eIF3c-NTD. The top panel shows the relative positions of the binding sites for eIF5 and eIF1 in eIF3c in *T. cruzi, S. cerevisiae,* and *H. sapiens*; adapted from^13,44^. The lower panels show GST pull-down analysis of the indicated radiolabeled human eIF3c fragments with GST alone, GST-eIF1 or GST-eIF5. Lane 1 contains 50% of the input amounts of *in vitro* translated proteins added to each reaction. The schematic of the fragments used is shown on the left. **(C)** Downregulation of eIF3c relaxes the stringency of the TIS selection *in vivo*. HeLa cells were transfected with the non-targeting (nt) or eIF3c siRNAs, and 48 h later, the second round of transfection with individual reporters (AUG or CUG, GUG, UUG, AAG) was carried out. Luciferase activities were measured in lysate 24 h after the second transfection. mRNA levels of each reporter in the same lysate were measured by qPCR and used for normalization. Plot shows the mean ratio ± S.D. of the near-cognate initiation rates relative to those of AUG; the ratio in the nt sample was set to 1. Dots represent results from individual biological replicates (n ≥ 3). One sample t test was used for statistical evaluation (* p ≤ 0.05, ** p ≤ 0.01, *** p ≤ 0.001, **** p ≤ 0.0001, ns = not significant). **(D)** The conserved R3, F4, and F5 residues at the eIF3c-NTD mediate binding to eIF5. The sequence illustrates conservation of the eIF3c N-terminal tip (for details, see Figure S1E). GST pull-down analysis was performed with the indicated radiolabeled human eIF3c 1-325 or full-length (f. l.) constructs and GST alone or GST-eIF5. Lane 1 contains 50% of the input amounts of *in vitro* translated proteins added to each reaction. The schematic of the fragments used is shown on the left. The substituted residues are highlighted in red. **(E)** Schematic model of competition between endogenous eIF3 and the eIF3c-FLAG fragment for eIF5 *in vivo*. **(f)** Competition between endogenous eIF3 and wt or mutant eIF3c-FLAG fragments for eIF5 *in vivo*. The C-terminally FLAG-tagged N-terminal fragments of eIF3c (1-180 aa) with indicated point mutations were overexpressed in HEK293T cells and immunoprecipitated using anti-FLAG beads. The levels of eIF5 and FLAG-tagged eIF3c fragments were detected by western blotting. (**G**) The wt but not mutant eIF3c-FLAG fragments eliminate the eIF5-overexpression effect on stringent TIS selection. The eIF3c fragments, same as in (f), were co-transfected either with AUG-or UUG-initiated firefly luciferase reporters, and either an eIF5-overexpressing vector or an empty vector (mock); renilla luciferase reporter was used for transfection efficiency normalization. Plot shows the mean ratio ± S.D. of UUG-mediated initiation rates relative to those of AUG; this ratio in the mock sample was set to 1. Dots represent results from individual biological replicates (n=4). Statistical significance was determined by One sample t test (* p ≤ 0.05, ** p ≤ 0.01, *** p ≤ 0.001). The levels of overexpressed eIF3c fragments and eIF5 were monitored by western blotting.

Here, we determined that human eIF3c interacts with eIF1 and eIF5 *via* two distinct N-terminal segments and mapped the precise contact points between eIF3c with eIF5. Using CRISPR/Cas9, we introduced the N-terminal disruptive substitutions directly into the eIF3c gene and examined the effects of the eIF3-eIF5 interaction impairment using genetic, biochemical, omics and single-molecule approaches. Our eIF3c-NTD single and triple point mutants, which represent the first genetically defined knock-in system for studying translation initiation in higher eukaryotes, have revealed that eIF3 promotes the stable binding of eIF5 to the scanning-arrested PICs and co-ordinates its role therein during TIS recognition. Impairment of this interaction altered translation efficiency for nearly 600 proteins, among which we found proteins bearing N-terminal targeting signal domains, such as nuclear-encoded mitochondrial proteins. Collectively, the effects observed lead to a complex phenotype, exemplified by impaired mitochondrial function and overall metabolic status.

## RESULTS

### eIF5 interacts with the eIF3c-NTD tip – mapping interactions between human eIFs

A systematic analysis of mutual contacts between various eIFs was carried out in *S. cerevisiae* but never in much more complex mammals (reviewed in^8,11^). Therefore, we subjected all 12 subunits of human eIF3 to a comprehensive *in vitro* protein-protein binding analysis with eIFs 1, 2 (subunits α-β-γ) and eIF5. Similar to yeast, the eIF3c subunit interacts with eIFs 1 and 5, eIF3a with eIF1 and eIF2β. In addition, the mammalian specific eIF3e binds eIF2β and eIF3l binds eIF1 (Figure S1A, Raw Data Figure S1).

We next focused on eIF3c because it binds two key AUG recognition modulators, eIFs 1 and 5. We showed that the extreme N-terminus of eIF3c binds eIF5, as the only eIF3 subunit, while a less well-defined region between residues 170-240 interacts with eIF1 (Figure 1B, Raw Data Figure 1)^44^. Interestingly, the architecture of the eIF3c-NTD differs between yeast and human proteins with the human protein having the eIF5 and eIF1 binding sites separated by a four α-helix bundle (Figure 1B)^13,44^. Based on this varying positional arrangement, we hypothesized that the putative eIF3c function in proper TIS selection might be conserved but not necessarily identical between yeast and humans. Note that given the rather complex character of the eIF3c–eIF1 interaction (Figure 1B)^43,44^, we further focused primarily on the eIF3c contact with eIF5.

### Knocking down eIF3c eliminates the eIF3-eIF5 interaction and reduces the AUG selection fidelity

To test the eIF3c role in proper TIS selection, we first took advantage of siRNA-mediated knock-downs of individual human eIF3 subunits previously established in our laboratory^34–36^. Knocking down eIF3k concurrently depletes eIF3l in cells but the rest of the complex stays intact; eIF3l^KD^ has the reciprocal effect^34–36^ (Figure S1B); therefore, both KDs were used in subsequent experiments as reciprocal perturbations of the same module. In eIF3e^KD^, protein levels of eIF3e, d, k and l are dramatically decreased, while the remaining eIF3 subunits are largely unchanged^34–36^. This results in the formation of three subcomplexes: eIF3a-b-g-i-c-f-h-m, eIF3f-h-m, and the YLC (eIF3a-b-g-i), which is in the majority (Figure S1B). The eIF3c^KD^ naturally reduces its own levels, in addition to eIF3e, d, k and l, but not the others. As a result, loss of eIF3c completely splits the eIF3a-b-g-i-c-f-h-m subcomplex into the YLC and f-h-m subcomplexes^34–36^ (Figure S1B).

Predictably so, the eIF3c^KD^ and eIF3e^KD^, but not eIF3k^KD^, markedly reduced eIF5-eIF3 association in HeLa cells, as determined by co-immunoprecipitation (Co-IP) analysis using anti-eIF5 antibodies (Figure S1B; Source Data Figure S1). Combined with the *in vitro* binding data (Figure S1A), these results strongly suggest that the eIF3c-NTD is the main and, most likely, only eIF3 anchor for eIF5. Interestingly, based on our ribo-seq data presented in detail below, eIF3c^KD^ and eIF3e^KD^, but not eIF3l^KD^, resulted in significantly increased mRNA and ribosome footprint (FP) levels of eIF5 (Figure S1C, upper panels, Source Data Figure S1, and Raw Data Figure S1). Consistently, increased eIF5 protein levels were seen mainly in eIF3c^KD^ and eIF3e^KD^ (Figure S1C, bottom panels, Source Data Figure S1), which may allow cells to compensate for the overall eIF3 disintegration and the lack of its interaction with eIF5. Noteworthy, FP and mRNA levels of eIF1 were also increased in eIF3e^KD^ but not in eIF3c^KD^ (Figure S1C, upper panels), but we did not pursue this observation further.

To examine the impact of these KDs on fidelity of TIS recognition in human cells (HEK293T and HeLa), we established a new Luciferase-based TIS selection assay (LuTISsa) in combination with siRNA knock-downs. In this assay, the ratio of initiation from near-cognate-initiated to AUG-initiated Luciferase reporters (all set in the strong Kozak context) was calculated individually for each experiment with siRNA KD and compared to control nt (non-targeting siRNA) cells. The eIF3c^KD^ (but not eIF3k^KD^ or eIF3l^KD^) revealed robustly increased initiation on GUG, UUG and AAG near-cognate TISs (Figure 1C, Source Data Figure 1, Figure S1D, and Source Data Figure S1). Interestingly, no significant effect was observed for CUG (Figure 1C), which is the most frequently utilized TIS among all the near-cognate codons and seems to behave similarly to canonical AUG when placed in the strong Kozak context^52–54^, as was the case of our reporters. In agreement, initiation efficiency on AUG *per se* also did not increase. It was even significantly reduced in eIF3c^KD^ (Figure 1C), due to overall downregulation of translation as was confirmed by the large-scale omics analysis presented below. Consistent with our Co-IP experiments, eIF3e^KD^ also increased initiation on near-cognate TISs (except for CUG), although not with statistical significance (Figure S1D). The lack of statistical significance most probably reflects the presence of three subcomplexes in these cells, two of which can associate with 43-48S PICs^34–36^, but only one of them still contains eIF3c (Figure S1B) and therefore its function in TIS selection might be partially preserved. This is not the case for eIF3c^KD^, as neither of the two subcomplexes generated in this knock-down naturally contains eIF3c, and eIF3f-h-m does not interact with PICs^34–36^. In addition, it can also be a consequence of the loss of the eIF3e-eIF2 contact (Figure S1A).

### Three N-terminal residues of eIF3c hold the key to the eIF3c-eIF5 interaction *in vitro* and *in vivo*

Given that eIF3c^KD^ dramatically impacts the eIF3 integrity^34–36^, the observed phenotypes cannot be solely attributed to the loss of the eIF3-eIF5 contact. Therefore, we subjected this contact to a detailed *in vitro* protein-protein binding analysis and revealed that the highly conserved Arg3, Phe4 and Phe5 residues (Figure 1D, Raw Data Figure 1, and Figure S1E) – representing the very N-terminal tip of eIF3c – are solely responsible for its binding to eIF5 (Figure 1D). The individual substitutions of Arg3 with Asn, but mainly of Phe4 and Phe5 with Ala, dramatically impaired the interaction, especially in combination, while the triple mutant R3N F4A F5A eliminated it completely.

To support these observations *in vivo*, we overexpressed N-terminal 180 amino acid (aa) residues of eIF3c (either intact or with the single or triple substitutions) fused to a FLAG tag at the C-terminus in HEK293T cells and immunoprecipitated them using anti-FLAG antibodies. We expected the wild type (wt) but not the mutant N-terminal 180 fragment to compete with the endogenous eIF3 complex for eIF5 (Figure 1E). Accordingly, only R3N displayed residual binding to eIF5, but the other eIF3c-NTD mutants completely failed to pull down eIF5 in contrast to the wt 180 fragment (Figure 1F, Raw Data Figure 1). Reassuringly, the full length eIF3c-FLAG protein carrying the triple mutation pulled down eIF1 but not eIF5 (Figure S1F, Raw Data Figure S1).

Overexpression of eIF5 was shown to relax stringency of AUG recognition and enhance initiation on non-AUG sites^16,39,53^ (see also below). In accord, HEK293T cells overexpressing eIF5 increased the UUG/AUG ratio ∼1.5-fold in our LuTISsa (Figure 1G, compare lanes 1 and 7, Source Data Figure 1, and Raw Data Figure 1). We reasoned that co-overexpression of the intact but not the mutant eIF3c-1-180-FLAG fragment could mitigate this eIF5-imposed increase by sequestering eIF5 excess in a non-productive binary complex, which was observed (Figure 1G, compare lanes 8 - 12 vs. 7). Fittingly, only R3N displayed a residual effect as in previous assays (Figure 1D and F). Collectively, these data strongly indicate that the conserved R3, F4 and F5 residues, especially the latter two, are critically required for the eIF5 association with eIF3 both *in vitro* and *in vivo*.

### Knock-in analysis of the contribution of the eIF3-eIF5 contact to TIS recognition

To investigate the physiological importance and specific role(s) of these residues, we next knocked-in the weaker R3N and stronger F4A and F5A single-point substitutions, as well as the strongest R3N F4A F5A triple substitution into HEK293T cells using the CRISPR-Cas9 system (Figure 2A and Figure S2A; see Methods for details). The real challenge was that the HEK293T cells are hypotriploid, carrying 3 copies of the eIF3c-encoding gene and 3 additional copies of a practically identical pseudogene coding for eIF3cL, both at CH16, and therefore, we were required to knock these substitutions into 6 alleles at the same time. After several rounds, we obtained 2 to 6 homozygous clones for each substitution (with the exception of F4A), the correct instalment of which was carefully verified according to standard field practices (see Methods for details).

**Figure 2.**
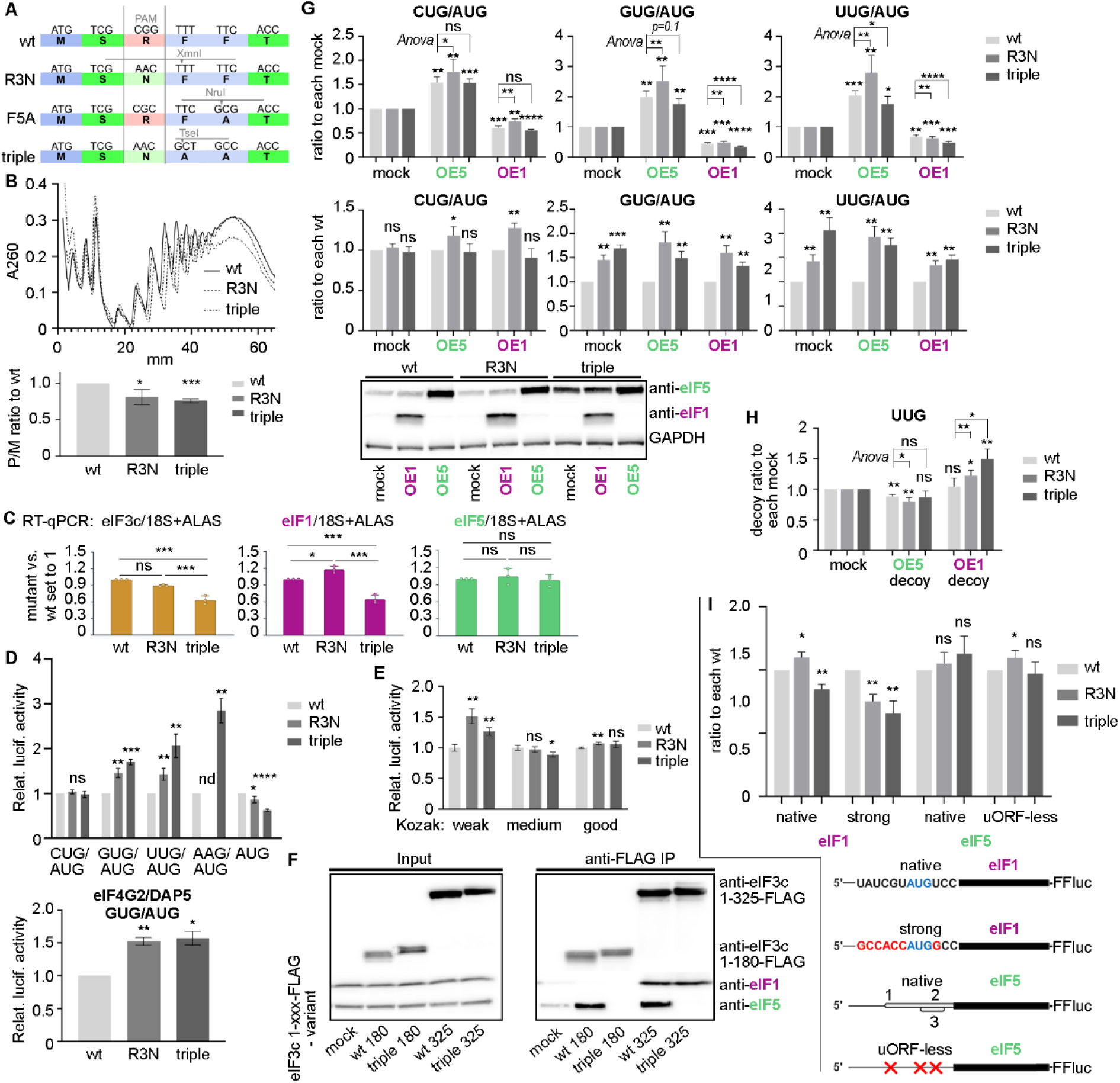
The eIF3c knock-ins impair stringency of TIS selection and disbalance eIF1 and eIF5 protein levels and autoregulation. **(A)** The coding sequences of wt eIF3c-NTD and the R3N, F5A, and triple mutants are shown; the introduced nucleotide substitutions and the corresponding amino acid changes are indicated. (**B)** The eIF3c knock-ins reduce translation initiation rates. Cytoplasmic extracts of wt, R3N, and triple mutant cells were separated by sucrose gradient centrifugation, and polysomal profiles were recorded by measuring absorbance at 260 nm; representative profiles are shown. The lower panel shows quantification of the polysome-to-monosome (P/M) ratio normalized to wt cells. Data are presented as mean ± S.D. (n=4). Statistical significance was determined by One sample t test (* p ≤ 0.05, *** p ≤ 0.001) **(C)** Analysis of endogenous mRNA of *EIF3C*, *EIF1*, and *EIF5* in the eIF3c knock-ins. The mRNA levels of *EIF3C* (golden brown), *EIF1* (magenta) and *EIF5* (sea green) were measured separately using specific primers. For each target gene, mRNA levels were normalized to *18S* rRNA and *ALAS1* reference genes and the quantified qPCR data were plotted (n=3). The differences between experimental groups were tested by ANOVA with post-hoc test. Variables are presented as mean ± S.D. (^∗^ p < 0.05, ^∗∗^ p < 0.01, ^∗∗∗^ p < 0.001). **(D)** The eIF3c knock-ins impair stringency of TIS selection. *Top panel:* wt, R3N and triple cells were transfected with individual firefly luciferase reporters (AUG or CUG, GUG, UUG, AAG); renilla luciferase reporter was used for transfection efficiency normalization. Plot shows the mean ratio ± S.D. of the near-cognate initiation rates relative to those of AUG (n = 4); this ratio in the wt sample was set to 1. Statistical significance was determined by One sample t test (* p ≤ 0.05, ** p ≤ 0.01, *** p ≤ 0.001, **** p ≤ 0.0001). *Lower panel:* The wt, R3N, and triple cells were transfected with luciferase reporters bearing the 5’ UTR sequence of the *eIF4G2/DAP5* gene, whose translation initiates at GUG, which we mutated to AUG in one of the reporters. Renilla luciferase reporter was used for transfection efficiency normalization. Plot shows the mean ratio ± S.D. of normalized GUG/AUG signals (n = 3); the ratio in the wt sample was set to 1. Statistical significance was determined by One sample t test (* p ≤ 0.05, ** p ≤ 0.01). **(E)** The eIF3c knock-ins show different effects on stringency of AUG TIS selection in varying Kozak contexts. The same as in panel (D), except that wt, R3N and triple cells were transfected with firefly luciferase reporters with AUG TIS set in the consensus Kozak sequences of varying strength. The “strong” Kozak sequence was used to normalize for the steady-state initiation rates. Plot shows the mean ratio ± S.D. of normalized “weak”, “medium”, and “good” and signals (n = 4); the ratio in the wt sample was set to 1. Statistical significance was determined by One sample t test (* p ≤ 0.05, ** p ≤ 0.01). **(F)** The eIF3c-NTD segment interacts concurrently with eIF1 and eIF5 *in vivo*. The C-terminally FLAG-tagged N-terminal fragments of eIF3c (1-180 aa, 1-325 aa), either intact (wt) or bearing the triple substitution, were overexpressed in HEK293T cells and immunoprecipitated using anti-FLAG beads. The levels of eIF1, eIF5, and FLAG-tagged eIF3c fragments were detected by western blotting. **(G)** The eIF3c-NTD controls eIF5 activity in pre-initiation complexes. *Top panel:* wt, R3N and triple cells were transfected with AUG- or near-cognate codon-initiated firefly luciferase reporters together with an empty vector (mock) or vectors overexpressing eIF1 or eIF5. Renilla luciferase reporter was used for transfection efficiency normalization. Plots show the mean ratio ± S.D. of normalized near-cognate/AUG signals (n = 4); the mock sample was set to 1 in each dataset. Statistical significance was determined by One sample t test (* p ≤ 0.05, ** p ≤ 0.01, *** p ≤ 0.001, **** p ≤ 0.0001). To determine whether the mutations significantly exacerbate or suppress the overexpression effect, the two-way RM Anova interaction test was performed on the normalized near-cognate/AUG values (*“Anova”* statistics). *Middle panel*: the same as in the top panel, except that each wt sample was set to 1. Plots show the mean ratio ± S.D. of normalized near-cognate/AUG signals (n = 4). Statistical significance was determined by One sample t test (* p ≤ 0.05, ** p ≤ 0.01, *** p ≤ 0.001, **** p ≤ 0.0001). *Lower panel*: eIF1 and eIF5 overexpression was verified by western blotting. **(H)** The impact of artificially altered eIF1 and eIF5 levels on stringency of TIS selection in the eIF3c knock-ins. wt, R3N and triple cells were transfected with UUG-initiated firefly luciferase reporter together with an empty vector (mock) or vectors expressing the eIF3c-NTD fragments: 1-180 aa designated as the “eIF5 decoy” (interacts only with eIF5), and 1-325 aa bearing the triple mutation designated as “eIF1 decoy” (interacts only with eIF1); see panel (F). Renilla luciferase reporter was used for transfection efficiency normalization. Plot shows the mean ratio ± S.D. of normalized decoy/mock ratios (n = 4); the mock sample was set to 1 in each dataset. Statistical significance was determined by One sample t test (* p ≤ 0.05, ** p ≤ 0.01, *** p ≤ 0.001, **** p ≤ 0.0001). To determine whether the mutations significantly exacerbate or suppress the decoy effects, the two-way RM Anova interaction test was performed on the renilla normalized UUG values (*“Anova”* statistics). **(I)** The eIF3c-NTD knock-ins exhibit different effects on the cross-autoregulated expression of eIF1 and eIF5. wt, R3N and triple cells were transfected with two 5’ UTR variants of eIF1 and eIF5 genes placed upstream of firefly luciferase reporters, as depicted at the bottom. Renilla luciferase reporter was used for transfection efficiency normalization. Relative luciferase activities are shown as the ratio to wt cells for each reporter construct. Data are presented as mean ± S.D. (n = 4). Statistical significance was determined by One sample t test (* p ≤ 0.05, ** p ≤ 0.01).

We first tested three clones of each substitution (2 clones obtained in the triple mutant case) for the growth fitness and relaxed TIS selection using LuTISsa (Figure S2B and S2C, Source Data Figure S2). All clones displayed decreased growth rates and increased utilization of GUG and UUG near-cognate TISs compared to wt cells. Interestingly, western blot analyses showed that the R3N clones have generally lower levels of eIF5 but increased levels of eIF1, whereas in F5A clones, the levels of both eIF1 and eIF5 somewhat differ (Figure S2D, Source Data Figure S2, and Raw Data Figure S2). Conversely, both triple substitution clones contain higher levels of eIF5, as in the case of eIF3c^KD^ (Figure S1C), and lower levels of eIF1 (Figure S2D). The eIF3c levels were modestly decreased in the R3N clones. Given the observed consistency across the R3N and triple clones (but not in F5A), varying severity of their phenotypes, and opposite trends in eIF1 and 5 protein levels, from this point on, we continued working with one clone of the R3N and triple substitutions only. In particular, we selected R3N *K10* (hereafter R3N) and R3N F4A F5A *5G8* (“triple”). Collectively, our data suggests that the underlying molecular defect differs in these two knock-ins, most probably due to the partially preserved *versus* lost ability of eIF5 to interact with eIF3. Consequently, cells must employ different compensatory measures to counteract these distinct defects.

Comparing directly these two mutants, they displayed reduced polysomes/monosome ratio, with the triple mutant exhibiting a more pronounced effect (Figure 2B, Source Data Figure 2). Consistently, the cell trace violet (CTV) labeling and forward scatter (FSC) analyses of the cells revealed a significantly lower total protein content and a smaller cell size in the triple mutant (Figure S2E and F, Source Data Figure S2). Given the varying eIF1 and eIF5 protein levels (Figure S2D), we also checked their mRNA levels and found that while the eIF5 mRNA levels in both mutants remained unchanged, the eIF3c and eIF1 mRNA levels were reduced in the triple (Figure 2C, Source Data Figure 2). On the contrary, R3N increased levels of eIF1 mRNA, together suggesting that transcriptional regulation of the eIF1-encoding gene may contribute to the observed expression changes in both mutants to some extent. Co-immunoprecipitation experiments carried out with anti-eIF3a antibodies from formaldehyde crosslinked cell lysates showed that selected eIF3 subunits, representing individual modules, efficiently co-precipitated with eIF3a, thereby confirming the intactness of the eIF3 complex in both mutants (Figure S2G, Raw Data Figure S2).

Similar to eIF3c^KD^, LuTISsa measurements in both mutants revealed no change in initiation frequency at CUG but a significant increase at GUG, UUG, and AAG (Figure 2D, upper panel, Source Data Figure 2). This increase was more pronounced for the triple knock-in. Notably, whereas R3N reduced the AUG selection efficiency only mildly, the triple reduced it by ∼40%, which further magnifies its impact on relaxed TIS recognition. Interestingly, the DAP5/eIF4G2 translon (translon is a general term for translated regions^55^) is known to start from a near-cognate GUG^56^. We therefore went on to test the expression of this translon fused to the luciferase reporter in our eIF3c mutants, replacing the “promiscuous” GUG with canonical AUG. Both mutants significantly increased the GUG/AUG ratio compared to wt cells (Figure 2D, bottom panel), further supporting the general mechanistic impact of the eIF3c knock-in mutants.

Next, we tested the impact of these knock-ins in different Kozak contexts. To do that, the luciferase activities obtained for individual contexts (weak, medium, and good) in each cell line were normalized to the value of the “strong” Kozak context and expressed as a ratio of wt set to 1. We found that both R3N and to a lesser extent the triple mutant displayed significantly increased AUG selection in the weak context (Figure 2E, Source Data Figure 2), supporting the idea of increased dependence of the weak Kozak context on eIF5 levels^53^. Note that these findings were confirmed by the large-scale omics analysis presented below.

### The eIF3c-NTD segment interacts concurrently with eIF1 and eIF5 *in vivo*

To test if eIF1 and eIF5 binding to eIF3c occurs simultaneously and if the loss of eIF5 binding in the triple mutant affects binding of eIF1, we performed the Co-IP assay with 1-180 and 1-325 aa eIF3c constructs (only the latter contains the eIF1 binding site) either intact or with the triple mutation (Figure 2F). Whereas the wt 180 construct pulled down only eIF5, the wt 1-325 construct co-purified with both. As predicted, the triple mutant 1-325 construct completely lost its affinity for eIF5 but displayed no concomitant change in eIF1 binding, supporting the idea that the eIF3c-NTD interacts with eIF1 and eIF5 independently of each other, at least in our experimental set-up. We recognize, however, that based on structural studies^12,13^, this might not be the case in the PIC, since the conformational and structural architecture of the 48S PIC highly likely imposes specific constraints on the eIF3c-NTD availability for the concurrent binding of eIF1 and eIF5 during the AUG recognition process.

### The eIF3c-NTD controls eIF5 activity in pre-initiation complexes

Genetic and biochemical studies suggest that eIF1 discriminates ideal from suboptimal TISs; increased eIF1 levels suppress usage of non-AUG TISs (i.e. increase stringency), whereas decreased eIF1 levels or activity enhance their usage^15,57–60^. eIF5 acts in a opposite way; increased eIF5 levels enhance initiation on non-AUG sites (i.e. decrease stringency), perhaps by increasing the rate of eIF1 release from initiation complexes^16,39,40,53,54,61^. It has been proposed that this concentration dependent interplay could reflect dynamic association and disassociation cycles of eIF1 and eIF5 within PICs to drive TIS selection. Therefore, we examined the effect of both knock-ins with individually overexpressed eIF1 (OE1) and eIF5 (OE5) on TIS selection using LuTISsa (Figure 2G; efficient OEs are displayed in the bottom panel, Source Data Figure 2, and Raw Data Figure 2). As expected, OE5 increased near-cognate initiation at all three tested TISs compared to “mock” (no overexpression) in all tested cell lines, whereas OE1 markedly decreased it (Figure 2G, upper panels). Consistent with Figure 2D, both knock-ins significantly increased initiation on GUG and UUG codons but not on CUG even in OE5 and OE1 conditions (Figure 2G, middle panels).

Interestingly, both OEs mitigated the typically greater impact of the triple mutant on TIS selection (compare Figure 2D and Figure 2G, lower panels), prompting us to investigate whether these knock-ins exacerbate or suppress OE effects. We used the two-way *Anova* interaction test and revealed that R3N exacerbates the effect of OE5 on all tested codons, while the triple partially suppresses the OE5 effect on UUG and GUG and is unresponsive on CUG (Figure 2G, upper panels – *Anova*). These differences can be explained by the triple’s already high eIF5 protein levels; therefore, a further increase brought about by OE5 does not have the additional effect. On the contrary, the triple can be suppressed by OE1 on GUG and UUG (i.e. stringency increases), while R3N (having high endogenous levels of eIF1) is largely irresponsive. These results further underscore the mechanistic differences between these two knock-ins, emanating from “no *versus* residual” eIF5 binding pattern (Figure 1D and F) combined with the opposite protein levels of eIF1 and eIF5 (Figure S2D). In other words, although the absolute levels of near-cognate TIS recognition are expectedly elevated in both knock-ins following OE5 (Figure 2G, upper panels), differing steady-state levels of eIF5 and eIF1 mask the expected effect of a further increase in eIF1 and eIF5 levels via OE in the individual mutants.

To examine this in detail, we introduced an additional imbalance in the levels of the endogenously altered eIF1 and eIF5 proteins in the R3N and triple mutant cells. We did this by co-expressing the UUG luciferase reporter together with eIF3c N-terminal fragments that have different binding capacities for eIF1 and eIF5, as shown in Figure 2F. We then monitored the UUG initiation efficiency of the mutant clones compared to wt (Figure 2H, Source Data Figure 2, Figure S2H, and Raw Data Figure S2). Co-expression of eIF3c-1-180-FLAG fragment (the “eIF5 decoy” that sequesters only eIF5) expectedly decreased UUG initiation in both clones (Figure 2H); however, the magnitude of the induced decrease relative to the mock condition was significantly greater in R3N (Figure 2H – *Anova*), which most probably reflects low eIF5 protein levels in R3N (Figure S2D). Conversely, both knock-ins increased the UUG initiation rate when the eIF3c-1-325-triple-FLAG fragment (the “eIF1 decoy” that sequesters only eIF1) was co-expressed (Figure 2H), however, this time the effect of the triple was greater (Figure 2H – *Anova*). These results clearly indicate that lower endogenous protein levels of eIF5 make the R3N cells more sensitive to eIF5 depletion, while higher endogenous eIF1 protein levels cause the triple cells to respond more strongly to eIF1 depletion. These results further support the masking of predicted effects by differing steady-state levels of eIF5 and eIF1, as observed in Figure 2G.

It is known that both eIF1 and eIF5 mutually maintain their levels in delicate balance through intricate cross-autoregulatory loops. This control involves the 5’ UTR of eIF5 with three short upstream translons (uTranslons^55^ -formerly uORFs) with weak Kozak AUGs on one side, and the eIF1 translon starting with a weak Kozak AUG on the other^53,54^. Increased eIF5 levels stimulate translation of its uTranslons, which in turn decreases eIF5 translation, while increasing eIF1 expression from its weak Kozak AUG. Elevated levels of eIF1 decrease its own translation as well as translation of eIF5 uTranslons, thereby stimulating translation of the eIF5 translon *per se*. To test the impact of our knock-ins on this regulatory circuit, we inserted both 5‘ UTRs in front of the luciferase reporter gene and measured the activity (Figure 2I, Source Data Figure 2). The R3N mutant significantly increased AUG selection in the native (weak) eIF1 AUG context, while the triple decreased it (Figure 2I). However, when AUG was placed in a strong context, both mutants reduced its recognition, in accord with Figure 2D. These results, together with increased eIF1 mRNA levels in R3N but decreased levels in the triple (Figure 2C), may explain differences in the eIF1 protein levels between the two mutants, through which they attempt to compensate for the weakened interaction between eIF3c and eIF5 in opposite ways. Since we did not observe similarly consistent results for the native 5’ UTR of eIF5, we are currently unable to explain the compensatory mechanism that takes place via altered eIF5 expression. In any case, taking altogether, it is apparent that both knock-ins impair the TIS selection process in different ways, resulting in different compensatory strategies.

### The inability of eIF5 to interact with eIF3 diminishes the eIF5 and eIF2 presence in the 43-48S PICs *in vivo*

To understand the underlying molecular mechanism of the observed phenotypes in either knock-in, we assayed the PIC assembly in both mutants using our well-established protocol, where samples are progressively cross-linked as they move through a 7-30% sucrose gradient during high-velocity centrifugation^36^. While we observed no dramatic changes in PIC composition in R3N (Figure 3A, Source Data Figure 3, Raw Data Figure 3), the triple mutant displayed increased eIF1A occupancy, along with significantly decreased occupancy of eIF2 and, in particular, eIF5. This may indicate either unstable binding of eIF5 and eIF2 to PICs or their premature release from PICs. Unlike eIF1 and eIF5, total protein levels of eIF2 and eIF1A are not changed in the eIF3c mutants (data not shown). We were unable to detect eIF1 because none of the eIF1 antibodies we tested worked with the gradient method for unknown reasons. In any case, given the similar effects of both knock-ins on near-cognate TIS selection (Figure 2D), these differing results clearly indicate that merely weakening the eIF3c–eIF5 interaction (as is R3N), rather than completely eliminating it (as in the triple), is sufficient to relax TIS selection stringency. Considering all the differences between both knock-ins, we concluded that the molecular mechanism underlying the R3N defect seems to be more complex and, therefore, more difficult to study. Therefore, we continued with the triple mutant only from this point on.

**Figure 3.**
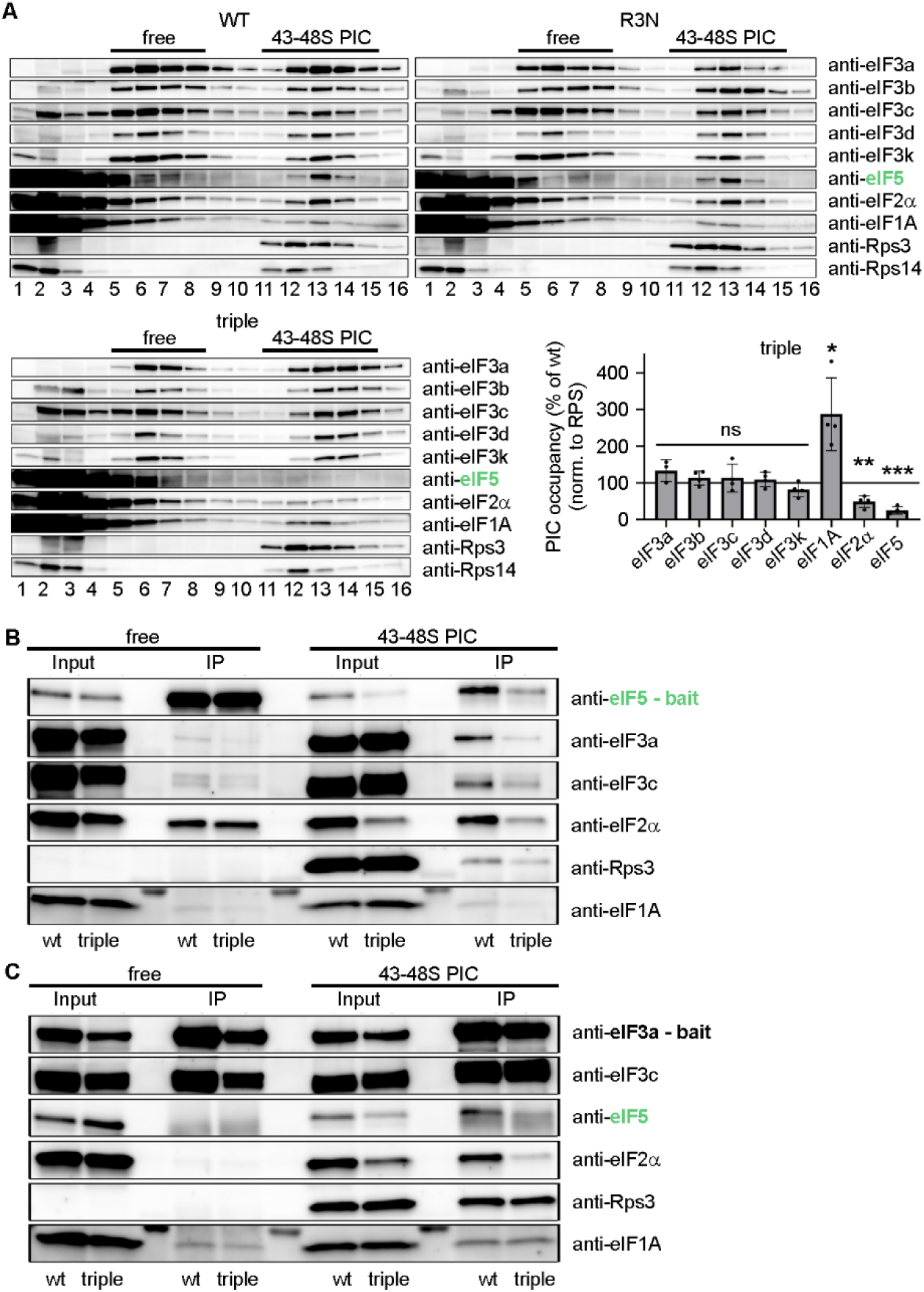
The inability of eIF5 to interact with eIF3 diminishes the eIF5 and eIF2 presence in the 43-48S PICs *in vivo*. **(A)** The 43S-48S PIC assembly analysis by sucrose gradient fractionation in wt, R3N and triple mutant cells. Fraction 1 is a pool of the first two fractions from the top of the gradient. Fractions containing ribosome free versus bound eIFs complexes are indicated at the top. Sucrose gradient fraction numbers are given at the bottom. Plot shows mean ± S.D. of relative amounts of individual eIF3 subunits and other eIFs bound per ribosome (normalized to the Rps14 or Rps3 amount) compared to wt cells. Dots represent results from individual biological replicates (n = 4). One sample t test was used for statistical evaluation (* p ≤ 0.05, ** p ≤ 0.01, *** p ≤ 0.001, ns = not significant). For quantifications, the Quantity One software from Bio-Rad was used. **(B)** Co-immunoprecipitation of eIF5 containing PICs using anti-eIF5 antibodies as a bait. Co-IP was performed separately in the eIF3 ribosome free fractions and in the 43S-48S PICs obtained from the sucrose gradients as in panel (A). Input (4.4% of immunoprecipitated fractions) and IP (25% of eluate) were resolved by SDS-PAGE and analyzed by western blotting using the antibodies indicated on the right. **(C)** Same as in panel (B), except that anti-eIF3a antibodies were used.

To examine the composition of PICs in the triple mutant more rigorously, we pooled 40S-containing gradient fractions, as well as ribosome-free fractions, and carried out Co-IP experiments using either eIF5 or eIF3a as separate baits. As expected, anti-eIF5 antibodies pulled down only eIF2α from the ribosome-free fractions (Figure 3B, Source Data Figure 3, Raw Data Figure 3), further suggesting that the ribosome-free eIF5-eIF3 complex either does not form or is highly unstable, despite the fact that eIF5 can form a stable minicomplex with the eIF3c-NTD fragments *in vivo* (Figs. 1F and 2F). Note that the existence of the eIF2-eIF5 binary complex was demonstrated before^62,63^.

Importantly, pulling down eIF5-containing complexes from a mixture of PICs (scanning and initiating) retrieved only small amounts of eIF3 and eIF2. This strongly supports our earlier single molecule findings that, as the GTP hydrolysis catalyst, eIF5 associates only with initiating PICs and only transiently^24^, therefore only a minority of existing PICs contains eIF5. Importantly, the triple mutant contained reduced amount of eIF5-containing PICs, which is consistent with our gradient analysis (Figure 3A). Pulling down eIF3-containing complexes from free fractions confirmed that neither eIF5 nor eIF2 binds stably to eIF3 outside of PICs (Figure 3C, Source Data Figure 3, Raw Data Figure 3). IP-ing eIF3-containing complexes from the mixture of PICs then clearly documented that most PICs in the triple mutant lack eIF2 and eIF5 factors (Figure 3C).

### eIF5 binding to initiation complexes is slower and less stable in the eIF3c triple mutant

To further probe the molecular defect underlying the triple mutant, we used a recently established real-time, single-molecule FRET assay that directly monitors eIF5 dynamics in a purified, reconstituted human translation initiation system^24^. The assay exploits the proximity of eIF5 to Met-tRNA_i_^Met^ when bound to the initiation complex: a donor dye (Cy3, green) is attached to U46 of Met-tRNA_i_^Met^ and an acceptor dye (Cy5.5, purple) to the N-terminal domain of eIF5, so that eIF5 binding produces a high FRET signal. This approach was previously used to show how eIF5 dynamics fine-tune the fidelity of start site selection during translation initiation^24^.

Here, we used this assay to test whether disrupting the eIF5–eIF3 interaction alters eIF5 dynamics during translation initiation. We assembled 43S PICs (40S subunits, eIF2-TC-Cy3 (green), eIF1, eIF1A, eIF3, and eIF5-Cy5.5 (purple)) containing either wt eIF3 or the eIF3c triple mutant and delivered the 43S PIC, together with eIF5B-Cy3.5 (gold) and 60S-Cy5 (red), to eIF4F-bound mRNA immobilized on the imaging surface (Figure 4A - B). Translation initiation was monitored in real time on mRNAs containing either an AUG or CUG TIS.

**Figure 4.**
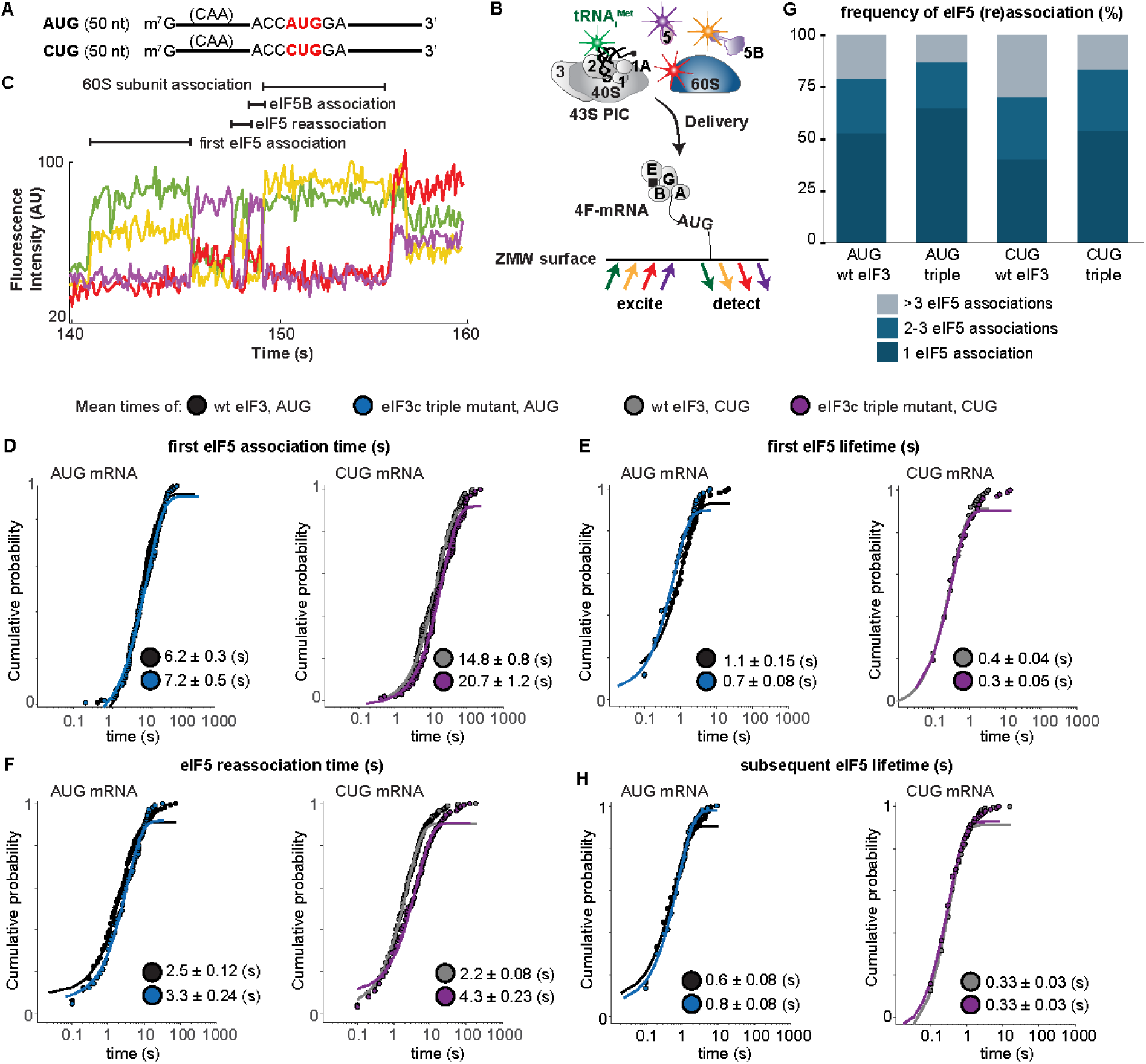
eIF5 binding to initiation complexes is slower and less stable in the eIF3c triple mutant. **(A)** Schematic of model the unstructured CAA repeat mRNAs used in the single-molecule assays. The mRNAs contain either an AUG or CUG TIS in Kozak context. **(B)** Experimental schematic for the real-time single-molecule human translation initiation assay; adopted from^24^. **(C)** Example single-molecule fluorescence data where tRNAi-Cy3 (green), eIF5-Cy5.5 (purple), eIF5B-Cy3.5 (gold), 60S-Cy5 (red) subunits were monitored during translation initiation on the AUG and CUG mRNAs. Molecular association times are annotated above the fluorescence trace; adopted from^24^. (**D**, **E**, **F**, **H**) Cumulative probability plots of the indicated parameters defined in panel (C). Lines represent fit to exponential functions, which were used to derive rates, residency times (lifetimes), and mean times (annotated on plots) for eIF5 with wt eIF3 and AUG mRNA (black), eIF3c triple mutant and AUG mRNA (blue), wt eIF3 and CUG mRNA (gray), and eIF3c triple mutant and CUG mRNA (purple). See Supplementary Excel File 1 for all rates, rate constants, and the number of complexes and binding events analyzed in each experiment. **(G)** Plot of the fraction percentage of translation initiation events that had a single, 2-3, or more than three eIF5 association binding events preceding eIF5B association.

The 43S PIC loading onto the mRNA was marked by a burst of Cy3 fluorescence (Figure 4C). Once the scanning ribosome reached the start site, a burst of eIF5-Cy5.5 fluorescence, correlated with decreased Met-tRNAi^Met^-Cy3 fluorescence intensity, marked eIF5 binding. eIF5 then transiently and repeatedly dissociated and rebound, evident from cycles of FRET loss and reappearance, until a final binding event triggered eIF5B-Cy3.5 association (Cy3.5 fluorescence) and 60S-Cy5 subunit joining (Cy5 fluorescence; Figure 4C).

Compared to wt eIF3, the triple eIF3 mutant delayed the initial association of eIF5 by 1.2-fold on the AUG mRNA and 1.4-fold on the CUG mRNA (Figure 4D, Supplementary Excel File 1), while shortening the initial eIF5 residency time, most notably on the AUG mRNA, where it was reduced 1.6-fold relative to wt eIF3 (Figure 4E, Supplementary Excel File 1). Subsequent eIF5 reassociation of the eIF3c triple mutant initiation complex was also slower and occurred less frequently for both the AUG and CUG mRNAs compared to the wt complex (Figure 4F – 4G, Supplementary Excel File 1). These effects suggest that the triple mutant destabilizes eIF5 interactions with the initiation complex and are consistent with our *ex vivo* observations (Figure 3A).

Interestingly, the residency time of subsequent eIF5 reassociations was longer on the AUG mRNA in the eIF3c triple mutant initiation complex compared to the wt complex, while no differences were observed on CUG mRNA (Figure 4H, Supplementary Excel File 1). One possible explanation is that TIS selection triggers the well-characterized P^out^ to P^in^ conformational transition^10,11^, which may expose additional eIF5-binding surfaces in the PIC that stabilize the interaction or facilitate eIF5 activation of GTP hydrolysis by eIF2, an effect that would be more pronounced at a cognate AUG start site. Consistent with this, the final eIF5 binding event and subsequent association of eIF5B are largely unaffected by the eIF3 triple mutant on the AUG mRNA, whereas both the final eIF5 association and eIF5B association are delayed on the CUG mRNA (Figure S3A – C, Supplementary Excel File 1).

Collectively, these results strongly support a role for eIF3 in promoting stable eIF5 binding to the initiation complex upon reaching AUG or a near-cognate TIS. They also align well with our *ex vivo* gradient data (Figure 3) and with the observation that, while the triple mutant relaxes initiation at UUG, GUG, and other near-cognate TISs, it shows reduced AUG selection efficiency relative to wt (Figure 2D). The absence of a measurable CUG effect in bulk TIS assays (Figs. 1C and 2D), despite clear differences here, likely reflects known differences in sensitivity between bulk and single-molecule measurements. We note that our single-molecule assay conditions favor stringent start codon selection, yielding low initiation frequencies on UUG and GUG mRNAs; this precluded quantitative analysis of near-cognate TISs showing relaxed stringency in the eIF3 mutants (Figs. 1C and 2D).

### Translatome-wide consequences of the relaxed stringency of AUG recognition

To determine what consequences the inability of eIF5 to associate with eIF3 has on the entire translatome, we subjected eIF3c^KD^ and our triple mutant cells to ribosome profiling– –ribo-seq analysis combined with RNA-seq. Generally, both the eIF3c^KD^ and the triple mutant had fewer ribosomes bound to coding regions, suggesting reduced initiation rates, as expected^34^ (Figure 2B). The effect was stronger for the eIF3c^KD^. This chapter highlights the key aspects of these analyses; a complete description, supplemented by Figures S4, S5 and S6, and Supplementary Excel File 2, is available in Supplementary Material.

We first examined differential translation efficiency by calculating translation efficiency (TE) from ribosome footprint (FP) and mRNA sequencing data using DESeq2^64^, where TE reflects ribosome footprint abundance along the entire translon normalized by mRNA levels. As expected, the eIF3c^KD^ exhibited a greater number of significant changes in TE, FP, and mRNA compared with the triple mutant (Figure S6 *vs.* S5, Supplementary Excel File 2). This reflects eIF3c^KD^’s deleterious effect on eIF3 integrity and, consequently, its much stronger impact on the functionality of the translation initiation mechanism.

We focused our TE analysis on the triple mutant, primarily because of its highly distinctive impact on translation initiation, which should allow for more direct interpretation. The triple mutant library contained 557 significant DTEGs (adjusted *p*-value < 0.05), comprising 385 TE-upregulated and 172 TE-downregulated genes. For the top hits and GO/KEGG enrichment analysis please see Figure S5 and Supplementary Results. The most interesting genes were downregulated DTEGs, which showed a strong enrichment in categories related to translation and ribosome (Figure S5C). These terms are consistent with the observed reduced P/M ratio (Figure 2B) and reduced total protein content and cell size (Figure S2E and F). Downregulated DTEGs were also enriched in mitochondrial function, including oxidative phosphorylation (Figure S5C), consistent with a significant overlap with the Human MitoCarta 3.0 database (Supplementary Excel File 2): 29 MitoCarta genes were represented in downregulated DTEGs (*p*-value < 0.01, hypergeometric test), and 7 in upregulated DTEGs (*p*-value > 0.05). They included mtDNA-encoded as well as nuclear-encoded OXPHOS subunits, and mitochondrial ribosomal proteins (Figure S5D).

### Wild-type translation initiation levels anticorrelate with initiation under the eIF3c-NTD impairment

To examine the effect of impaired eIF3c on the efficiency/frequency and stringency of initiation on individual canonical and non-canonical TISs, we derived a measure of translation initiation (TI) for all start codons in the 5’ UTR and for the main translon’s AUG. We calculated TI as the ribosome footprints localized to the P-site at each respective start codon, normalized to the sequencing depth and RNA level. We applied several quality filters to obtain a high confidence set of occupied TISs. In addition, we conducted several tests to confirm that the reads observed on the start codons originated from truly initiating ribosomes rather than from ribosomes idling on the TIS (Figure S7A). Specifically, we confirmed that (i) reads downstream of the TIS were present, (ii) the downstream reads were correlated with TI, and (iii) a downstream read position followed a 3-nucleotide periodicity pattern consistent with a truly elongating ribosome. We found that even under highly impaired conditions, i.e. eIF3c^KD^, ribosome footprints on start sites were initiating.

Using TI, we systematically analyzed the effect of impaired eIF3c on all TISs using TI fold change (TI FC) from the baseline wt TI to the TI in the impaired condition. Ribosome occupied start codons had expected frequencies in both datasets (Figure S7B): CUG, GUG, AUG, and UUG were the most frequently occupied codons, while AUC, AUA, AAG were the least frequently occupied codons. Similarly, median initiation capacity per codon type followed expected patterns (Figure 5A): AUG was the strongest initiator, irrespective of its location ahead of the main translon or upstream in the 5’ UTR; ACG, CUG, GUG, UUG demonstrated medium TI; and AAG and AGG showed the smallest TI in the wt cells. Note that TI in Figure 5A is not a fold-change: it can be negative simply due to the normalization procedure.

**Figure 5.**
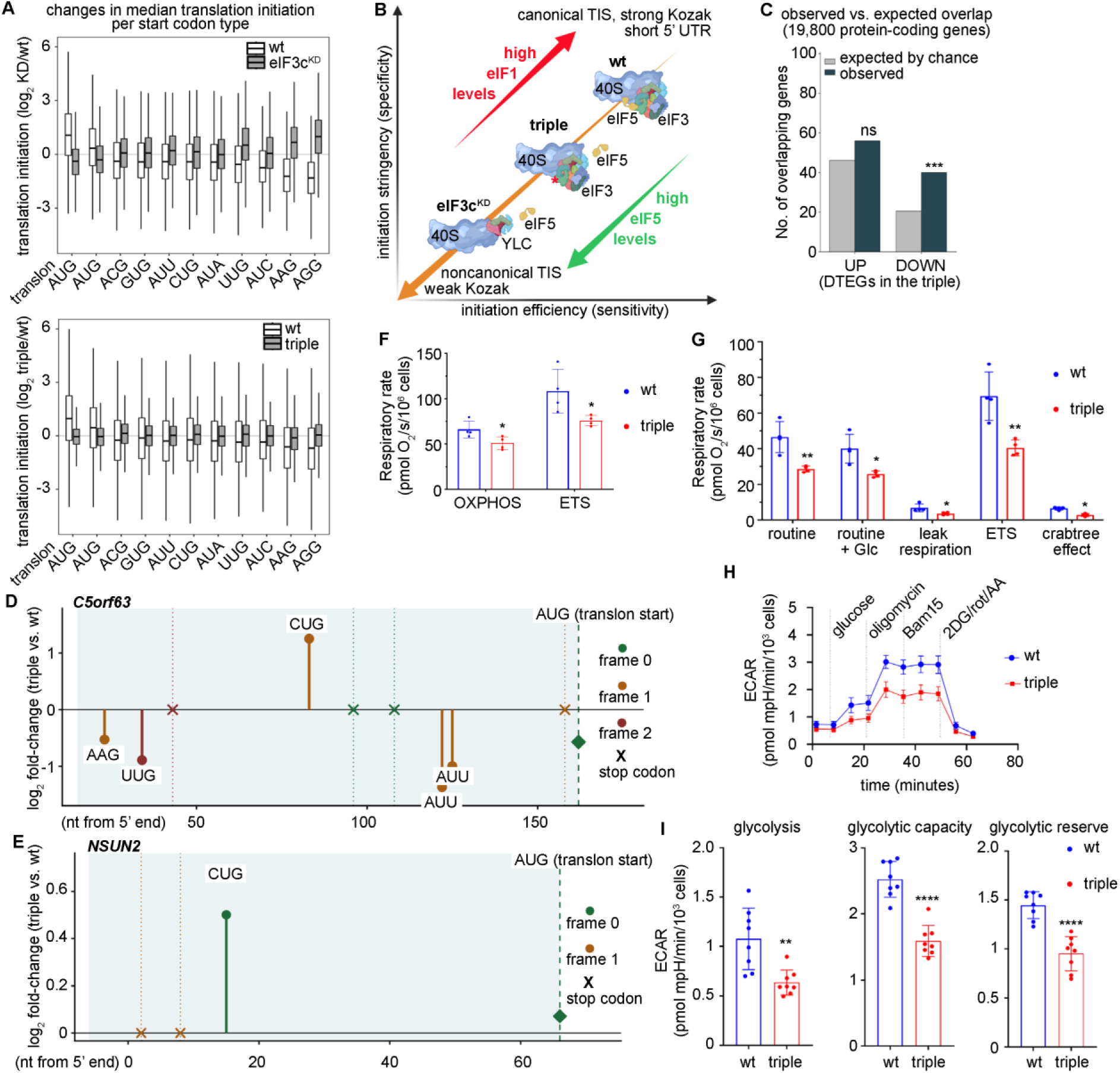
Impaired eIF3c function reshapes TIS selection and compromises mitochondrial and glycolytic activity. **(A)** Changes in median translation initiation per start codon type. Translation initiation (TI) was estimated by normalizing the ribosome footprints with the P-site on the start codon by sequencing depth and RNA levels. The boxplots show the median of log_2_ TI in wt and eIF3c^KD^ (top panel) or wt and the triple mutant (lower panel) with the interquartile ranges for all TIS types in the 5’ UTR, and for the main translon’s AUG. Start codons were ordered from left to right with increasing wt TI. eIF3c^KD^ (KD) and triple mutant are shown in grey, and the corresponding wt in white. **(B)** Model of changes in specificity and sensitivity in upon the eIF3c-NTD impairment. The ribosome is shown in its canonical initiation state, the triple mutant state and the eIF3c^KD^ state. The x-axis denotes sensitivity regarding the overall probability to initiate. The y-axis denotes specificity with respect to requiring canonical start codons or strong Kozak sites. Arrows show the decrease between wt and each impaired state, as well as the impact of increased protein levels of eIF1 and eIF5 on the aspects studied. **(C)** Intersection of genes with alternative N-terminal isoforms (Ly et al.)^7^ with differentially translated genes. Number of overlapping genes observed (dark) *versus* expected by chance (grey), for genes with downregulated (decrease in TE from wt to triple mutant) or upregulated (increase in TE from wt to triple mutant) translation efficiency. Asterisk denotes significance at *p*-value < 0.001 (two-tailed hypergeometric test). **(D)** Diagram of 5’ UTR of *C5orf63* with upstream out-of-frame CUG, 79 nt upstream of the main translon’s AUG, which shows an increase in translation initiation in the triple mutant. Stem height gives the change in initiation, log_2_ (TI) at each alternate start codon (circle) and the change in translation efficiency, log_2_ (TE) at the main translon’s AUG (diamond). Stop codons are denoted by a cross and color denotes reading frames, with frame 0 as the frame of the main translon’s AUG. **(E)** Diagram of 5’ UTR of *NSUN2* with an in-frame CUG, 51 nt upstream of the main translon’s AUG, with increased TI in the triple mutant. Stem height gives the change in initiation, log_2_ (TI) at each alternate start codon (circle) and the change in translation efficiency, log_2_ (TE) at the main translon’s AUG (diamond). Stop codons are denoted by a cross and color denotes reading frames, with frame 0 as the frame of the main translon’s AUG. **(F)** Oxygen consumption measurements in permeabilized wt and triple mutant cells. Respiratory fluxes associated with coupled respiration (OXPHOS), reflecting ATP-generating respiratory capacity, and FCCP-induced electron transport system (ETS) capacity are shown. Data are presented as mean respiratory rate per 10⁶ cells ± S.D.; replicates: wt (n=4), triple mutant (n=4). Statistical significance was tested by unpaired two-tailed *t*-test (* p ≤ 0.05). **(G)** Oxygen consumption measurements in intact wt and triple mutant cells. Respiratory fluxes associated with routine respiration, glucose-stimulated respiration (routine + Glc), leak respiration after oligomycin addition, and FCCP-induced maximal electron transport system (ETS) capacity are shown. The Crabtree effect was calculated as the difference between routine respiration and routine respiration after glucose addition. Data are presented as mean respiratory rate per 10⁶ cells ± S.D.; replicates: wt (n=4), triple mutant (n=4). Statistical significance was tested by unpaired two-tailed *t*-test or nonparametric Mann Whitney U test (* p ≤ 0.05; ** p ≤ 0.01). **(H)** Seahorse XF analysis of extracellular acidification rate (ECAR) in wt and triple mutant cells. Mean ECAR traces (± S.D.) measured in adherent wt and triple mutant cells using Seahorse XF analysis. Curves show dynamic changes in ECAR following sequential substrate and inhibitor injections used to interrogate glycolytic function; biological replicates: wt (n=8), triple mutant (n=8). **(I)** Quantification of glycolytic parameters derived from ECAR measurements. Basal glycolysis (glycolysis under glucose stimulation), glycolytic capacity (maximum ECAR following oligomycin injection), and glycolytic reserve (difference between glycolytic capacity and basal glycolysis) are shown. Data are expressed as ECAR per 10³ cells and presented as mean ± S.D.; biological replicates: wt (n=8), triple mutant (n=8). Statistical significance was tested by unpaired two-tailed *t*-test (** p ≤ 0.01; **** p ≤ 0.0001).

Next, we examined the average fold-change in TI under impaired eIF3c (eIF3c^KD^ or the triple) per codon type, organizing codons in the same order as before, in decreasing order of wt TI (Figure 5A). Overall, the average increase in median TI from wt to eIF3c^KD^ was much larger than the increase from wt to the triple, consistent with what we discussed for TE above. For eIF3c^KD^, but also for the triple, we observed a strong codon dependence of change in: TI decreased most strongly on AUG codons, and changed little for ACG, GUG, CUG, UUG, AUU, and AUA. TI increased the most for AUC, AAG, and AGG; i.e. precisely the codons with the lowest average TI in wt. For the triple, the fold-changes deviated to some extent compared to eIF3c^KD^, most likely due to the overall more subtle effects on eIF5 recruitment discussed above.

Based on these results, we hypothesized that the change in TI upon eIF3c impairment was mostly determined by TI in the wt, regardless of the codon type. Indeed, the fold-change in TI (TI FC) and TI wt are inversely correlated in both eIF3c^KD^ (*R=*0.35; Figure S7C) and to a smaller extent in the triple. This finding suggests that highly utilized start codons, such as AUG, have reached saturation with respect to translation initiation under wt conditions; however, upon eIF3c impairment, the general translation initiation efficiency decreases, thereby initiation at these codons decreases as well (Figure 5B). This is consistent with our LuTISsa reporter data (Figure 2D). Conversely, for codons that are rarely initiated in wt, such as AAG and AGG, initiation increases upon disruption of eIF3c function, likely due to a reduction in PIC stringency (Figure 5B).

Next, we used quantitative modeling to determine the relative contributions of TI and other factors to TI FC under eIF3c impairment. Due to the smaller overall TI FC, the triple mutant was much less predictable than the eIF3c^KD^ but again showed similar qualitative trends. As sequence-based features, we tested the TIS (start codon) type, the strength of the translation context (Kozak score), the GC content around the TIS, the overall GC content of the 5’ UTR, the length of the 5’ UTR and the position of the TIS within the 5’ UTR. The variables were not correlated with each other (*R*<0.80). Figure S8A displays the results as the correlation coefficient between observed TI FC and the predicted (modeled) TI FC for individual and sets of features; Supplementary Excel File 3 contains all input data. The analyses showed that indeed TI wt - which is determined mostly by TIS type and Kozak context - were the features that held the most predictive power.

Overall, this analysis confirmed what we had observed for the LuTISsa reporter constructs for both eIF3c^KD^ and the triple (Figs. 1C and 2D – E). Similarly to the above, we found that in TIS such as AAG or AGG with a low wt TI, or in TISs with suboptimal Kozak, disruption of the eIF3c function resulted in a greater increase in TI than in other cases. It also further supported our triple mutant single molecule analysis (Figure 4), showing that the impairment of the eIF3c–eIF5 contact makes eIF5 to bind to PICs more slowly, depart from it more rapidly, and transiently rebind the PICs less frequently, with the degree of eIF5 perturbation depending on the non-AUG codon identity. In the words of the omics analysis, it leads to a loss of both initiation efficiency and TIS recognition stringency, resulting in reduced initiation at AUG and strong Kozak sites but increased initiation at non-canonical start codons and weak Kozak sites (Figure 5B).

### Genes of mitochondrial localization and function are more affected by eIF3c impairment than other genes

Next, we explored the significant enrichment in mitochondrial genes (specifically, 29 of the total 172 downregulated DTEGs were found in Human MitoCarta3.0; *p*-value < 0.01; Supplementary Excel File 2), particularly within the OXPHOS pathway, which was observed in the triple mutant (Figure S5C - D). We probed for possible mechanisms that affect mitochondrial genes more than other genes, beyond the general balancing effect of decreased PIC sensitivity and specificity of the mutant eIF3c. First, mitochondrial genes tend to be highly translated in wt; i.e. have high wt TI on the AUG of the main translon (Figure S8B). Therefore, following general trends, TI is expected to markedly decrease upon eIF3c impairment. Second, in the triple mutant, we identified examples of mitochondrial genes with increased TI at near-cognate TIS in the 5’ UTR, which were out of frame with the main translon’s TIS. Naturally, initiation at these upstream TISs would reduce downstream translation of the main translon by the canonical uTranslon-mediated repressive mechanism. One such gene is C5ORF63, a Glutaredoxin-Like Protein, which contains CUG in its 5’ UTR in frame 1 (Figure 5D). This CUG exhibits greatly increased ribosome occupancy (increased TI) in the triple mutant at the expense of decreased ribosome occupancy at the canonical AUG start, whereas no change or reduction occurs at any of the other upstream TISs. This clearly suggests that initiation at the CUG represses downstream translation of the main translon. Third, many mammalian nuclear-encoded mitochondrial mRNAs have very short 5′ UTRs and a specialized start-site motif called TISU. Unlike conventional initiation, TISU-dependent translation is stimulated by eIF1^65^. The decrease in eIF1 levels in the triple mutant (Figure S2D and Figure 5B) renders decreased translation of TISU-containing genes highly feasible.

Importantly, we suggest another, previously overlooked route how erroneous initiation in the triple mutant may affect mitochondrial genes, i.e. through the production of protein isoforms with different N-termini. Although the origin of these isoforms is poorly explored, it is known that they often have alternative localization within the cell and include in-frame N-terminal extensions or truncations^7^. Indeed, we found a significant overlap between the downregulated DTEGs and a published dataset of proteins with different N-termini^7^ (p-value < 0.05, hypergeometric test) (Figure 5C and Figure S8C). Strikingly, we also detected a significant overlap between the dataset of proteins with alternative N-termini^7^ and annotated mitochondrially-imported genes as defined in MitoCarta 3 (p-value < 0.001, hypergeometric test). From the latter set, we identified genes with a reported N-terminal extension, which have in-frame upstream near-cognate TIS(s) but no in-frame stop codon downstream, and whose TI changed in the triple. The list of these genes, which are candidates for true isoform switching, leading to the mislocalization of the mitochondrially-imported proteins but not necessarily to the altered TE, is shown in Supplementary Excel File 3. A prime example is NSUN2, an RNA methyltransferase, which is partially localized to mitochondria^66^. Its 5’ UTR has an in-frame CUG at position 15 that exhibits increased TI in the triple (Figure 5E), suggesting the synthesis of NSUN2 N-terminal extension. Aberrant translation from this CUG would interfere with proper function of the MLS of the canonical NSUN2 isoform.

Collectively, our results suggest that, in addition to general changes in TE of some mitochondrial genes, they can also be affected in terms of translation of the correct protein isoform with a proper targeting signal, which may or may not alter their overall expression levels. These findings lead us to propose that mitochondrial genes, along with essentially any other genes encoding proteins with N-terminal targeting signal domains, constitute a group of one of the most vulnerable genes, whose incorrect translation initiation adversely affects many cellular processes. In support, we present two examples of the N-terminal switching of non-mitochondrial genes in PPP1CB and CITED2^7^. PPP1CB encodes a serine/threonine-protein phosphatase PP1-beta catalytic subunit^67^ and is upregulated in our triple dataset. It has an in-frame CUG at position 27 nt upstream of the AUG with increased TI and no in-frame stops (Figure S8D). It is cytoplasmic/nuclear and its extended isoform is predicted to maintain the same localization^7^. CITED2, encoding Cbp/p300-interacting transactivator 2^68^, has the in-frame AUA 183 nt upstream of AUG with no stops in-between, which exhibited very high TI (Figure S8E), indicating synthesis of its N-terminal isoform. CITED2 is cytoplasmic/nuclear too, but its extension is predicted to shift localization^7^.

### The inability of eIF5 to interact with eIF3 compromises mitochondrial function

The fact that the loss of binding between eIF3c-NTD and eIF5 decreased expression of mtDNA-encoded, nuclear-encoded OXPHOS subunits, together with structural subunits of the mitochondrial ribosome (Figure S5D, Supplementary Excel File 2), strongly suggests reduced mitochondrial respiration. Conversely, upregulation of PDPR, CA5B, and GPD2, which we also observed (Figure S5D, Supplementary Excel File 2), indicates metabolic rewiring through enhanced pyruvate utilization. Increased expression of MIEF1, MTX3, SLC30A9, and AHCYL1 then points to the adaptive regulation of mitochondrial dynamics and homeostasis. Collectively, these changes indicate a complex compensatory response to mitochondrial dysfunction.

To study mitochondrial function in the triple mutant cells, we measured oxygen consumption using a high-resolution respirometry system (Oroboros). Respiration was assessed in the permeabilized cells (digitonin-treated, substrate-saturated conditions) as well as in intact cells. In the permeabilized cells, the triple mutant exhibited significantly reduced oxygen consumption in the coupled, i.e. phosphorylating state (OXPHOS), as well as when electron transport system (ETS) capacity was assessed after FCCP addition, when compared to wt cells (Figure 5F and Source Data Figure 5). Similarly, measurements in intact cells (Figure 5G and Source Data Figure 5) revealed significantly reduced routine respiration, respiration after addition of glucose, oligomycin-insensitive respiration (leak respiration), and FCCP-induced maximal ETS capacity. Together, these results demonstrate limited mitochondrial respiratory function in the triple mutant cells. Interestingly, the Crabtree effect (drop in oxygen consumption after glucose addition), expectedly observed in wt cells, was significantly less pronounced in the triple mutant (Figure 5G). Lack of Crabtree effect clearly indicates impaired coupling between glycolytic and mitochondrial respiration fluxes.

Next, we performed Seahorse XF analysis, which enables simultaneous measurement of oxygen consumption rate (OCR) and extracellular acidification rate (ECAR) in adherent cells. As shown in Figures S9A – B (Source Data Figure S9), the OCR analysis yielded data similar to those obtained from measurements using the oxygraph Oroboros. Compared to wt cells, the triple mutant cells exhibited reduced basal (routine) respiration, basal respiration after glucose addition, leak respiration, and maximal respiration, indicating a diminished respiratory reserve. In addition, the Crabtree effect was again significantly less prominent in cells with a triple mutation (Figure S9B). Interestingly, ATP-linked respiration remained comparable. Together, these data suggest that mitochondrial ATP production is largely preserved under physiological non-stress conditions, whereas the capacity to increase respiration under conditions of elevated energy demand is compromised. In addition, we measured the ECAR parameter, which serves as an indicator of the glycolytic rate. Strikingly, this analysis revealed a marked reduction in glycolysis, glycolytic capacity, and glycolytic reserve in triple mutant cells (Figure 5H – I and Source Data Figure 5). These data suggest that glycolytic function is suppressed in the triple mutant cells, which is in line with the decreased Crabtree effect (Figure 5G and Figure S9B).

Taken altogether, these results provide the first direct causal link between stringent TIS recognition and proper mitochondrial respiration and glycolytic activity, implicating the eIF3–eIF5 contact in maintaining metabolic homeostasis.

## DISCUSSION

To elucidate precise molecular contribution of genes encoding eIFs and ribosomal proteins to translational control requires targeted *in cellulo* mutagenesis. However, the predominant (if not exclusive) approach to date has been based on CRISPR-Cas- or siRNA-mediated knockout or knockdown, or the use of a degron, such as dTAG or auxin-inducible (AID) (see for example^34–36,69–73^). These approaches have several limitations, the most problematic of which are: i) the essentiality of most genes associated with translation, ii) the relatively long time required to effectively deplete the factor under study, and iii) the concomitant degradation of related but untargeted factors. To circumvent all these constraints, we first identified key eIF3c residues for its binding to eIF5 and, using CRISPR-Cas, introduced specific point mutations in the extreme N-terminus of the eIF3c subunit in HEK293T cells without disturbing the eIF3 integrity. We then analyzed how disrupting eIF3 binding to eIF5 affects the stringency and efficiency of AUG recognition at the mechanistic, translatome-wide, and single gene levels.

Mechanistically, we have demonstrated that the conserved R3, F4 and F5 residues of eIF3c, particularly the latter two, are critically required for eIF5 binding to eIF3 both *in vitro* and *in vivo* (Figure 1 and Figure S1). Our LuTISsa reporter assay revealed that disturbing the eIF3c–eIF5 contact *in vivo* results in less efficient AUG selection compared to wt, despite otherwise more relaxed initiation at UUG, GUG and other near-cognates TISs, or at AUG in the weak Kozak context (Figure 2D – E). These observations align very well with our bioinformatics analysis showing that decreased PIC sensitivity and specificity due to eIF3c dysfunction causes decreased bulk initiation at the canonical AUG TIS but increased initiation at non-canonical TISs (summarized in Figure 5A - b). Therefore, we propose that impairment of the eIF3–eIF5 contact interferes with the proper activation of GTP hydrolysis on eIF2-TC by eIF5 in such a way that the rate of initiation at the AUG TIS is reduced, while at the same time the discrimination against near-cognate TIS is mitigated, allowing for increased initiation.

Strikingly, whereas the R3N knock-in merely weakens the eIF3–eIF5 contact, the triple mutant eliminates it entirely, and only the complete loss of binding greatly impairs the stable association of eIF5 with the 48S PIC that has just reached the AUG or near-cognate TIS (Figs. 3 and 4). However, since both knock-ins impair the stringency of TIS selection to a similar extent (Figure 2), it is apparent that the underlying molecular mechanisms differ. Indeed, both mutants disturb the balance between eIF1 and eIF5 protein levels in opposite ways (Figure S2D), clearly suggesting that cells also employ different compensatory measures to counteract these distinct defects at least partly through the cross-autoregulation of eIF1 and eIF5 protein expression^53,54^ (Figure 2I). While we did not examine the effect of R3N in great detail due its apparent complexity and a less pronounced phenotype, our *ex vivo* and single molecule experiments carried out with the triple mutant revealed that the eIF3c-eIF5 interaction has distinct roles at different stages of eIF5 binding to PICs (Figs. 3 and 4).

It is well-known that the C-terminal domain (CTD) of eIF5 is a central hub for interactions with PIC components, namely eIF1, eIF1A-NTT, eIF3c-NTT and eIF2β-NTT^74–78^. The interaction surfaces of these factors on the eIF5-CTD overlap, suggesting rapid dynamics of mutually exclusive binding events during the eIF5 presence in the PIC^79^. This is corroborated by the fact that eIF5-CTD has not yet been captured in any of the available PIC structures (the sole exception is our structure of the specific *T. cruzi* PIC^44^; however, we did not detect any binding between eIF3c and eIF5 recombinant proteins in *T. cruzi*). Therefore, the sequence of the eIF5-CTD-mediated interactions likely reflects the well-characterized conformational changes in the PIC upon TIS selection. Our findings that the initial binding of eIF5 to PIC is delayed, that the first association lifetime is shorter, and that the subsequent eIF5 reassociation times are also slower and occur less frequently compared to wt (Figure 4), clearly implicate eIF3c in the recruitment of eIF5 to the PIC. The fact that the residency time of subsequent eIF5 reassociations is prolonged then suggests that the conformational transition of the PIC from the P^out^ to P^in^ states in the absence of the eIF3c-eIF5 contact is disturbed, highly likely due to impaired eIF5-CTD interactions with its other binding partners. Finally, the fact that the final eIF5 lifetime and eIF5B association are delayed on the CUG mRNA supports our theory that eIF3c controls the eIF5 activation of GTP hydrolysis on eIF2 and their subsequent release from the PIC. In addition, it may also explain why the triple mutant alleviates discrimination against near-cognate TIS (Figs. 2 and 5).

Consistent with the proposed eIF3c control over activation of GTP hydrolysis, we observed depletion of eIF5 and eIF2 proteins from the 43S-48S PICs-containing fractions in the triple mutant, while there was a concurrent increase in eIF1A levels (Figure 3). The departure of eIF2 from the PIC and the stabilization of eIF1A binding on the PIC upon TIS recognition are known hallmarks of the closed/scanning-arrested state of the PIC^80^. This stabilization is most likely mediated by the eIF1A-NTT, which becomes structured in the P site of the closed PIC, and is thus unavailable for interaction with eIF5-CTD, unlike in the open state^12^. We therefore propose that the triple mutant allows premature GTP hydrolysis on eIF2 and temporarily locks the PIC in the closed/P^in^ state. In support, it is known that disruption of the eIF5-CTD-eIF2β contact disturbs the closed PIC conformation and increases initiation stringency^77,81^. Since the eIF1A-NTT, eIF3c and eIF2β-NTT interaction sites on the eIF5-CTD overlap, it is possible that elimination of the eIF3c–eIF5-CTD interaction during TIS recognition shifts the interaction equilibrium among these proteins towards the eIF5-CTD-eIF2β contact. This may accelerate GTP hydrolysis and thus the adoption of the closed/P^in^ state, as well as the subsequent ejection of eIF2 and eIF5 form the PIC, thereby deregulating the initiation stringency.

Strikingly, however, it seems obvious that in order to maintain sustainable physiological levels of translation in the triple mutant, a natural compensatory strategy for insufficient eIF5 recruitment to the PIC would be to reduce eIF1 levels and increase eIF5 levels, all at the expense of deregulated initiation stringency due to an impaired progression of sequential eIF5-CTD interactions with its binding partners during TIS selection, as we observed (Figure 2 and Figure S2). It is plausible that this compensatory mechanism results systemically in decreased AUG recognition in the triple mutant (Figure 2D). In summary, our data strongly support the observation that human eIF5 associates with the PIC as late as during the actual selection of a cognate or near-cognate TIS^24^, and that its stable binding and proper functioning during TIS selection strictly require attachment to eIF3. The latter conclusion is well supported by the latest structural analysis showing the eIF3c-NTD occupancy at the 40S interface side even after the arrival of eIF5^12^.

At the translatome-wide level, we have become most interested in 36 genes (out of total 557 DTEGs) with altered translational efficiency in the triple mutant that were associated with mitochondrial function (Figure S5). Given this significant number of hits, we proposed several possible mechanisms affecting translation of mitochondrial genes. Next, we examined mitochondrial fitness of the triple knock-in cells and found reduced oxygen consumption, affecting respiratory fluxes associated with OXPHOS and ETS capacity (Figure 5F – i and Figure S9). In addition, we also detected impaired coupling between glycolytic activation and mitochondrial respiratory regulation. Indeed, the triple mutant cells exhibited a marked reduction in glycolysis, glycolytic capacity, and glycolytic reserve (Figure 5H - I). Together, these data indicate a coordinated suppression of both mitochondrial respiration and glycolytic function in the triple mutant. Noteworthy, the expression of known markers of the mitochondrial unfolded protein response (UPR^mt^), which is essential for protecting mitochondria from proteotoxic damage^82^, namely HSPD1, HSPA9, HSPE1, LONP1, remained unchanged. It is therefore unlikely that the triple mutant cells undergo a severe UPR^mt^ response.

How might defective TIS selection in the cytoplasm affect mitochondrial function? In other words, what is this link? Ly et al. analyzed existing N-terminal protein isoforms and identified hundreds of differentially localized isoform pairs, including dual-localized isoforms, which are essential for both mitochondrial and host cell function^7^. As we have identified significant overlaps between these isoforms and our downregulated DTEGs, as well as between these isoforms and annotated mitochondrially imported genes, we propose that – in addition to the overall dysregulation of translation efficiency of many mitochondrial genes – the imbalance in production of the correct mitoprotein isoforms represents another important factor contributing to mitochondrial dysfunction observed in the triple mutant. Mistranslated isoforms will most likely fail to reach their intended destination due to a missing or incomplete MLS, which always occurs at the N terminus of mitoproteins^83^. Note that between 1,000 – 1,500 (MitoCarta3.0 lists 1,136 human genes) nuclear-encoded proteins are targeted to the mitochondria through various mechanisms, which requires a complex import, processing, and assembly system^84^. Therefore, even a relatively minor disruption to this system can have a fairly dramatic impact on mitochondrial fitness.

When viewed from an even broader perspective, the function of virtually any protein with two or more N-terminal isoforms will be affected by improper translation initiation. Some isoforms of the same protein may be destined to a specific compartment, while others are meant to remain in the cytoplasm. An imbalance in distribution of these isoforms can have pathological consequences in many cellular processes. For example, dysregulation of mitochondrial processes has been linked to numerous rare human diseases^7,85^. The N-terminal switching-dependent localization and its underlying molecular mechanism thus represent a novel, hitherto underappreciated phenomenon, the significance of which we are only beginning to recognize.

### Limitations of the Study

This study has several limitations. One of them is that, with the exception of eIF1 overexpression (Figure 2), we have not examined in detail the interplay of our eIF3c mutants with other regulators of TIS selection, such as eIF1, eIF1A and eIF2. Biochemical delineation of eIF5-CTD domain interaction dynamics with eIF1, eIF1A, eIF2β, and eIF3c mutants *in vitro* would shed light on potential synergistic or antagonistic effects of their binding. However, the actual dynamics of these interactions in the PIC is constrained by the complex context of RNA-RNA and RNA-protein interactions, which rapidly change to accommodate the TIS. Our study, therefore, lacks structural insight into the architecture of the PIC with the abolished eIF3c-eIF5 interaction. Nonetheless, these studies are currently under way. While our findings support the existence of altered N-terminal isoforms with potentially altered subcellular localization, verifying their size differences by western blot proved difficult^7^. The main reason is that targeting signals for mitochondrial import and ER secretion are proteolytically removed upon entry into these organelles and the truncated products tend to be comparable in size to isoforms initiated from downstream start codons, making it difficult to distinguish the two forms on an SDS-PAGE gel based on migration alone^7^. To systematically identify alternative N-terminal protein isoforms and their mislocalization within the cell, it is, therefore, necessary to employ highly sensitive approaches enabling detection of protein N-termini (N-terminomics). Although this was not feasible given 1) the already large scale of this study and 2) the high technical demands of this task, we are currently developing a new method in collaboration with the specialized mass spectrometry laboratory to provide this evidence. Finally, it would also be highly desirable to generate homozygous mice bearing our mutations and to explore the impact of dysregulated TIS selection in the context of the entire organism, which is one of your main goals for the future.

## RESOURCE AVAILABILITY

### Lead contact

Requests for further information and resources should be directed to and will be fulfilled by the lead contact, Leos Valasek.

### Materials availability

Reagents generated in this study will be made available upon request.

### Data and code availability

All data are available in the main text or the supplementary materials with the following exceptions. The high-throughput sequencing datasets were deposited into the ArrayExpress collection in BioStudies under the accession numbers: E-MTAB-17426 and E-MTAB-17427; access it through: https://www.ebi.ac.uk/biostudies/arrayexpress/studies/E-MTAB-17426?key=6d2012b4-21bf-4a6d-902c-f16bdc4ac088, and https://www.ebi.ac.uk/biostudies/arrayexpress/studies/E-MTAB-17427?key=5391922f-142b-4645-a9e0-72d65e347121. All custom computer codes and scripts that were wrote for this project are publicly available on https://github.com/lml9331/eIF3_start_recognition.

## ACKNOWLEDGEMENTS

This work was supported by the Czech Science Foundation grant (24-10013S to L.S.V.), ERDF and MEYS (CZ.02.01.01/00/22_008/0004575 RNA for therapy to L.S.V.), Praemium Academiae grant provided by the Czech Academy of Sciences (to L.S.V.), ELIXIR CZ research infrastructure project (MEYS Grant No: LM2023055), including access to computing and storage facilities, and US National Institutes of Health grants (AG064690 and GM145306 to J.D.P.; T32AG052909 to S.H.; and 5R35GM127089 to C.V.).

We thank Alena Pecinová, Petr Pecina and Tomáš Mráček from the Institute of Physiology, CAS in Prague for their assistance with the respirometry measurements and inspiring discussions, and Veronika Kincová for the overall support. We also thank all members of our laboratories for fruitful discussions and helpful comments on the manuscript.

## AUTHOR CONTRIBUTIONS

F.T., T.P., A.H. and L.S.V. conceived the study and designed the research. F.T., T.P., A.M.S., R.G., L.L.Y., J.V., M.P.M., P.K., S.H., K.F., S.W., and A.H. performed the experiments and analyzed the data. F.T., T.P., A.M.S., R.G., L.L.Y., J.V., M.P.M., P.K., S.H., K.F., S.W., J.D.P, C.V., A.H. and L.S.V. interpreted the data. F.T., A.M.S., R.G., L.L.Y., J.V., J.D.P, C.V., A.H. and L.S.V. wrote the manuscript. L.S.V. administered and supervised the project.

## STAR★METHODS

Detailed methods are provided in the online version of this paper and include the following:

Key resources table

Materials and Methods

Supplementary Text

Figs. S1 to S9

Tables S1 to S5

Supplementary Excel files 1 – 3.

Data files such Excel Source Data and Raw Data – uncropped gels for the corresponding Figures.

