## Supplementary material for "Human eIF3 and eIF5 coordinate AUG recognition to sustain energy homeostasis": sup figures and text

### SUPPLEMENTARY INFORMATION

#### MATERIALS AND METHODS

##### Construction of plasmids

All Recombinant DNA plasmids and oligonucleotides/genestrings used for cloning are listed in Supplementary Tables S2 – 3, respectively.

pGL4-SV40-AAG-firefly luc2 was made by inserting the *Hind*III-*Bsr*GI digested genestring pGL4_AAG_LUC into *Hind*III-*Bsr*GI digested pGL4-SV40-firefly luc2.

pGL4-CMV-h3c-7-325, pGL4-CMV-h3c-1-325 R3N, pGL4-CMV-h3c-1-325 F4A, pGL4-CMV-h3c-1-325 F5A, pGL4-CMV-h3c-1-325 F4A F5A and pGL4-CMV-h3c-1-325 R3N F4A F5A were made by inserting the *Pme*I-*Fse*I digested PCR products obtained with primer AH-h3c-325-FseI-R in combination with TP-h3c-7-325-PmeI-FW, TP-h3c-1-325-R3N-PmeI-FW, KP_pGL4-CMV_h3c_1-325_F4A_F, KP_pGL4-CMV_h3c_1-325_F5A_F, TP-h3c-1-325-F4A-F5A or TP-h3c-1-325-R3N-F4A-F5A respectively, using pGL4-CMV-h3c-1-325 as a template into *Pme*I-*Fse*I digested pGL4-CMV-h3c-1-325.

pGL4-CMV-h3c R3N F4A F5A was made by inserting the *Pme*I-*Fse*I digested PCR product obtained with primers TP-h3c-1-325-R3N-F4A-F5A and AH-h3c-FseI-R using pGL4-CMV-h3c as a template into *Pme*I-*Fse*I digested pGL4-CMV-h3c.

pGL4-CMV-eIF3c-1-180-WT-c-FLAG, pGL4-CMV-eIF3c-1-180-R3N-c-FLAG, pGL4-CMV-eIF3c-1-180-F4A-c-FLAG, pGL4-CMV-eIF3c-1-180-F5A-c-FLAG, and pGL4-CMV-eIF3c-1-180-R3N-F4A-F5A-c-FLAG were made by inserting the *Hind*III-*Fse*I digested PCR product obtained with forward primers eIF3c_HindIII_kATG_WT_F, eIF3c_HindIII_kATG_R3N_F, eIF3c_HindIII_kATG_F4A_F, eIF3c_HindIII_kATG_F5A_F, and eIF3c_HindIII_R3NF4AF5A_F and reverse primers eIF3c_FseI_FLAG_180a_R/ FseI_FLAG_STOP_uni_R using pGL4-CMV-h3c as a template into *Hind*III-*Fse*I digested pGL4-CMV-firefly luc2.

pGL4-CMV-eIF3c-1-325-WT-c-FLAG and pGL4-CMV-eIF3c-1-325-R3N-F4A-F5A-c-FLAG were made by inserting the *Hind*III-*Fse*I digested PCR product obtained with forward primers eIF3c_HindIII_kATG_WT_F and eIF3c_325_NAA_T95_FWD, respectively, and reverse primers eIF3c_FseI_FLAG_325_R/ FseI_FLAG_STOP_shrt_R using pGL4-CMV-h3c as a template into *Hind*III-*Fse*I digested pGL4-CMV-firefly luc2

pGL4-CMV-eIF3c-1-913-R3N-F4A-F5A-c-FLAG was prepared by inserting the *Pme*I-*Blp*I digested fragment of pGL4-CMV-eIF3c-1-325-R3N-F4A-F5A-c-FLAG into *Pme*I-*Blp*I pGL4-CMV-eIF3c-1-913-FLAG.

pGL4-CMV-eIF3c-1-913-FLAG was prepared by inserting *Sph*I-*Fse*I digested PCR product obtained with AH-h3c-SphI and AH-h3c-FLAG2-R using pGL4-CMV-h3c as a template into *Sph*I-*Fse*I digested pGL4-CMV-h3c.

All pX330-h3c-gRNA plasmids were made by inserting annealed oligos into *Bbs*I digested pX330-U6-Chimeric_BB-CBh-hSpCas9 Addgene #42230. The used oligos are listed:

pX330-h3c- gRNA_1: pX330-h3c-qRNA_1_sense and pX330-h3c-gRNA_1_antisense
pX330-h3c- gRNA_2: pX330-h3c-gRNA_2_sense and pX330-h3c-gRNA_2_antisense
pX330-h3c- gRNA_3: pX330-h3c-gRNA_3_sense and pX330-h3c-gRNA_3_antisense
pX330-h3c- gRNA_5: pX330-h3c-gRNA_5-sense and pX330-h3c-gRNA_5-antisense
pX330-h3c- gRNA_6: pX330-h3c-gRNA_6-sense and pX330-h3c-gRNA_6-antisense

All pARV-GFP-RFP-gRNA plasmids were made by inserting annealed oligos into *Eco*RV-*Pvu*I digested pARV-GFP-RFP. The used oligos are listed:
pARV-GFP-RFP-gRNA_1-ON_target: pARV-h3c-gRNA_1-on_target-sense and pARV-h3c-gRNA_1-on_target-antisense
pARV-GFP-RFP-gRNA_1-off_target_1: pARV-h3c-gRNA_1-off_target1-sense and pARV-h3c-gRNA_1-off_target1-antisense
pARV-GFP-RFP-gRNA_1-off_target_2: pARV-h3c-gRNA_1-off_target_2-sense and pARV-h3c-gRNA_1-off_taget_2-antisense
pARV-GFP-RFP-gRNA_1-off_target_3: pARV-h3c-gRNA_1-off_target_3-sense and pARV-h3c-gRNA_1-off_target_3-antisense
pARV-GFP-RFP-gRNA_2-ON_target: pARV-h3c-gRNA_2-on_target-sense and pARV-h3c-gRNA_2-on-target-antisense
pARV-GFP-RFP-gRNA_2-off_target_1: pARV-h3c-gRNA_2-off_target_1-sense and pARV-h3c-gRNA_2-off_target_1-antisense
pARV-GFP-RFP-gRNA_2-off_target_2: pARV-h3c-gRNA_2-off_target_2-sense and pARV-h3c-gRNA_2-off_target_2-antisense
pARV-GFP-RFP-gRNA_2-off_target_3: pARV-h3c-gRNA_2-off_target_3-sense and pARV-h3c-gRNA_2-off_target_3-antisense
pARV-GFP-RFP-gRNA_2-off_target_4: pARV-h3c-gRNA_2-off_target_4-sense and pARV-h3c-gRNA_2-off_target_4-antisense
pARV-GFP-RFP-gRNA_2-off_target_5: pARV-h3c-gRNA_2-off_target_5-sense and pARV-h3c-gRNA_2-off_target-5-antisense
pARV-GFP-RFP-gRNA_3-ON_target: pARV-h3c-gRNA_3-on_target-sense and pARV-h3c-gRNA_3-on_target-antisense
pARV-GFP-RFP-gRNA_3-off_target_1: pARV-h3c-gRNA_3-off_target_1-sense and pARV-h3c-gRNA_3-off_target_1-antisense
pARV-GFP-RFP-gRNA_3-off_target_2: pARV-h3c-gRNA_3-off_target_2-sense and pARV-h3c-gRNA_3-off_target_2-antisense
pARV-GFP-RFP-gRNA_3-off_target_3: pARV-h3c-gRNA_3-off_target_3-sense and pARV-h3c-gRNA_3-off_target_3-antisense
pARV-GFP-RFP-gRNA_5-ON_target: pARV-h3c-gRNA_5-on_target-sense and pARV-h3c-gRNA_5-on_target-antisense
pARV-GFP-RFP-gRNA_5-off_target_1: pARV-h3c-gRNA_5-off_target_1-sense and pARV-h3c-gRNA_5-off_target_1-antisense

pARV-GFP-RFP-gRNA_5-off_target_2: pARV-h3c-gRNA_5-off_target_2-sense and pARV-h3c-gRNA_5-off_target_2-antisense
pARV-GFP-RFP-gRNA_5-off_target_3: pARV-h3c-gRNA_5-off_target_3-sense and pARV-h3c-gRNA_5-off_target_3-antisense
pARV-GFP-RFP-gRNA_6-ON_target: pARV-h3c-gRNA_6-on_target_sense and pARV-h3c-gRNA_6-on_target-antisense
pARV-GFP-RFP-gRNA_6-off_target_1: pARV-h3c-gRNA_6-off_target_1-sense and pARV-h3c-gRNA_6-off_target_1-antisense
pARV-GFP-RFP-gRNA_6-off_target_2: pARV-h3c-gRNA_6-off_target_2-sense and pARV-h3c-gRNA_6-off_target_2-antisense
pARV-GFP-RFP-gRNA_6-off_target_3: pARV-h3v-gRNA_6-off_target_3-sense and pARV-h3c-gRNA_6-off_target_3-antisense

pGL4-CMV-h3c-exon1-gRNA1-WT was made by inserting the *Eco*RI-*Fse*I digested PCR product obtained with primers TP-h3c-5UTR-EcoRI-FW and TP-h3c-intron-FseI-R using genomic DNA isolated from HEK293T cells as a template into *Eco*RI-*Fse*I digested pGL4-CMV-h3c.

pGL4-CMV-h3c-exon1-gRNA1-R3N was made by inserting the *Eco*RI-*Fse*I digested fusion PCR product into *Eco*RI-*Fse*I digested pGL4-CMV-h3c. First step of phusion PCR was performed with primer pairs: i.) TP-h3c-5UTR-EcoRI-FW and TP-gen_h3c-R3N-gRNA1-fushion_pair1_R, ii.) TP-h3c-fushion-pair2_FW and TP-h3c-intron-FseI-R, using genomic DNA isolated from HEK293T cells as a template. Second step was performed with TP-h3c-5UTR-EcoRI-FW and TP-h3c-intron-FseI-R primers using products from first step as a template.

pGL4-CMV-h3c-exon1-gRNA1-F5A was made by inserting the *Eco*RI-*Fse*I digested phusion PCR product into *Eco*RI-*Fse*I digested pGL4-CMV-h3c. First step of phusion PCR was performed with primer pairs: i.) TP-h3c-5UTR-EcoRI-FW and TP-h3c-F5A_gRNA1-fushion_pair1-R, ii.) TP-h3c-fushion-pair2_FW and TP-h3c-intron-FseI-R, using genomic DNA isolated from HEK293T cells as a template. Second step was performed with TP-h3c-5UTR-EcoRI-FW and TP-h3c-intron-FseI-R primers using products from first step as a template.

pGL4-SV40-eIF1-native-firefly luc2 and pGL4-SV40-eIF1-strong-firefly luc2 were made by inserting the *Hind*III-*Apa*I digested genestring eIF1-native_LUC and eIF1-strong_LUC (where first 12 codons of firefly were preserved) into *Hind*III-*Apa*I digested pGL4-SV40-firefly luc2. pGL4-SV40-eIF5-native-firefly luc2 and pGL4-SV40-eIF5-uORF-less-firefly luc2 were made by inserting the *Xho*I-*Apa*I digested genestring eIF5-native_LUC and eIF5-uORF-less_LUC (where first 12 codons of firefly were preserved) into *Xho*I-*Apa*I digested pGL4-SV40-firefly luc2.

pGL4-SV40-Luc2-weak, pGL4-SV40-Luc2-medium, pGL4-SV40-Luc2-good, and pGL4-SV40-Luc2-strong were made by inserting the *Xho*I-*Apa*I digested PCR product obtained with forward primer TP_SV40_FW and reverse primers TP_SV40-weak, TP_SV40_medium, TP_SV40_good, and TP_SV40_strong, respectively, using pGL4-SV40 as a template into *Xho*I-*Apa*I digested pGL4-SV40.

pGL4-SV40-NAT1-GTG and pGL4-SV40-NAT1-ATG were made by inserting the *Hind*III-*Bsr*GI digested genestrings NAT1_AUG_GTG or NAT1_AUG_AUG into *Hind*III-*Bsr*GI digested pGL4-SV40, according to[1] .

**Bacterial cultivation and transformation**

*E. coli* culture was grown in LB liquid medium or on LB agar plates at 37°C. The medium was supplemented with 150 μg/ml ampicillin or 160 μg/ml blasticidin depending on the plasmid. Transformation was carried out by heat shock in the case of DH5α and BL21 Star (DE3) strains, or by electroporation in the case of DH10B strain.

**Mammalian cells cultivation**

HEK293T (ATCC, CRL-3216; RRID: CVCL_0063) and HeLa (ATCC Cat# CCL-2, RRID:CVCL_0030) cells were grown in DMEM (Sigma, cat #D6429) supplemented with 10% FBS (Gibco, cat # 10270–106) at 37°C and 5% $CO_2$. If not stated otherwise, cells were grown without antibiotics. Cells were regularly tested for mycoplasma infection.

**siRNA transfection**

HeLa cells were grown in 6WP or Ø15 cm dishes for Luciferase assay or CoIP experiments respectively. The slower growth rates of $eIF3e^{KD}$ and $eIF3c^{KD}$ cells were compensated by the number of seeded cells. Twenty-four hours after seeding, cells were transfected with targeted or non-targeted ON-TARGETplus siRNA cocktail system from Dharmacon at a final concentration of 5 nM. Catalog numbers for all siRNAs used in this study are listed in Supplementary Table S3. INTERFERin (Polyplus, cat # 101000016) was used as a transfection reagent. Cells were harvested 72 hr after transfection at approximately 80% confluency.

**Cell lysate preparation**

Cells were washed with ice cold 1 x PBS and lysed directly on the dish in buffer A (10 mM HEPES (pH 7.5), 62.5 mM KCl, 10 mM $MgCl_2$) supplemented with 1 mM DTT, 1 mM PMSF, 1 μg/ml Aprotinin, 1 μg/ml Leupeptin, 1 μg/ml Pepstatin, mini–Complete EDTA-free (Roche) 1 tablet/5 ml, 1% Triton X-100. The resulting lysate was collected and cleared by centrifugation. For Polysome profiling, 43S-48S assembly analysis and 43S-48S CoIP experiments, cells were pre-treated with Cycloheximide (Sigma, cat # C7698) at a final concentration of 100 μg/ml for 1 min prior to harvesting and same concentration of Cycloheximide was also added to the lysis buffer.

**SDS-PAGE and Western blot**

All samples were resolved using SDS-PAGE followed by western blotting. All primary antibodies used in this study are listed in Supplementary Table S4. The signal was developed using SuperSignal West Femto Maximum Sensitivity Substrate (Thermo Fisher Scientific, cat # 34096) and detected in a G-Box imager (Syngene) using a series of varying exposure times. Signals were processed with Quantity One (Bio-Rad) and GelAnalyzer 23.1.1. Only signals from the same strips and with the same exposure times were compared.

**Co-Immunoprecipitation assays from lysates - siRNA treated HeLa cells**

eIF3 complexes were immunoprecipitated from lysates, prepared as above, using Protein G Mag Sepharose Xtra (Cytiva, cat # 28-9670-70) with anti-eIF5 (Santa Cruz, cat # sc-28309) antibody. Lysate containing 1 mg of total protein was used for each sample and pre-incubated with 35 μl of the eIF5 (Santa Cruz Biotechnology, sc-28309) antibody for 2 h while rotating at 4°C, then beads were added, and the mixture was incubated overnight at 4°C while rotating. Three washing steps were performed with buffer A and coimmunoprecipitated proteins were eluted in a denaturing loading buffer by boiling. The eluate was analyzed by SDS-PAGE followed by Western blotting.

**Co-Immunoprecipitation assays from HCHO crosslinked lysates**

HEK293T cells were crosslinked on the cell culture dish for 5 min with formaldehyde at a final concentration of 0.8% in ice cold 1x phosphate-buffered saline (PBS), followed by 5 min incubation with 75 mM Glycine in ice cold 1x PBS to quench the crosslinking reaction. Fixed cells were then washed with ice-cold 1×PBS and lysed directly on the cell culture dish as above. The resulting lysate was cleared by centrifugation. For coimmunoprecipitation of eIF3 complex, lysates containing 1 mg of total protein and 24 μl of anti-eIF3a (Cell Signaling Technology, cat #3411) were utilized per assay and Co-IP experiments were performed as above.

**Luciferase-based TIS selection assays (LuTISsa).**

LuTISsa assays were performed in two formats producing consistently comparable results. Please see the figure legends. *qPCR format*: cells were transfected (in siRNA experiments after 48 hours of silencing) with firefly luciferase reporter plasmids (ranging between 700-1500 ng for 6WP transfections) using TurboFect (Thermo Scientific; catalog no. R0531) according to the manufacturer´s instruction. Cells were harvested 24 hours after reporter transfection in 1x Glo lysis buffer (Promega). The lysate was divided into two parts: one part of the lysate was used for detection of luciferase activity with Bright-Glo™ Luciferase Assay (Promega) and the other was used for RNA isolation followed by

RT-qPCR. To control for a potential material loss, spike RNA (*S. cerevisiae* RPL41a) was added to the lysate prior to RNA isolation. Total RNA was isolated using RNA Blue reagent (TopBio, cat # R013), contaminating DNA was removed by TurboDNAse treatment, which was subsequently inactivated by Turbo-DNase Inactivation Reagent (Invitrogen). Reverse transcription was performed (Applied Biosystems; catalog no. 4368814) and the resulting cDNA was diluted 5 times. The levels of firefly luciferase, Spike, ALAS, and 18S mRNAs (endogenous targets) were analyzed with quantitative PCR in triplicates. To account for variability in plasmid transfection efficiency, the firefly luciferase mRNA levels, normalized to Spike/ALAS/18S, were used for the luminescence signal normalization. The ratio of initiation from individual near-cognate codons to AUG was then calculated for each treatment. *Renilla normalization format*: Cell seeded in 24WP were transfected with 250 ng of firefly luciferase reporter plasmids together with 33 ng of a normalization vector encoding Renilla luciferase using TurboFect (Thermo Scientific; catalog no. R0531) in three technical replicates. In overexpression experiments with eIF1, eIF5, and the eIF3c-NTD fragments, 250 ng of eIF1 / eIF5 / eIF3c expressing vectors or mock vector were co-transfected with the luciferase reporters. Cells were harvested 24 hours after plasmid transfection in 1x Glo lysis buffer (Promega) and the activities of firefly and renilla luciferases were detected using BrighGlo™ and RenillaGlo™ as substrates (Promega). Renilla-normalized firefly luciferase signals were used for further calculations (see figure legends).

**GST PD assay**

Glutathione S-transferase (GST) pull down experiments with GST fusions and *in vitro* synthesized $^{35}$S-labeled polypeptides were conducted as described previously[2]. Briefly, individual GST-fusion proteins were expressed in *Escherichia coli* (BL-21 Star DE3). Bacterial culture was grown at 37°C in the LB medium to OD 0.6-0.8 and the synthesis of GST-fusion proteins were induced by the addition of 1 mM IPTG. After 2 h of shaking at 37°C, the cells were harvested, resuspended in a Phosphate-buffered saline (PBS), and subjected to mechanical lysis with a subsequent agitation in the presence of 1-1.5% Triton X-100 for 30 min at 4°C. The GST-proteins were then immobilized on glutathione Sepharose beads (GE Healthcare, cat # GE17-0756-01) from the pre-cleaned supernatant, followed by three washing steps with the 1 ml of phosphate buffered saline.$^{35}$S-labeled polypeptides were produced *in-vitro* by the TnT® Quick Coupled Transcription/Translation System (Promega cat # L1170) according to the vendor's instructions. To examine the binding, individual GST fusions were incubated with $^{35}$S-labeled proteins at 4°C for 2 h in buffer B (20mM HEPES (pH 7.5), 75 mM KCl, 0.1 mM EDTA, 2.5 mM $MgCl_2$, 0.05% IGEPAL, 1 mM DTT). Subsequently, the beads were washed three times with 1 ml of PBS and interacting proteins were separated by SDS-PAGE. Gels were first stained with Gelcode Blue stain reagent (Thermofisher, cat # 24592) and then subjected to autoradiography.

**RT-qPCR**

Endogenous *EIF3C*, *EIF1* and *EIF5* mRNA levels in wt, R3N and triple mutant HEK293T cells were analyzed by RT-qPCR using the same general qPCR workflow as for the LuTISsa assay. Total RNA was isolated from the indicated stable cell lines, reverse-transcribed into cDNA, and analyzed using target-specific primers. qPCR reactions were performed in four technical replicates. Primer concentrations and annealing temperatures were optimized prior to analysis. Relative mRNA levels were calculated using ALAS and 18S rRNA as reference genes, with the arithmetic mean of their Ct values used for normalization. Data were expressed relative to wt cells set to 1.

**Polysome profiles**

HEK293T cell lysates prepared from cells pre-treated with Cycloheximide (see Cell lysate preparation) were separated by high velocity sedimentation through a 20-60% sucrose gradient at 39000 rpm for 2.5 h using the Beckman Coulter SW41Ti rotor. Polysome profiles were recorded as absorbance at 260nm ($A_{260}$) using Biocomp Instruments Fractionation and Profiling system.

**The 43S-48S PIC assembly and Co-IP experiments**

HEK293T cell lysates from cells pre-treated with Cycloheximide were prepared (see Cell lysate preparation). The resulting lysates were cleared by centrifugation (5 min / 16,000 g / 4°C) and absorbance at 260 nm was measured. 12U of $A_{260}$ aliquots were prepared and flash frozen in liquid nitrogen and stored at -80°C until used in an experiment. For the 43S-48S PIC assembly assay, as well as for the 43S-48S PIC Co-IP experiment using anti-eIF3a antibodies, one 12U aliquot was used per sample. For the 43S-48S PIC Co-IP experiment using anti-eIF5 antibodies, two 12U aliquots were used per sample (material obtained from two gradients was pooled together after centrifugation). In order to enrich the sample of 43S-48S PICs, 1 µl of RNase I (Thermofisher, cat # AM2295) was added per 12U of lysate and incubated at 24°C for 10 min. Digestion was quenched with 2 µl of SuperaseIN RNase inhibitor (Thermofisher, cat # AM2696). Each 12U aliquot was then separated by high velocity sedimentation through a 7–30% sucrose gradient containing a top-to-bottom increasing concentration of formaldehyde at 22,200 rpm for 18.5 h using the Beckman Coulter SW41Ti rotor as in[3]. Such a gradient was prepared by mixing 30% sucrose in buffer A (10 mM HEPES [pH 7.5], 62.5 mM KCl, 10 mM MgCl2, 1mM DTT) containing 0.05% formaldehyde with 7% sucrose in buffer A lacking formaldehyde, using the Gradient Master 108 (Biocomp Instruments). Fractions of 620 µl were collected and either precipitated with 100% ethanol overnight at –20°C for the 43S-48S PIC assembly analysis or pooled and used for 43S-48S PIC Co-IPs (see below). After precipitation, the pellet was dried and dissolved in 1× standard loading buffer. Samples were incubated at

95°C for 15 min to reverse cross-linking and analyzed by SDS-PAGE followed by western blotting.

For the 43S-48S PIC Co-IPs, four fractions containing either ribosome free factors or the 43S-48S PICs were pooled separately. 500 µl of pooled fractions was taken for Input sample and precipitated by ethanol as above. Triton X-100 was added to the remaining pool to the final concentration of 0.1%. 30 µl of eIF3a or eIF5 antibodies was added to the pool and incubated for 2 h while rotating at 4°C. The beads (Protein G Mag Sepharose Xtra (Cytiva, cat # 28-9670-70)) were then added, and the mixture was incubated overnight at 4°C while rotating. Three washing steps were performed with buffer A containing 0.1% Triton-X100 and one final washing step with buffer A only. Coimmunoprecipitated proteins were eluted in a denaturing loading buffer by boiling. The eluate was analyzed by SDS-PAGE followed by western blotting. Catalog numbers for all antibodies used in this study are listed in Supplementary Table S4.

**Growth assays, protein content and cell size**

A total of $6.6 \times 10^5$ cells were serially diluted in culture medium at a 2:1 ratio across a 96-well plate and incubated for 48 hours. Seeding was performed in triplicates. Following incubation, resazurin was, according to the manufacturer's recommendations, added to each well at the final concentration of 40 µM and cells were incubated for additional 3 h at 37 °C and 5% $CO_2$. Fluorescence was subsequently measured using a Tecan Infinite 200 PRO reader with excitation/emission wavelengths set at 560/590 nm. Fluorescence intensity values were plotted against the logarithm (base 10) of the initial cell number per well and analyzed in GraphPad Prism 8 using a four-parameter logistic (4PL) nonlinear regression model.

To evaluate the total protein content, wt and mutant cell lines were stained with CellTrace™ Violet dye (Thermo Fisher Scientific), which covalently binds to free amine groups. Briefly, cells were washed with PBS and incubated with CellTrace Violet at a final concentration of 0.1 µM in 2 ml pre-warmed PBS per well for 20 min at 37°C. Control samples received the corresponding amount of DMSO diluted in PBS. After staining, cells were washed with complete DMEM containing FBS, detached with 0.025% trypsin, and neutralized with complete medium. Cells were collected by centrifugation at 300×g for 5 min, resuspended in phenol red-free DMEM, and analyzed by flow cytometry. Propidium iodide (1 µg/ml) was added immediately before acquisition to exclude dead cells. A total of 100,000 events per sample were acquired and data were analyzed using FlowJo™ v11 software (FlowJo LLC). Relative cell size of the cell lines was evaluated based on the forward scatter (FSC) parameter.

**FLAG affinity precipitation**

HEK293T cells were seeded in Ø10 cm cultivation dish 24 hours before transfection with constructs encoding C-terminally FLAG-tagged eIF3c variants (8 µg of DNA per dish, for transfection procedure details, see the LuTISsa method). Next, the cell lysates were prepared (see Cell lysate preparation) in buffer A supplemented with 1 mM DTT, 1 mM PMSF, 1 µg/ml Aprotinin, 1 µg/ml Leupeptin, 1 µg/ml Pepstatin, mini–Complete EDTA-free (Roche) 1 tablet/5 ml, 1% Triton X-100. The anti-FLAG M2 beads (Sigma, cat #A2220) washed with Buffer A were added to the lysate and incubated for 2h while rotating at 4°C. Next, three washing steps were performed with buffer A containing 0.1% Triton-X100. Coimmunoprecipitated proteins were eluted by addition of FLAG peptide (Sigma, cat #F3290) at final concentration of 250 ng/µl in buffer A. The eluate was analyzed by SDS-PAGE followed by western blotting.

**CRISPR/Cas9 stable cell line generation**

To introduce CRISPR/Cas9 into the cells by plasmid transfection, we employed the system with two plasmids; the first vector pX330-U6-Chimeric_BB-CBh-hSpCas9 (further referred as pX330) encoding both humanized S. pyogenes Cas9 protein and single-guide RNA, and the second pARV-GFP-RFP (further referred as pARV) acting as a reporter plasmid. As for the latter vector, the GFP protein is produced upon pARV transfection and acts as a glowing marker of its efficiency. The production of RFP is more complex. There are two RFP fragments in pARV; the first, short one carries the start codon, the second encodes the rest of RFP starting from the 2nd aa. They are separated by a copy of a given target sequence of interest. When Cas9 and gRNA are produced in cells containing pARV,

| | pX330 | pARV | pARV-off target |
|---|---|---|---|
| gRNA 1 | pX330-h3c-gRNA_1 | ON-target | 1: CH3 trafficking kinesin binding protein exon |
| | | | 2: CH19 guanine nucleotide binding protein exon |
| | | | 3: CH16 intergenic CASKIN1/MLST8 |
| gRNA 2 | pX330-h3c-gRNA_2 | ON-target | 1:CH3 intergenic NECTIN3/DPPA4 |
| | | | 2: CH13 leukecia associated protein 7 intron |
| | | | 3: CH2 intergenic OR5S1P/OR9S24P |
| | | | 4: CH22 ENTHD1 –intron |
| | | | 5: CH10 TNKS2 exon |
| gRNA 3 | pX330-h3c-gRNA_3 | ON-target | 1: CH14 ZBTB42 exon |
| | | | 2: CH15 SPRED1 (5UTR) |
| | | | 3: CH6 ARMC2 (5UTR) |
| gRNA 5 | pX330-h3c-gRNA_5 | ON-target | 1: CH4 intergenic |
| | | | 2: CHX itergenic SSBL2P/centromera |
| | | | 3: CH9 NUTM26 exon |
| gRNA 6 | pX330-h3c-gRNA_6 | ON-target | 1: CH7 GATAD1 intron |
| | | | 2: CH7 intergenic loc105375194/100506236 |
| | | | 3: CH7 TRIM74 exon |

Supplementary Table S1. Off-targets of the selected gRNAs as predicted by CRISTA and CRISPOR tools.

the target sequence between the RFP fragments is recognized and cut by Cas9. Only then do the two parts of RFP recombine and the RFP protein is produced. RFP is therefore a marker for active CRISPR/Cas9 in the cell[4].

Before designing gRNAs, we isolated genomic DNA from the HEK293T cell line and performed PCR on the target region to ensure that the actual sequence matched the annotated sequence. We then carried out a sequence search for protospacer adjacent motifs (PAMs) and potential gRNA targets. Since the distance of the Cas9 cut site from the mutation site negatively affects the efficiency of introducing the mutation, we selected five of them that were closest to the mutation site (referred as gRNA 1, 2, 3, 5 and 6, see Supplementary Table S3). We then predicted *in silico* the ON-target and off-target activity of the gRNAs using the CRISTA program[5] (Supplementary Table S1). All 5 gRNAs performed similarly, we therefore decided to test all of them *in vivo*. (Nonetheless, it is noteworthy that gRNAs 2, 3 and 6 each predicted a single off-target with a very high Crista score (around 0.7). In addition, gRNA2 has a few hits with a somewhat higher score (around 0.5)). We first cloned the spacer part of individual gRNAs into the pX330 vector.

Furthermore, we cloned three off-target sequences for each gRNA (according to highest CRISTA score) into the reporter plasmid pARV; among these, we included at least one off-target located in an exon for relevance. We prepared 5 off-targets for gRNA2, as it was predicted to have a high number of off-targets with a relatively high CRISPOR score (https://crispor.gi.ucsc.edu/, Supplementary Table S1).

To test the off-targets of gRNAs, HEK293T cells were seeded onto 24 well-plate (40,000 cells/well), and after 24 hours, the cells were co-transfected with combination of pX330 and pARV plasmids carrying either ON-target or off-target sequence using TurboFect transfection reagent according to the manufacturer's recommendations. Briefly, 250 ng of each plasmid was mixed with 1.5 µl of TurboFect in 100 µl of serum-free DMEM. After two days, we detected CRISPR/Cas9 activity by fluorescence microscopy (Insert 1). All gRNAs exhibited strong ON-target activity. gRNA3 and gRNA6 showed high activity on off-target 1. In addition, we detected a weak RFP signal in cells transfected with gRNA5 off-target 2. Therefore, we selected gRNA1 for generation of all mutants (single and triple knock-ins), since it has better *in silico* specificity prediction compared to gRNA2.

Next, we proceeded with the donor design. Donor ssODNs (single-stranded oligodeoxynucleotides) (Supplementary Table S3) were constructed as follows. First, ssODN corresponds to the gRNA1 target strand and contains symmetrical 40 nt homology arms at each site, starting either from the cut site or from the mutation. Second, the PAM sequence is mutated in the donor sequence. Conveniently, no additional PAM substitution is required for the R3N donor (as well as for R3N F4A F5A donor), as the codon encoding R3 also acts as the PAM sequence for gRNA1. In the case of F5A ssODN, however, a same-sense single nucleotide substitution is introduced to mutate the PAM sequence. Moreover, all mutations were designed to create a new restriction enzyme (RE) site (Figure 2A). The introduced RE sites were used for genotyping the generated HEK293T clones. The substitution encoding R3N is recognized by *Xmn*I, the substitutions encoding

F5A were extended by an additional single nucleotide, same-sense mutation to create a restriction site for *Nru*I. The substitution encoding R3N F4A F5A is recognized by *Tse*I, as a part of substitution encoding F4A single nucleotide mutation. Mammalian codon usage was considered in the design of the ssODN. Furthermore, we added three phosphorothioate bond modifications at both ends of ssODN to increase its stability[6].

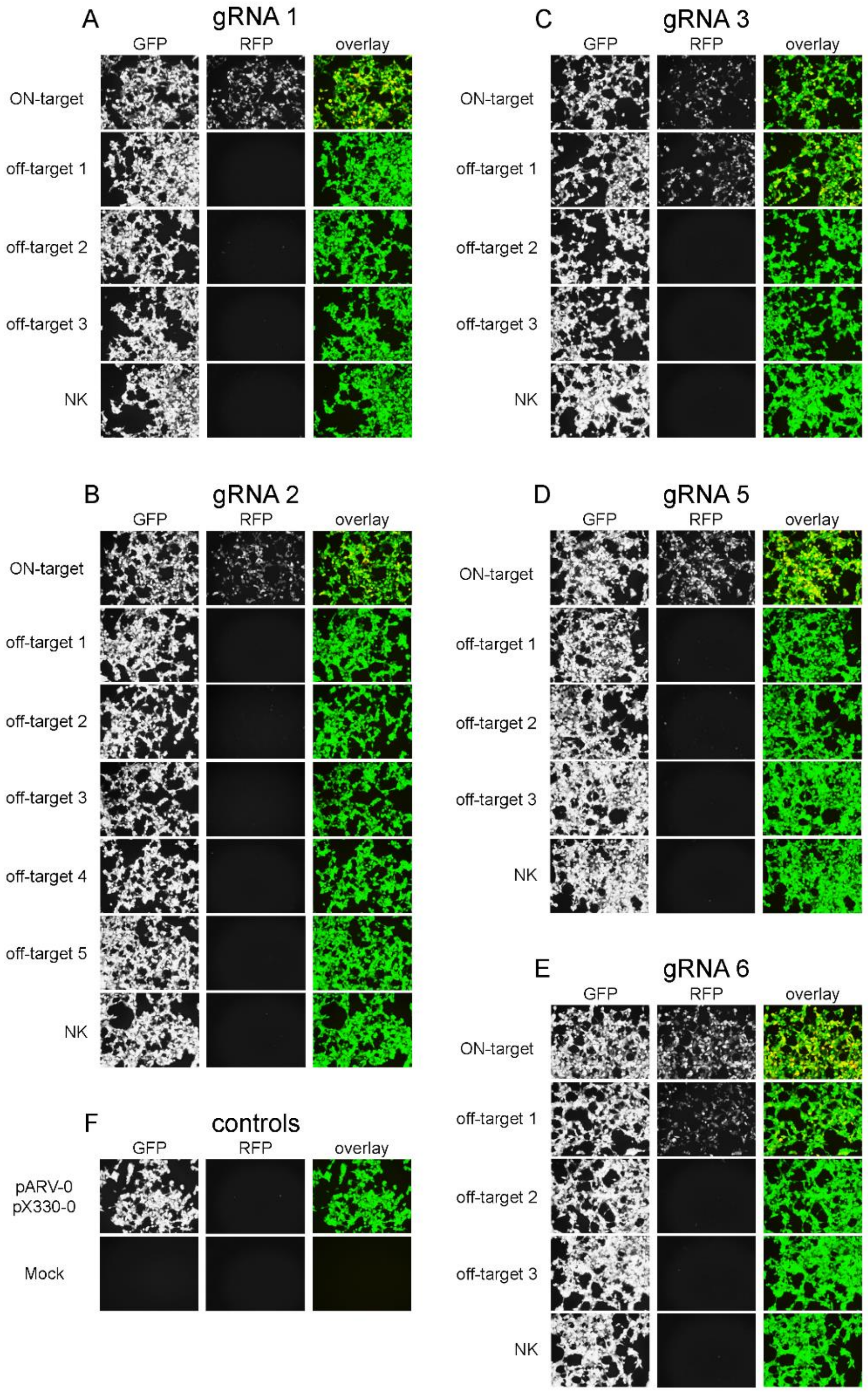


Insert 1. pARV testing of ON-target and potential off-target activity of selected gRNAs.

To establish HEK293T stable cell lines using selected gRNA1 and ssODNs, the cells were seeded into 6WP dish (60,000 cells/well), containing 2 ml DMEM high glucose supplemented with 10% FBS. The following day, the cells were transfected with a combination of plasmids pX330-h3c-gRNA_1, pARV-GFP-RFP-gRNA_1-ON_target and donor ssODN. The reaction was prepared by mixing 300 ng (150 ng in case of the triple substitution) of each plasmid, 60 pmol (30 pmol in case of the triple substitution) of ssODN, 3.6 µl (1.8 µl of the triple substitution) of Turbofect and DMEM without FBS in a total volume of 500 µl. After 20 minutes of incubation at room temperature (RT), the reaction was applied to the cells. The medium was changed after 6 hours and then every day until sorting. To obtain monoclonal cells, the cells were harvested three days after transfection and sorted on BD Influx (BD Biosciences) with a 100 µm nozzle setting (Insert 2). Cells were sorted by RFP signal to a 96 WP dish with conditioned medium (collected from 80 % confluent HEK293T cells) supplemented with antibiotics (high-glucose DMEM supplemented with 9% FBS and PenStrep mixed 70:30 with filtrated conditioned medium, with an additional 5% of the total volume of FBS). Cells were cultured in antibiotic-supplemented medium for one subsequent passage before being transferred back to antibiotic-free medium. Generated clones were used for isolation of genomic DNA and subsequent screening for the RE site in the amplified target sequence.

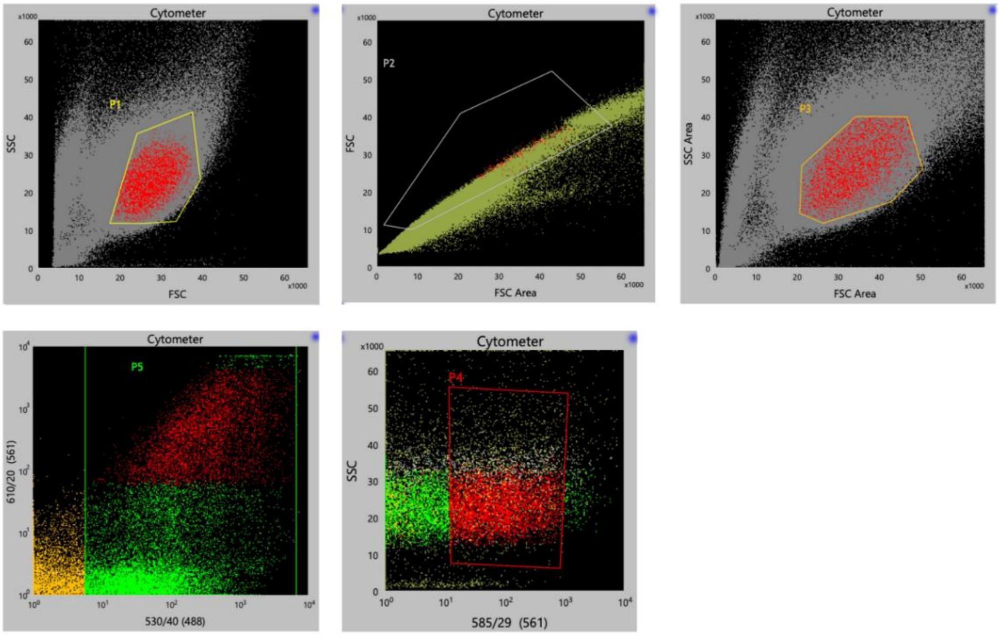


Populations: Cytometer

| Populations | Events | % Total | % Parent |
|---|---|---|---|
| All Events | 500,000 | 100.00% | #### |
| P1 | 379,714 | 75.94% | 75.94% |
| P2 | 357,137 | 71.43% | 94.05% |
| P3 | 324,327 | 64.87% | 90.81% |
| P5 | 22,197 | 4.44% | 6.84% |
| P4 | 4,417 | 0.88% | 19.90% |
| NOT(P1) | 120,286 | 24.06% | 24.06% |

Insert 2. Gating strategy for RFP-GFP double monoclonal sorting.

Genomic DNA from stable HEK293T cell lines was isolated using the DNeasy® Blood & Tissue Kit – Purification of total DNA from Animal Blood or cells (spin-column) and used as a template (40 ng of genomic DNA per reaction) for PCR using the Phire Hot Start II PCR Master Mix with the TP_eIF3c_FW genotyp pair2 and TP_eIF3c_R genotyp pair2 primers. Genomic DNA from wt HEK293T cells and pGL4-CMV-h3c-exon1-gRNA1-WT plasmid were used as negative controls. The pGL4-CMV-h3c-exon1-gRNA1-R3N or pGL4-CMV-h3c-exon1-gRNA1-F5A (pGL4-CMV-h3c-exon1-gRNA1-R3NF4AF5A in case of the triple mutant) plasmids were used as positive controls. The obtained products were purified by QIAprep Spin Miniprep Kit using the PCR purification protocol (QIAGEN). The purified DNA was then used for the RE reaction using the corresponding enzymes (Figure 2A). The resulting DNA fragments were separated electrophoretically on an EtBr agarose gel (data not shown).

After this initial screening, selected positive clones were subjected to further testing. First, the RE analysis was repeated using high fidelity Q5 DNA polymerase and 5X Q5 High GC Enhancer. Next, the obtained PCR products were used for TOPO-TA cloning and sequencing. TOPO-TA cloning was performed as follows. First, a 3´A overhang was generated using Platinum™ Taq DNA Polymerase by subjecting 0.75 pmol of PCR product and 200 µM of dATP to a 25 µl reaction (2 min 94°C, 20 min 72°C). The resulting product was used for cloning using the TOPO™ TA Cloning™ Kit for sequencing according to the vendor’s instructions and then transformed into *E. coli* DH5α by heat shock. We selected 24-36 single colonies per HEK293T clone and sequenced the insert to confirm the knock-in sequence (representative chromatograms are shown in Figure S2A).

In addition, genomic DNA was used as a template for a set of PCR reactions using Q5 polymerase and corresponding primer pairs to analyze larger genomic segments surrounding the CRISPR/Cas9 target region. The resulting PCR products were subjected to size analysis by electrophoresis and sequencing, and we did not detect any indels in any of the clones tested (data not shown).

**Single-molecule assays**

Purification and fluorescent labeling of human ribosomal subunits, and initiation factors, preparation of 5' m7G capped and 3'-biotinylated mRNAs, and execution of real-time single-molecule initiation assays and data analysis were done as described previously[7].

**Ribosome profiling and RNA-seq library**

Ribosome profiling and RNA-seq library preparation and subsequent analysis of sequencing data was carried out similarly to our previous analysis[8]. For footprint (FP) libraries, cycloheximide was added to a final concentration of 100 µg/ml 1 min prior to harvesting. Cells were washed with ice cold 1x PBS and lysed on the dish in buffer A (10 mM HEPES [pH 7.5], 62.5 mM KCl, 10 mM $MgCl_2$) supplemented with 1 mM DTT, 1 mM PMSF, 1 µg/ml Aprotinin, 1 µg/ml Leupeptin, 1 µg/ml Pepstatin, Complete Mini EDTA-free

[Roche, cat # 11836170001] – 1 tablet/5 ml, 1% Triton X-100, and 100 μg/ml Cycloheximide. The resulting lysate was collected and cleared by centrifugation. Aliquots of twelve or five absorbance units (AU) $OD_{260}$ of lysate were flash frozen in liquid nitrogen and used for FP isolation or control polysome profiling, respectively. In case of the $eIF3c^{KD}$ Hela cells, an aliquot of 100 μl was mixed with 1 ml of RNA Blue (TopBio, cat # R013) and taken for RNA isolation and control qPCR to evaluate the level of eIF3c downregulation. When the lysate passed the control experiments (polysome profile, qPCR), one aliquot of 12 AU $OD_{260}$ was thawed on ice and digested with 6 μl (HeLa) or 2 μl (HEK293T) RNAseI (Ambion, cat # AM2295) for 30 min at 24 °C with mild shaking (300 rpm). RNAseI was inactivated with SuperaseIN (Ambion, cat #AM2696). 80S ribosomes were separated by high-velocity sedimentation through a 7% to 30% sucrose gradient at 39,000 rpm for 3.5 h using the SW41Ti rotor. The gradients were scanned at 260 nm to visualize the ribosomal species and fractions containing 80S ribosomes were collected and mixed with RNA Blue. RNA was isolated according to vendor's instructions, FP RNA was size selected on 15% TBE-Urea PAGE (Biorad, cat # 3540092) in range 20–40 nt and size selected FPs were dephosphorylated. In the case of Hela $eIF3c^{KD}$ samples, rRNA depletion was performed using in-house designed biotinylated oligos listed in Supplementary Table S5. In the case of HEK293T eIF3c triple cell line, riboPOOL Kit (siTOOLs, cat # dp-K024-042) was used according to vendor's instructions.

Footprint library was prepared using SMARTer smRNA-Seq Kit (TAKARA, cat # 635029) and sequenced on NovaSeq6000 (Illumina) with read length 150 nt.

For mRNA libraries, the growth media was aspirated and cells were harvested into 5 ml of RNA Blue per Ø 15 cm dish. RNA was isolated according to vendor's instructions. mRNA was isolated from the total RNA using Poly(A)purist MAG kit (Ambion, cat # AM1922), fragmented using RNA Fragmentation Reagents (Ambion, cat # AM8740) and size selected on 15% TBE-Urea PAGE in range 50–100 nt. Size selected mRNA was then dephosphorylated and library was prepared using SMARTer smRNA-Seq Kit (TAKARA, cat # 635029) according to vendor's instructions. mRNA libraries were sequenced on NovaSeq6000 (Illumina) with read length 150 nt.

**Analysis of sequencing data**

Analysis of sequencing data was carried out as described previously[8]. In particular, the first 3 nucleotides on the 5' end were removed, and poly(A) sequence (10xA) was trimmed from the 3'end. Reads shorter than 15 nt were discarded. For FP nt and $eIF3c^{KD}$ libraries, reads longer than 45 nt were also discarded. rRNA and tRNA reads were discarded after alignment with Bowtie2 (Langmead and Salzberg, 2012). Remaining reads were aligned to genome and transcriptome by the STAR aligner[9]. Differential expression analysis was performed by DESeq2[10].

<u>Volcano plots</u>: Volcano plots were generated from DESeq2 translation efficiency (TE) output tables using Python. The underlying DESeq2 tables were processed using

Ensembl gene identifiers, while selected top hits were labelled by HGNC gene symbol for visualization. Genes with (padj) < 0.05 and positive or negative $log_2$ fold change were classified as significantly upregulated or downregulated, respectively; genes not meeting this threshold were classified as unchanged. No additional absolute $log_2$ fold-change cutoff was applied.

The x-axis shows $log_2$ fold change and the y-axis shows $-log_{10}$(padj). Ten most significant upregulated and downregulated genes were labelled by a gene symbol. Data processing and visualization were performed in Python v3.10.19 using pandas v2.3.3, NumPy v2.2.5, and matplotlib v3.10.8.

Function enrichment analysis: Significant TE genes were separated into upregulated and downregulated groups using padj < 0.05 and the sign of the $log_2$ fold change. Gene symbols were used as input, and Enrichr analysis was performed in Python using gseapy v1.1.10. Enrichment was tested separately for upregulated and downregulated gene sets using GO Biological Process 2025, GO Cellular Component 2025, GO Molecular Function 2025, and KEGG 2026 human libraries. The top enriched terms were visualized as horizontal bar plots showing $-log_{10}$ of the Enrichr adjusted p-value.

MitoCarta overlap analysis: Significant TE, FP, and mRNA gene sets were overlapped with the Human MitoCarta3.0 database using Ensembl gene identifiers. MitoCarta entries containing multiple Ensembl identifiers were split before matching. Significance of the overlap between MitoCarta genes and each significant gene set was assessed using a hypergeometric test, separately for upregulated and downregulated genes. For volcano plot visualization, significant TE genes overlapping with MitoCarta were highlighted on the corresponding TE volcano plot and grouped into broad mitochondrial categories, including mtDNA-encoded OXPHOS subunits, mitochondrial ribosomal proteins, nuclear-encoded OXPHOS subunits, mitochondrial translation factors, mitochondrial metabolism, and other mitochondrial genes. MitoCarta overlap analysis and visualization were performed in Python v3.10.19 using pandas v2.3.3, NumPy v2.2.5, and matplotlib v3.10.8.

Read processing

The following packages were used to process the reads: FastQC (v0.11.9) for read quality, Cutadapt (v4.9) for adapter trimming, STAR (v2.7.11a) for alignment to the human hg38 reference genome and Samtools (v1.20) to process the resulting BAM files. Uniquely mapped reads were retained and read length distributions were visualized to determine the length cutoffs for the monosome peak.

Start codon occupancy quantification

All downstream analyses were performed in R (v4.5.1). Reads were initially normalized to total sequencing depth across reads of length 15–50 nt. To increase stringency for downstream metrics, reads were further restricted to 15–40 nt matching the most likely monosome peak. In both eIF3c$^{KD}$ and triple, all replicates showed consistently lower read numbers than wt, prompting us to scale the samples. In the eIF3c$^{KD}$ dataset, the eIF3c$^{KD}$ samples had on average 0.39× the uniquely mapped reads of wt samples, and in the triple dataset, the triple samples had on average 0.84× the uniquely mapped reads of wt. eIF3c$^{KD}$ and triple occupancies were therefore scaled by 0.39 and 0.84, respectively, to preserve the relative read representation.

Canonical and non-canonical start codons in the 5'UTR were predicted from sequence. For this, isoform transcripts were selected first by the APPRIS annotation, then by the longest transcript. The following start codons were predicted from these transcripts: canonical 5'UTR start codons, non-canonical 5'UTR start codons and the AUG start codons that initiate the main translon. No start codons in the main translon were used apart from the main start. The P-site offset was determined by visualizing the average 5′ read end occupancy around predicted start codons. Based on this, the average number of reads at −12 and −13 was used as the P-site offset. The background signal was defined as the mean read count at positions −15 to −20 upstream of the start codon. The background signal was subtracted from the P-site offset for each start codon to correct for non-initiation noise in the 5'UTR. Occupancy values were averaged across replicates to yield a single initiation metric per start codon. For the genome-wide analyses, we retained start codons with measurements in at least 2 replicates per condition (wt or eIF3$^{KD}$/triple).

Downstream occupancy analysis for true initiation

The average read count across the two codons immediately downstream of the start codon was also calculated, with the same background subtraction, by aggregating over replicates per position, then averaging over the -9 to -4 position. For the downstream occupancy analysis, several filters were applied to ensure the start sites used were initiating with downstream occupancy. Firstly, start codons were retained if they had non-zero occupancy in at least 3 of the replicates in either condition in each dataset. Start codons also were filtered for an average initiation and downstream occupancy greater than pseudocounts used in the analysis in wt and eIF3$^{KD}$ or TM conditions. After filters were applied for occupancy at the P-site and downstream of the P-site, the RNA correction was applied, and the two values were correlated with each other in a scatter plot.

The periodicity metric was calculated by first averaging the reads across samples per position (from -9 to -1) then per codon (downstream codon 1 and 2). The reads at each position per codon were ranked from 1, 2 and 3, where 3 would be the highest occupancy at a position. The ranks were then averaged to obtain an average rank per frame.

Filtering

An intercorrelation matrix was used to calculate the spearman correlation between wt replicates. The average correlation between replicates was calculated for every combination of wt replicates, and the two groups with the smallest difference were taken as the two wt groups. The average across replicates was taken for each group, and only one of them was used as the denominator to calculate fold change as $log_2$(eIF3c$^{KD}$ / wt). For the mutant data, the two replicates that were least correlated were averaged. Fold change in the mutant data was then calculated with the average of the two out of three replicates. For analysis on fold changes, start codons were retained if they had non-zero occupancy in at least 3 of the replicates in either condition in each dataset, and at least 2 non-zero occupancies in all wt groups and the eIF3c$^{KD}$ or triple condition. Start codon occupancies were then averaged across all samples to obtain one initiation occupancy per start codon per condition. Start codons were filtered for an average initiation occupancy greater than the pseudocount value used in wt and eIF3$^{KD}$ or triple conditions.

RNA correction

To correct for transcript-level changes in RNA abundance, the $log_2$ fold-change in RPFs was regressed against the $log_2$ fold-change in RNA TPM (derived from RNA-seq data). The residuals from this regression were taken as the RNA-corrected $log_2$ fold-change in ribosome occupancy. A similar method was applied to the $log_2$ wt initiation and $log_2$ triple or eIF3c$^{KD}$ initiation values. The resulting residuals were taken as the translation initiation (TI) or fold change (TI_FC) at each start codon. Start codons without corresponding RNA data were removed from the datasets. The final, normalized datasets are provided in the Supplementary Excel File 3.

Modelling

We used Weka 3.9.7 was used to compare features against various models. The M5Rules method, which generates a set of conditions for the model, was used. The following attributes were tested: $log_2$ wt initiation, $log_2$ relative position from the 5'UTR, local GC content (obtained from the 10 downstream and upstream bases from the start codon), start codon group, Kozak score and 5'UTR length. The Kozak score was calculated by adapting scripts from TIS-Predictor into R. A correlation matrix for the features was plotted, and none of the variables were found to have a Pearson correlation > 0.8.

The start codon type was grouped into the following groups due to imbalance in abundance and similar coefficients observed in testing models. AAG and AGG start codons were grouped together, and AUC, AUU and AUA start codons grouped together. All other start codons were used on their own. 10-fold cross validation was used with default Weka settings.

Similar modelling was performed to predict wt TI using the various sequence-based features mentioned above.

Intersection with other data

Genes producing alternative N-terminal isoforms were taken from Supplementary Table S1 (file mmc2.xlsx), sheet "DeepLoc2" of Ly et al.[11], comprising 2,371 genes, each annotated with two or more in-frame N-terminal isoforms (annotated, N-terminally extended and/or truncated).

Enrichment of the isoform set within each query set was assessed by the two-sided hypergeometric test. Expected overlap was calculated as n·K/N and fold enrichment as k/expected where k is the observed overlap, n the query size, K = 2,371 and N the background size.

Genes were also intersected with the Mitocarta 3 data as described above. The entire, integrated data was then screened for examples as per the criteria discussed in the text.

**Other Bioinformatics Methods**

All statistical analyses and plots generation were carried out in GraphPad Prism 9. The statistical method used for each tested hypothesis is provided in the figure legends. All custom computer codes and scripts that were written for this project are publicly available on https://github.com/lml9331/eIF3c. Example genes were selected as described and visualized in R.

**High-resolution respirometry**

The semi-confluent cells (70-80%) were harvested by trypsin (0.025%, 10 minutes), which was subsequently inactivated by the addition of DMEM medium with 10 % fetal bovine serum. Number of cells were counted by Corning Automated Cell Counter. Cells were washed twice with cold PBS and the protein amount was determined by Bradford protein assay (Bio-Rad, 5000006).

Mitochondrial respiration was measured at 37 °C as described[12] using Oxygraph-2k (Oroboros Instruments GmbH). The respiration was measured both in substrates-saturated conditions in permeabilized cells and in intact, non-permeabilized cells. To measure the respiration in permeabilized cells, the cells (0.4 mg of protein) were resuspended in MiR05 medium (0.5 mM EGTA, 3 mM MgCl2, 60 mM lactobionic acid, 20 mM taurine, 10 mM KH2PO4, 20 mM HEPES, 110 mM d-sucrose, and 1 g/l of bovine serum albumin, pH 7.1), and digitonin (0.05 g/g protein) was used to permeabilize the plasma membrane. The following substrates and inhibitors were used for the measurement: 2 mM malate, 10 mM pyruvate, 1 mM ADP, 5 µM cytochrome c, 10 mM glutamate, 10 mM succinate, 10 mM glycerol 3-phosphate, 0.5 µM oligomycin, 0.2 to 1 µM FCCP, 0.5 µM rotenone, 10 mM malonate, 0.25 µM antimycin A, 2 mM ascorbate, 1 mM TMPD, and 0.5 mM KCN. The residual non-mitochondrial oxygen consumption after addition of antimycin A was subtracted as a background correction. For data reporting, two respiratory states were chosen: (1) OXPHOS (oxidative phosphorylation system) - actively phosphorylating cells fully saturated with substrates (pyruvate, glutamate, malate, succinate, and glycerol 3-phosphate) and ADP, (2) ETS (Electron Transfer System) - maximal respiratory capacity of the ETS (fully saturated with substrates, after FCCP ionophore uncoupling). To measure the respiration in non-permeabilized cells, the cells were resuspended in DMEM (Merck, D5030, pH 7.4, 37°C) supplemented with 0.2% (w/w) BSA, 2 mM glutamine and 1 mM pyruvate. For the measurements the following substrate and inhibitors were used: 10 mM glucose (10 mM), 0.5 µM oligomycin, 0.5 to 1.25 µM FCCP, 0.5 µM rotenone, and 0.25 µM antimycin A. The residual non-mitochondrial oxygen consumption after addition of antimycin A was subtracted as a background correction. For data reporting five respiratory states were chosen: (1) routine - basal respiration of the intact cells, (2) routine + GLC - basal respiration after addition of glucose, (3) leak respiration – respiration after addition of oligomycin that inhibits complex V (ATP synthase) (4) ETS - maximal respiratory capacity of the electron transfer system after FCCP ionophore uncoupling, (5) Crabtree effect – the difference between routine respiration and routine respiration + GLC. Data were acquired and analyzed using Datlab 7.4 software (Oroboros Instruments GmbH) and the oxygen consumption was expressed in pmol $O_2$/s/$10^6$ cells.

**Intact mitochondrial respiration and glycolysis**

Seahorse Extracellular Flux (XF) Analyzer (Agilent Technologies, USA) was used to assess intact mitochondrial respiration and glycolytic rate in parallel, as described in[13]. Cells were seeded in pentaplicates (2.5 - 4 × $10^4$) on poly-L-lysine-coated wells of a 24-well measuring plate (Agilent Technologies, USA). After overnight incubation under standard conditions, cells were washed with 1 ml of DMEM (Merck, D5030, pH 7.4, 37°C) supplemented with 0.2% (w/w) BSA, 2 mM glutamine and 1 mM pyruvate and incubated in 500 µL of the media for 30 min at 37°C without $CO_2$. Basal metabolic rate and rates after subsequent additions of the following substrates and inhibitors were measured: oligomycin (1 µM), Bam15 (5 µM), and a mixture of rotenone (1 µM), antimycin A (1 µg/ml), 2-deoxyglucose (100 mM) and Hoechst 33342 (5 µg/ml) to detect nuclei for subsequent counting of cells. Images of whole wells were captured using a Cytation 3 Cell Imaging Reader (BioTek, USA) and the number of the cells was analyzed using the Gen5 software (BioTek). The oxygen consumption (OCR) and extracellular acidification (ECAR) rates were normalized to the cell count and expressed as pmol/min/$10^3$ cells and mpH/min/$10^3$ cells, respectively. Residual OCR and ECAR measured after the addition of 2-deoxyglucose, rotenone, and antimycin A were subtracted as background.

**Statistical analysis**

All statistical analyses and plots generation were carried out in GraphPad Prism 9. The statistical method used for each experiment is described in the figure legends.

#### SUPPLEMENTARY TEXT

##### Translatome-wide consequences of the relaxed stringency of AUG recognition

To determine what consequences the inability of eIF5 to associate with eIF3 has on the entire translatome, we subjected eIF3c$^{KD}$ and our triple mutant cells to ribosome profiling––ribo-seq analysis combined with RNA-seq. Analysis of obtained data revealed enrichment of footprints in coding sequences as well as the expected triplet periodicity (Figure S4A and B, see Methods for details). Both the eIF3c$^{KD}$ and the triple mutant had fewer ribosomes bound to the coding region and UTRs, suggesting reduced initiation rates, as expected (Figs. 2B and 3A). The effect was stronger for the eIF3c$^{KD}$.

We first examined differential translation efficiency by calculating translation efficiency (TE) from ribosome footprint (FP) and mRNA sequencing data using DESeq2[10], where TE reflects ribosome footprint abundance along the entire translon normalized by RNA levels. The obtained reads were aligned to the human genome and transcriptome from Ensembl (GRCh38.p13) using the STAR aligner[9]. Transcripts with adjusted p-values (padj) < 0.05 were considered significantly altered (after correction for multiple testing using the Benjamini-Hochberg method) and are referred to as differential translation efficiency genes (DTEGs)[14]. For each library, the corresponding TE, FP, and mRNA results are given in the Supplementary Excel File 2.

In the case of the triple mutant (3 wt and 3 mutant datasets collected for HEK293T cells), the unfiltered dataset contained 18,011 genes in total (Supplementary Excel File 2). Principal component analysis (PCA) showed clear separation of the triple and wt samples, while biological replicates clustered closely within each group (Figure S5A). In addition, FP and mRNA samples formed distinct clusters, indicating good reproducibility and clear separation of the sample groups. The triple mutant library contained 557 significant DTEGs, including 385 upregulated and 172 downregulated genes. The top 10 upregulated hits include VCL, MTR, IBTK, and RICTOR, while the top 10 downregulated hits include ribosomal proteins, such as RPS2, RPL18A, and RPS11 (Figure S5B).

Gene Ontology (GO) enrichment analysis[15] of the upregulated DTEGs identified mainly signaling- and phosphorylation-associated categories (Figure S5C). In detail, significant DTEGs were selected from the full TE table and separated using padj < 0.05 and then split into upregulated and downregulated groups by the sign of the $\log_2$ fold change. Enrichr terms were ranked by Enrichr adjusted p-value, and terms with adjusted p-value < 0.05 were considered significant. Kyoto Encyclopedia of Genes and Genomes (KEGG) analysis[16] identified signaling pathways, including longevity regulating pathway, hedgehog signaling pathway, phosphatidylinositol signaling system, focal adhesion, and AMPK/FOXO signaling (Figure S5C). In contrast, the downregulated DTEGs were strongly enriched for categories related to translation and ribosome, including cytoplasmic translation, translation, ribosome biogenesis, rRNA metabolic process, ribosomal subunits, and the KEGG term ribosome (Figure S5C). These terms are consistent with the observed reduced ratio of polysomes to monosomes, reduced total protein content, and cell size (Figure S2E and F).

Notably, the downregulated DTEG fraction also showed a mitochondrial signature, with oxidative phosphorylation appearing among the KEGG terms in this group (Figure S5C). To assess this signature directly, significant DTEGs were intersected with Human MitoCarta 3.0 database (Supplementary Excel File 2). MitoCarta genes were significantly enriched among downregulated DTEGs (29 genes, $p = 0.00000079$, hypergeometric test), but not among upregulated DTEGs (7 genes, $p > 0.05$), indicating that the mitochondrial signature is mainly driven by the downregulated fraction. The downregulated MitoCarta genes included mtDNA-encoded OXPHOS genes, mitochondrial ribosomal proteins, and nuclear-encoded OXPHOS components and mitochondrial translation factors (Figure S5D).

In the case of eIF3c$^{KD}$, the unfiltered dataset contained 17,101 genes in total from 6 wt and 4 knockdown experiments in HeLa cells (Supplementary Excel File 2). PCA analysis showed clear separation of eIF3c$^{KD}$ and wt HeLa samples, while biological replicates clustered closely within each group (Figure S6A). The eIF3c$^{KD}$ library contained 2,536 significant DTEGs, including 1,367 upregulated and 1,169 downregulated genes. The top 10 upregulated hits included VCL encoding vinculin (a crucial cytoskeletal protein involved in cell-cell and cell-matrix junctions[17], as in the case of the triple, and MARS1, FLNB, and SPTBN1, whereas the most severely downregulated hits included CSDE1, CREBBP, GIT1, and DVL3 (Figure S6B). As expected, the eIF3c$^{KD}$ exhibited a greater number of significant changes in TE, FP, and mRNA compared with the triple mutant (Supplementary Excel File 2). This reflects its deleterious effect on eIF3 integrity and, consequently, its much stronger impact on the functionality of the translation initiation mechanism.

Not surprisingly, the eIF3c$^{KD}$ library showed a larger number of significant DTEGs than the triple mutant. In contrast to the triple mutant, upregulated DTEGs in the eIF3c$^{KD}$ were enriched for translation- and ribosome-associated categories, whereas downregulated DTEGs were enriched for kinase-, phosphorylation-, and cell-cycle-related processes (Figure S6C). Together, these results indicate that altered eIF3 function affects related functional categories, but that the outcome differs depending on whether eIF3 is selectively impaired in eIF5 binding or more broadly affected by eIF3c depletion.

Comparative analysis with Human MitoCarta3.0 similarly pointed to genes associated with mitochondrial function (Figure S6D). In particular, we identified 244 mitochondria-associated genes, 159 upregulated and 85 downregulated (Supplementary Excel File 2). Although the eIF3c$^{KD}$ library exhibited a broader range of mitochondrial-associated traits than the triple mutant, we focused the detailed mitochondria-related analysis on the triple mutant. This was because eIF3c depletion strongly affects the eIF3 integrity and this general translation, whereas the triple mutant has a very specific impact on translation initiation stringency and efficiency. This should allow for a more direct interpretation of the mitochondrial phenotype (this analysis is shown below).

#### FIGURE S LEGENDS

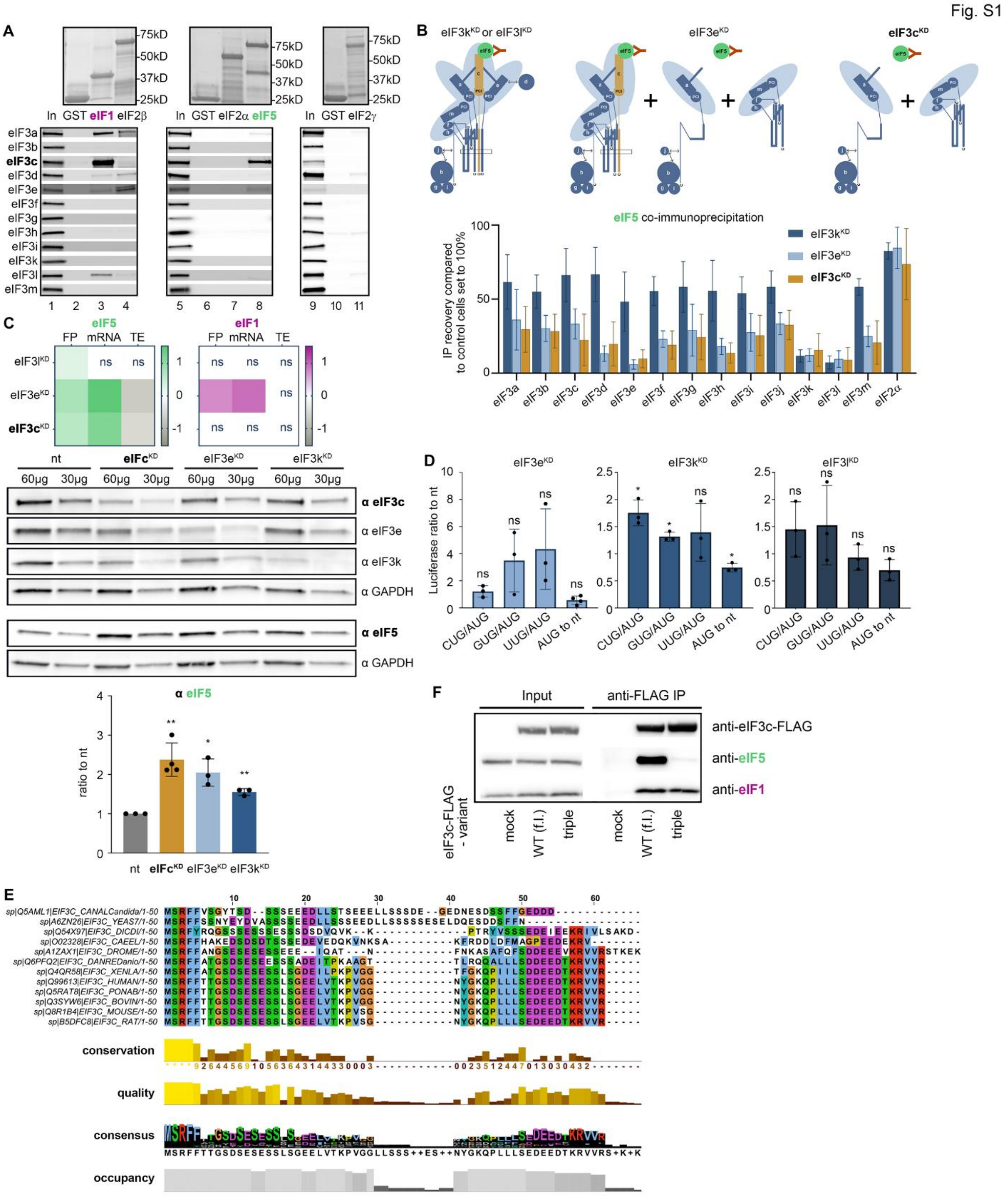


**Figure S1. Protein-protein and knock down analysis of eIF3 interactions with its associated eIFs.**

(**A**) GST pull-down analysis of interactions between individual radiolabeled human eIF3 subunits and the indicated initiation factors fused to GST. GST alone served as negative control. Lane 1 contains 50% of the input amounts of *in vitro* translated proteins added to each reaction.

(**B**) The top panel shows schematics of the eIF3 subcomplexes generated by individual knock-downs of the eIF3k, eIF3l, eIF3e and eIF3c subunits, as determined in[18,19]. The eIF5-binding capacity of these eIF3 subcomplexes and the eIF5 co-immunoprecipitation strategy are also depicted. The bottom plot shows the results of eIF5 co-immunoprecipitation from HeLa cell lysates with $eIF3k^{KD}$, $eIF3e^{KD}$ and $eIF3c^{KD}$. Immunoprecipitated eIF3 subunits, along with eIF2α and eIF5, were detected by western blotting, quantified using QuantityOne, and the signals were normalized to eIF5 (bait). The resulting values obtained with the control nt cells were set to 100%, and those obtained with individual KDs were expressed relative to the nt. Plot summarizes results from n = 4–6 biological replicates. For details see Source Data Figure S1.

(**C**) Effect of the eIF3 subunit KDs on eIF5 and eIF1 endogenous expression. The upper panels show ribosome footprint (FP), mRNA, and TE changes for eIF5 and eIF1 in the indicated knockdowns. The middle panels show representative western blots of indicated proteins in non-targeting (nt) control and eIF3 KD cells. GAPDH was used as a loading control. The bottom panel shows quantification of western blotting. It was performed in multiple technical replicates per each of the 3-4 biological replicates ($n \geq 3$); for the statistical analysis, the technical replicate with the most uniform loading from each biological replicate was selected (dots). The plot depicts the mean ratio ± S.D. of eIF5 protein levels in individual KDs relative to nt control, which was set to 1. One sample t test was used for statistical evaluation (* $p \leq 0.05$, ** $p \leq 0.01$).

(**D**) The impact of selected eIF3 KDs on stringency of the TIS selection. HeLa cells were transfected with the non-targeting (nt) or eIF3e, eIF3k and eIF3l siRNAs, and 48 h later, the second round of transfection with individual reporters (AUG or CUG, GUG, UUG, AAG) was carried out. Luciferase activities were measured in lysate 24 h after the second transfection. mRNA levels of each reporter in the same lysate were measured by qPCR and used for normalization. Plot shows the mean ratio ± S.D. of the near-cognate initiation rates relative to those of AUG; this ratio in the nt sample was set to 1. Dots represent results from individual biological replicates ($n \geq 3$). One sample t test was used for statistical evaluation (* $p \leq 0.05$, ** $p \leq 0.01$, *** $p \leq 0.001$, **** $p \leq 0.0001$, ns = not significant.

(**E**) Multiple-sequence alignment of the extreme N-terminal region of eIF3c from representative eukaryotic species. Conservation, quality, consensus, and occupancy are given below the alignment.

(**F**) Competition between endogenous eIF3 and wt or mutant eIF3c-FLAG protein for eIF5 *in vivo*. The C-terminally FLAG-tagged full-length eIF3c was overexpressed in HEK293T cells and immunoprecipitated using anti-FLAG beads. The levels of eIF5, eIF1, and eIF3c-FLAG were detected by Western blotting.

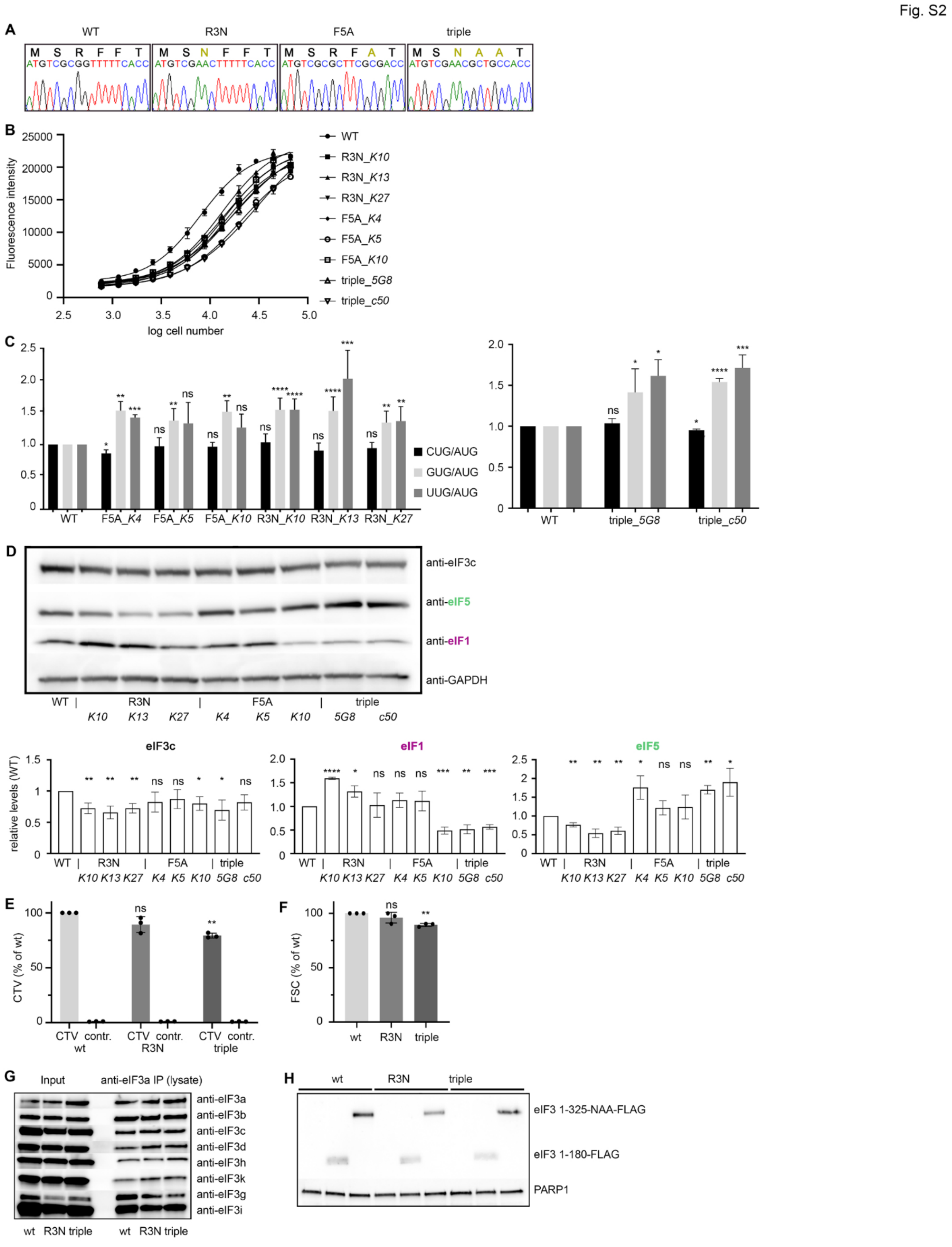


**Figure S2. Characterization of the eIF3c knock-in clones and their effects on growth, TIS selection, and protein levels of eIF1 and eIF5.**

(**A**) Validation of the eIF3c-NTD knock-in mutations by Sanger sequencing. Representative DNA sequence chromatograms of the *EIF3C* N-terminal coding region in wt, R3N, F5A, and triple mutant HEK293T cells. The introduced nucleotide substitutions and the corresponding amino acid changes are indicated above the chromatograms.

(**B**) Resazurin reduction assay growth curves. An equal number of wt cells and cells from all available knock-in clones was serially diluted in culture medium at a 2:1 ratio into a 96-well plate and incubated for 48 hours. Resazurin was then added to each well for 3 hours, and fluorescence was measured. Fluorescence intensity is plotted against the logarithm of the initial cell count in each well, and the curves were fitted using a four-parameter nonlinear regression model (4PL) (GraphPad Prism).

(**C**) The eIF3c knock-ins impair stringency of TIS selection. *Left panel*: wt and 3 clones of F5A and R3N mutant cells each were transfected with AUG- or near-cognate codon-initiated firefly luciferase reporters. Firefly luciferase mRNA levels determined by qPCR were used for normalization. Plot shows the mean ratio ± S.D. of the near-cognate initiation rates relative to those of AUG ($n \geq 4$); this ratio in the wt sample was set to 1. Statistical significance was determined by mixed-effects model (REML) with Dunnett's multiple-comparisons test (* $p \leq 0.05$, ** $p \leq 0.01$, *** $p \leq 0.001$, **** $p \leq 0.0001$). *Right panel*: wt and 2 clones of triple mutant cells were transfected with AUG- or near-cognate codon-initiated firefly luciferase reporters. Renilla luciferase reporter was used for transfection efficiency normalization. Plot shows the mean ratio ± S.D. of the near-cognate initiation rates relative to those of AUG ($n \geq 4$); this ratio in the wt sample was set to 1. Statistical significance was determined by paired t test (* $p \leq 0.05$, ** $p \leq 0.01$, *** $p \leq 0.001$, **** $p \leq 0.0001$).

(**D**) Western blot analysis of eIF1, eIF5, and eIF3c protein levels in all available knock-in clones. Densitometric analysis was performed by GelAnalyzer and the signal intensities were normalized to GAPDH. Plot shows the mean ± S.D. of protein levels of indicated eIFs in knock-in mutants relative to those in wt ($n = 4$); which were set to 1. Statistical significance was determined by One sample t test (* $p \leq 0.05$, ** $p \leq 0.01$, *** $p \leq 0.001$, **** $p \leq 0.0001$).

(**E**) CellTrace Violet signal in wt and selected eIF3c knock-ins. wt, R3N, and triple mutant HEK293T cells were stained with CellTrace Violet (CTV; 0.3 μM) or DMSO-treated (mock control) and measured by flow cytometry. CTV staining in mutant cells is expressed as percentage of the wt signal. Control samples indicate background fluorescence. Data from independent replicates are presented as mean ± S.D. ($n = 3$). Statistical significance was determined by One sample t test (** $p \leq 0.01$; ns, not significant).

(**F**) Quantification of relative cell size of wt, R3N, and triple mutant HEK293T cells by flow cytometry. Relative cell size was estimated from the forward scatter (FSC) parameter. Data from independent replicates are presented as mean ± S.D. ($n = 3$). Statistical significance was determined by One sample t test (** $p \leq 0.01$; ns, not significant).

(**G**) Co-immunoprecipitation of the eIF3 complex demonstrates its intact integrity. Formaldehyde-crosslinked cell lysates of wt, R3N, and triple mutant cells were subjected to immunoprecipitation with anti-eIF3a antibodies followed by western blotting.

(**H**) Related to Figure 2H; the levels of C-terminally FLAG-tagged eIF3c fragments: 1-180 aa representing "eIF5 decoy", and 1-325 aa representing "eIF1 decoy", which were overexpressed in wt, R3N, and triple cells, were monitored by western blotting.

Fig. S3

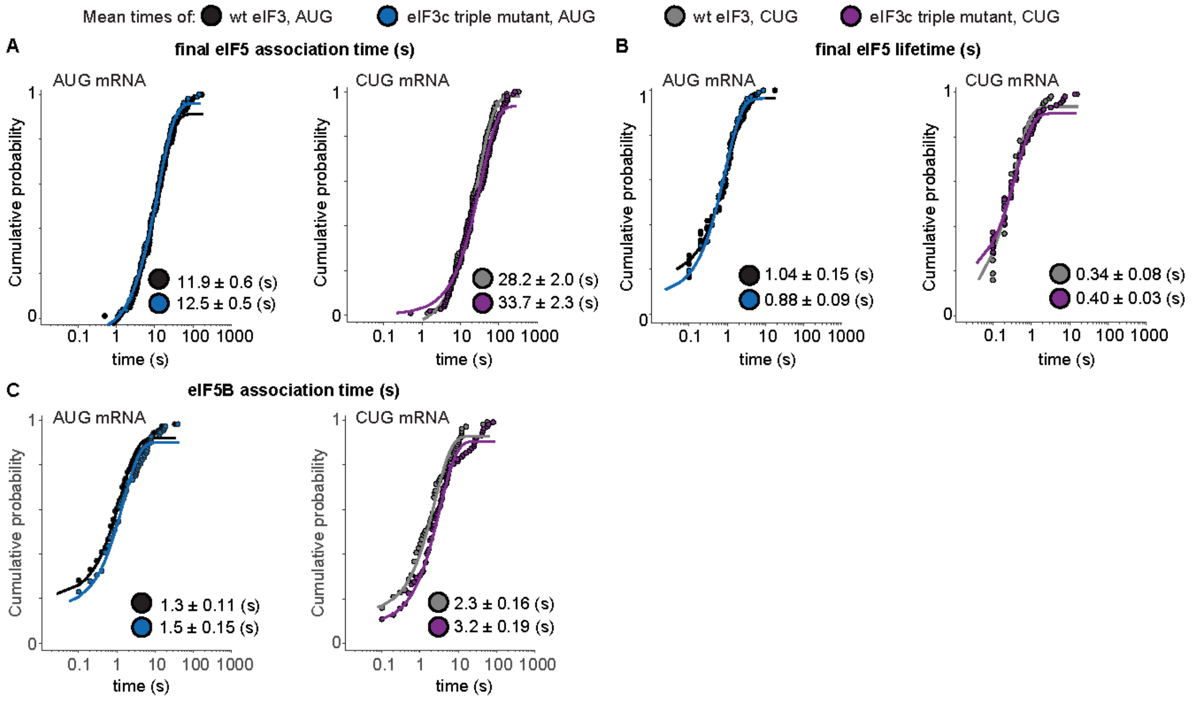


**Figure S3. eIF5 binding to initiation complexes is slower and less stable in the eIF3c triple mutant.**

(**A** – **C**) Cumulative probability plots of the indicated parameters (defined in Figure 4C). Lines represent fit to exponential functions, which were used to derive rates, residency times (lifetimes), and mean times (annotated on plots) for eIF5 and eIF5B with wt eIF3, AUG mRNA (black), eIF3c triple mutant, AUG mRNA (blue), wt eIF3, CUG mRNA (gray), and eIF3c triple mutant, CUG mRNA (purple). See Supplementary Excel File 1 for all rates, rate constants, and the number of complexes and binding events analyzed in each experiment.

Fig. S4

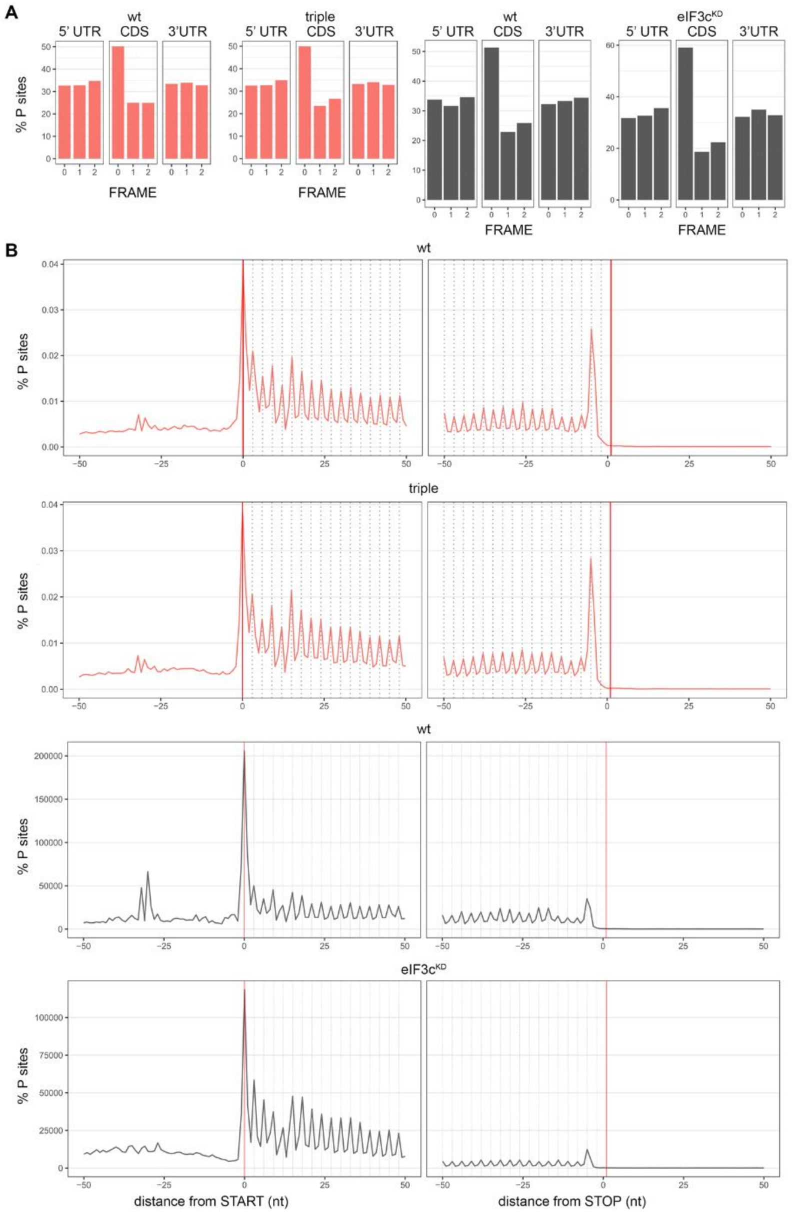


**Figure S4. Quality control of ribosome-profiling libraries confirms expected reading-frame periodicity and metagene profiles.**

(**A**) Frame distribution of ribosome footprint (FP) reads in wt and eIF3c mutant cells. Percentage of FP reads mapping to the three possible reading frames in the 5′ UTR, CDS, and 3′ UTR regions of wt and triple mutant (HEK293T cells), wt and eIF3c$^{KD}$ (HeLa cells) libraries. All libraries show the expected enrichment of reads in the main coding frame within the CDS, consistent with the 3-nt periodicity of translating ribosomes.

(**B**) The P-site metaprofiles of ribosome footprint libraries. P-site frequencies are shown relative to annotated start codons and stop codons in the triple mutant and the corresponding wt, and eIF3c$^{KD}$ and the corresponding wt libraries. All libraries display a strong peak at the start codon and the expected 3-nt periodicity across coding regions, consistent with high-quality ribosome profiling data.

Fig. S5

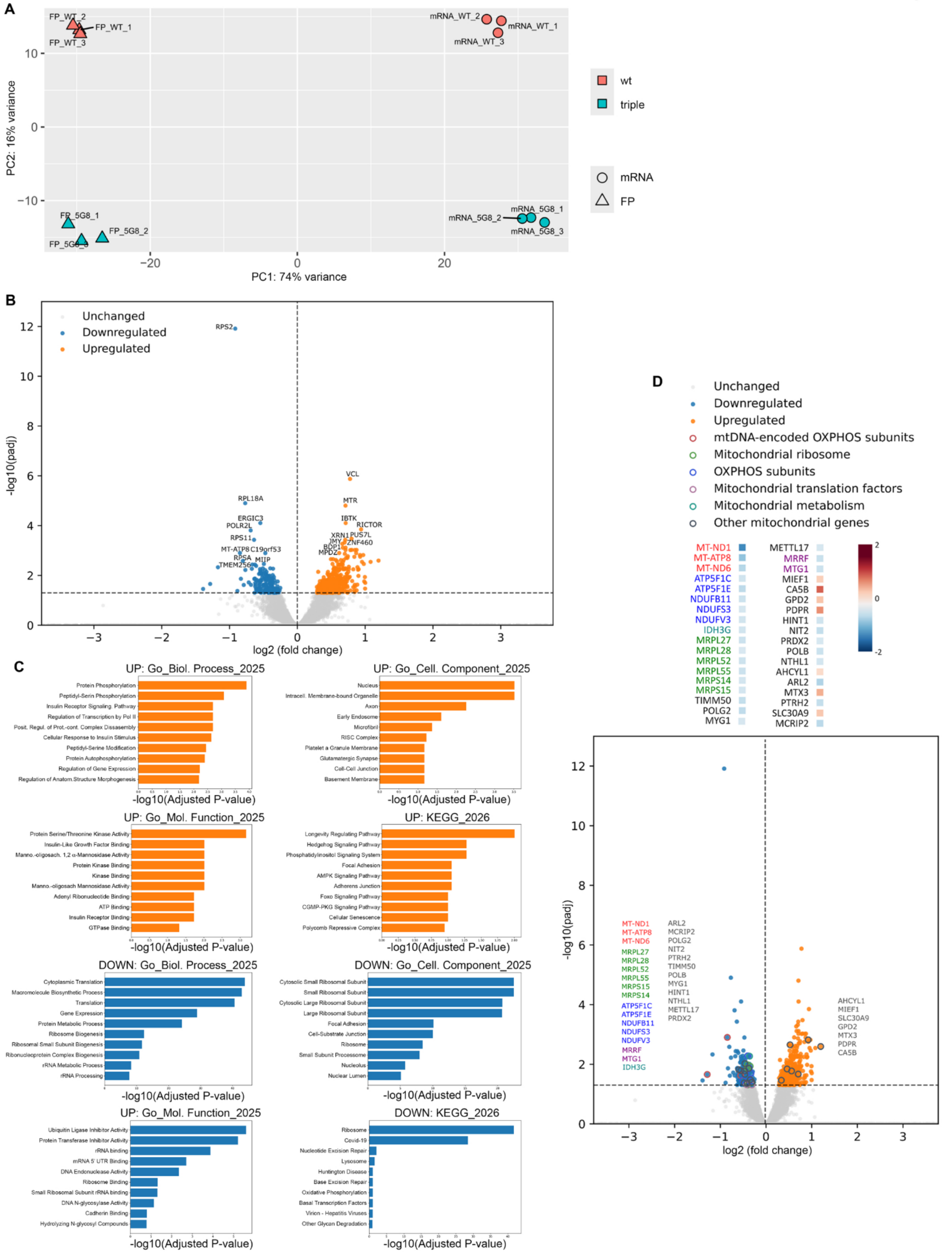
A
PC1: 74% variance
PC2: 16% variance
FP_WT_1
FP_WT_2
FP_WT_3
mRNA_WT_1
mRNA_WT_2
mRNA_WT_3
FP_5G8_1
FP_5G8_2
mRNA_5G8_1
mRNA_5G8_2
mRNA_5G8_3
wt
triple
mRNA
FP
B
Unchanged
Downregulated
Upregulated
-log10(padj)
log2 (fold change)
RPS2
RPL18A
ERGIC3
POLR2L
RPS11
MT-ATP8
C19orf53
RPSA
MIIP
TMEM256
VCL
MTR
IBTK
RICTOR
XRN1
PUS7L
ZNF460
MPDZ
C
UP: Go_Biol. Process_2025
Protein Phosphorylation
Peptidyl-Serin Phosphorylation
Insulin Receptor Signaling Pathway
Regulation of Transcription by Pol II
Posit. Regul. of Prot.-cont. Complex Dissasembly
Cellular Response to Insulin Stimulus
Peptidyl-Serine Modification
Protein Autophosphorylation
Regulation of Gene Expression
Regulation of Anatom.Structure Morphogenesis
-log10(Adjusted P-value)
UP: Go_Cell. Component_2025
Nucleus
Intracell. Membrane-bound Organelle
Axon
Early Endosome
Microfibril
RISC Complex
Platelet a Granule Membrane
Glutamatergic Synapse
Cell-Cell Junction
Basement Membrane
-log10(Adjusted P-value)
UP: Go_Mol. Function_2025
Protein Serine/Threonine Kinase Activity
Insulin-Like Growth Factor Binding
Manno.-oligosach. 1,2 α-Mannosidase Activity
Protein Kinase Binding
Kinase Binding
Manno.-oligosach Mannosidase Activity
Adenyl Ribonucleotide Binding
ATP Binding
Insulin Receptor Binding
GTPase Binding
-log10(Adjusted P-value)
UP: KEGG_2026
Longevity Regulating Pathway
Hedgehog Signaling Pathway
Phosphatidylinositol Signaling System
Focal Adhesion
AMPK Signaling Pathway
Adherens Junction
Foxo Signaling Pathway
CGMP-PKG Signaling Pathway
Cellular Senescence
Polycomb Repressive Complex
-log10(Adjusted P-value)
DOWN: Go_Biol. Process_2025
Cytoplasmic Translation
Macromolecule Biosynthetic Process
Translation
Gene Expression
Protein Metabolic Process
Ribosome Biogenesis
Ribosomal Small Subunit Biogenesis
Ribonucleoprotein Complex Biogenesis
rRNA Metabolic Process
rRNA Processing
-log10(Adjusted P-value)
DOWN: Go_Cell. Component_2025
Cytosolic Small Ribosomal Subunit
Small Ribosomal Subunit
Cytosolic Large Ribosomal Subunit
Large Ribosomal Subunit
Focal Adhesion
Cell-Substrate Junction
Ribosome
Small Subunit Processome
Nucleolus
Nuclear Lumen
-log10(Adjusted P-value)
UP: Go_Mol. Function_2025
Ubiquitin Ligase Inhibitor Activity
Protein Transferase Inhibitor Activity
rRNA binding
mRNA 5' UTR Binding
DNA Endonuclease Activity
Ribosome Binding
Small Ribosomal Subunit rRNA binding
DNA N-glycosylase Activity
Cadherin Binding
Hydrolyzing N-glycosyl Compounds
-log10(Adjusted P-value)
DOWN: KEGG_2026
Ribosome
Covid-19
Nucleotide Excision Repair
Lysosome
Huntington Disease
Base Excision Repair
Oxidative Phosphorylation
Basal Transcription Factors
Virion - Hepatitis Viruses
Other Glycan Degradation
-log10(Adjusted P-value)
D
Unchanged
Downregulated
Upregulated
mtDNA-encoded OXPHOS subunits
Mitochondrial ribosome
OXPHOS subunits
Mitochondrial translation factors
Mitochondrial metabolism
Other mitochondrial genes
MT-ND1
MT-ATP8
MT-ND6
ATP5F1C
ATP5F1E
NDUFB11
NDUFS3
NDUFV3
IDH3G
MRPL27
MRPL28
MRPL52
MRPL55
MRPS14
MRPS15
TIMM50
POLG2
MYG1
METTL17
MRRF
MTG1
MIEF1
CA5B
GPD2
PDPR
HINT1
NIT2
PRDX2
POLB
NTHL1
AHCYL1
ARL2
MTX3
PTRH2
SLC30A9
MCRIP2
-log10(padj)
log2 (fold change)

**Figure S5. Translatome-wide effects of the eIF3c triple mutation reveal altered TE and mitochondrial-gene enrichment**

(**A**) Principal component analysis (PCA) of ribosome footprint (FP) and mRNA-seq samples from triple mutant (5G8) and wt HEK293T cells. Biological replicates cluster closely within each condition.

(**B**) Volcano plot of differential TE in the triple mutant compared to wt. Significantly upregulated DTEGs are shown in orange, significantly downregulated DTEGs in blue, and non-significant genes in grey. The x-axis shows $\log_2$ (fold change) and the y-axis shows -$\log_{10}$ (adjusted p-value). The 10 most significant upregulated and downregulated genes are labeled. The analysis was performed using Ensembl gene identifiers (ENSG).

(**C**) Enrichr-based enrichment analysis of upregulated (orange) and downregulated (blue) TE genes in the triple mutant compared to wt. The x-axis shows -$\log_{10}$ (adjusted p-value).

(**D**) Volcano plot of differential TE in the triple mutant compared to wt with Human MitoCarta3.0 overlap highlighted. The x-axis shows $\log_2$ (fold change) and the y-axis shows -$\log_{10}$ (adjusted p-value). Significantly upregulated genes are shown in orange, significantly downregulated genes in blue, and non-significant genes in grey. Colored outlined circles mark the 36 significant mitochondria-associated TE genes identified by overlap with Human MitoCarta3.0, and gene labels are color-coded by category. The highlighted genes are grouped into six broad categories: mtDNA-encoded OXPHOS subunits (n=3), mitochondrial ribosome (n=6), OXPHOS subunits (n=5), mitochondrial translation factors (n=2), mitochondrial metabolism (n=1), and other mitochondrial genes (n=19).

Fig.S 6

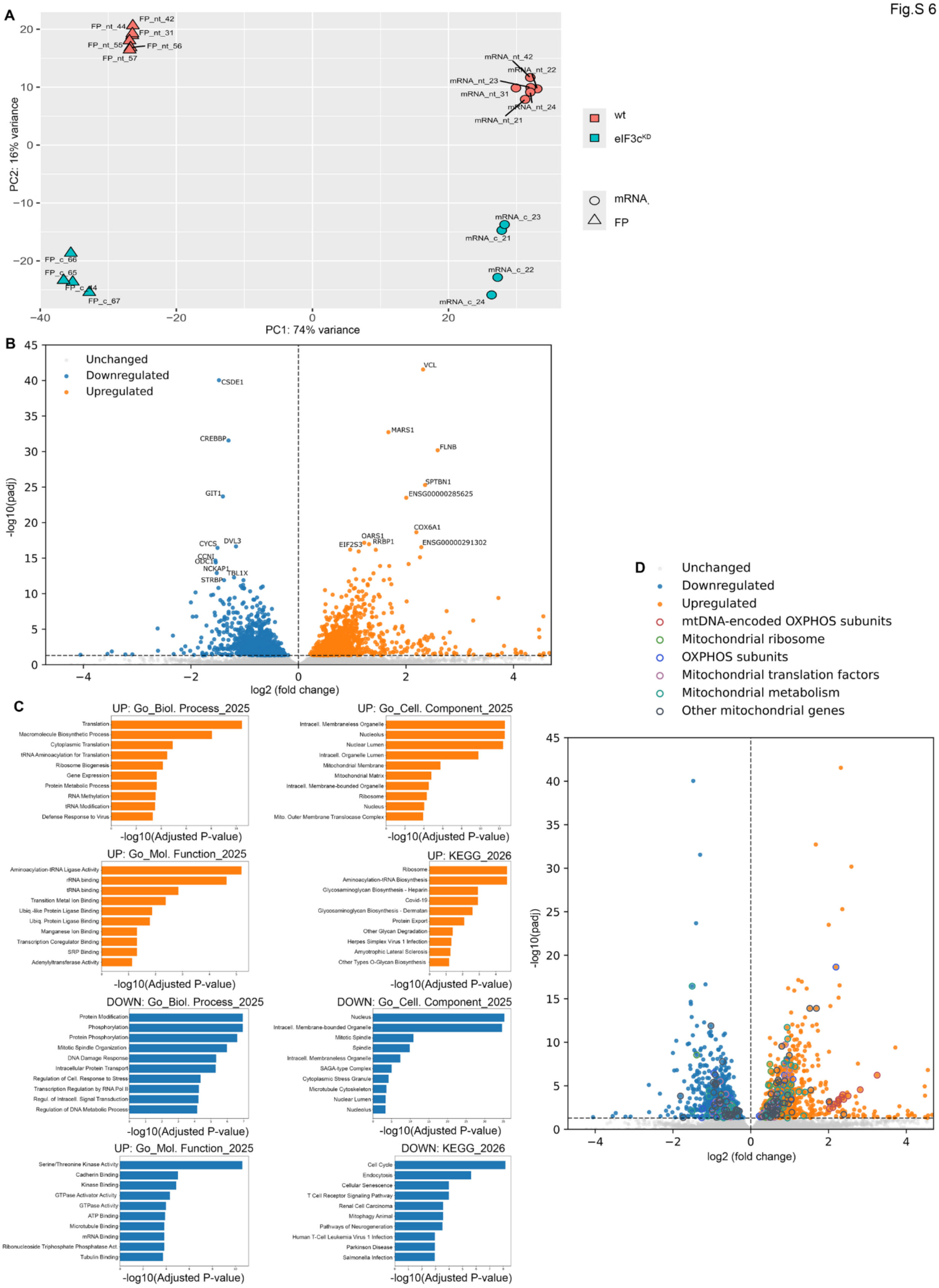
A
PC1: 74% variance
PC2: 16% variance
FP_nt_42
FP_nt_44
FP_nt_31
FP_nt_55
FP_nt_56
FP_nt_57
mRNA_nt_42
mRNA_nt_22
mRNA_nt_23
mRNA_nt_31
mRNA_nt_24
mRNA_nt_21
mRNA_c_23
mRNA_c_21
mRNA_c_22
mRNA_c_24
FP_c_66
FP_c_65
FP_c_44
FP_c_67
wt
eIF3c$^{KD}$
mRNA
FP
B
Unchanged
Downregulated
Upregulated
-log10(padj)
log2 (fold change)
CSDE1
CREBBP
GIT1
CYCS
DVL3
CCNI
ODC1
NCKAP1
TBL1X
STRBP
VCL
MARS1
FLNB
SPTBN1
ENSG00000285625
COX6A1
OARS1
RRBP1
EIF2S3
ENSG00000291302
C
UP: Go_Biol. Process_2025
Translation
Macromolecule Biosynthetic Process
Cytoplasmic Translation
tRNA Aminoacylation for Translation
Ribosome Biogenesis
Gene Expression
Protein Metabolic Process
RNA Methylation
tRNA Modification
Defense Response to Virus
-log10(Adjusted P-value)
UP: Go_Cell. Component_2025
Intracell. Membraneless Organelle
Nucleolus
Nuclear Lumen
Intracell. Organelle Lumen
Mitochondrial Membrane
Mitochondrial Matrix
Intracell. Membrane-bounded Organelle
Ribosome
Nucleus
Mito. Outer Membrane Translocase Complex
-log10(Adjusted P-value)
UP: Go_Mol. Function_2025
Aminoacylation-tRNA Ligase Activity
rRNA binding
tRNA binding
Transition Metal Ion Binding
Ubiq.-like Protein Ligase Binding
Ubiq. Protein Ligase Binding
Manganese Ion Binding
Transcription Coregulator Binding
SRP Binding
Adenylyltransferase Activity
-log10(Adjusted P-value)
UP: KEGG_2026
Ribosome
Aminoacylation-tRNA Biosynthesis
Glycosaminoglycan Biosynthesis - Heparin
Covid-19
Glycosaminoglycan Biosynthesis - Dermatan
Protein Export
Other Glycan Degradation
Herpes Simplex Virus 1 Infection
Amyotrophic Lateral Sclerosis
Other Types O-Glycan Biosynthesis
-log10(Adjusted P-value)
DOWN: Go_Biol. Process_2025
Protein Modification
Phosphorylation
Protein Phosphorylation
Mitotic Spindle Organization
DNA Damage Response
Intracellular Protein Transport
Regulation of Cell. Response to Stress
Transcription Regulation by RNA Pol II
Regul. of Intracell. Signal Transduction
Regulation of DNA Metabolic Process
-log10(Adjusted P-value)
DOWN: Go_Cell. Component_2025
Nucleus
Intracell. Membrane-bounded Organelle
Mitotic Spindle
Spindle
Intracell. Membraneless Organelle
SAGA-type Complex
Cytoplasmic Stress Granule
Microtubule Cytoskeleton
Nuclear Lumen
Nucleolus
-log10(Adjusted P-value)
UP: Go_Mol. Function_2025
Serine/Threonine Kinase Activity
Cadherin Binding
Kinase Binding
GTPase Activator Activity
GTPase Activity
ATP Binding
Microtubule Binding
mRNA Binding
Ribonucleoside Triphosphate Phosphatase Act.
Tubulin Binding
-log10(Adjusted P-value)
DOWN: KEGG_2026
Cell Cycle
Endocytosis
Cellular Senescence
T Cell Receptor Signaling Pathway
Renal Cell Carcinoma
Mitophagy Animal
Pathways of Neurogeneration
Human T-Cell Leukemia Virus 1 Infection
Parkinson Disease
Salmonella Infection
-log10(Adjusted P-value)
D
Unchanged
Downregulated
Upregulated
mtDNA-encoded OXPHOS subunits
Mitochondrial ribosome
OXPHOS subunits
Mitochondrial translation factors
Mitochondrial metabolism
Other mitochondrial genes
-log10(padj)
log2 (fold change)

**Figure S6. The eIF3c$^{KD}$ causes broad translatome-wide changes in TE and enrichment of mitochondrial-associated genes.**

(**A**) Principal component analysis (PCA) of ribosome footprint (FP) and mRNA-seq samples from eIF3c$^{KD}$ and wt HeLa cells. Biological replicates cluster closely within each condition.

(**B**) Volcano plot of differential TE in eIF3c$^{KD}$ compared to wt. Significantly upregulated DTEGs are shown in orange, significantly downregulated DTEGs in blue, and non-significant genes in grey. The x-axis shows $\log_2$ (fold change) and the y-axis shows $-\log_{10}$ (adjusted p-value). The 10 most significant upregulated and downregulated genes are labeled. The analysis was performed using Ensembl gene identifiers (ENSG).

(**C**) Enrichr-based enrichment analysis of upregulated (orange) and downregulated (blue) TE genes in eIF3c$^{KD}$ compared to wt. The x-axis shows $-\log_{10}$ (adjusted p-value).

(**D**) Volcano plot of differential TE in eIF3c$^{KD}$ compared to wt with Human MitoCarta3.0 overlap highlighted. Upregulated genes are shown in orange, downregulated genes in blue, and non-significant genes in grey. Mitochondria-associated TE genes are grouped into six categories: mtDNA-encoded OXPHOS subunits, mitochondrial ribosome, OXPHOS subunits, mitochondrial translation factors, mitochondrial metabolism, and other mitochondrial genes.

Fig.S7

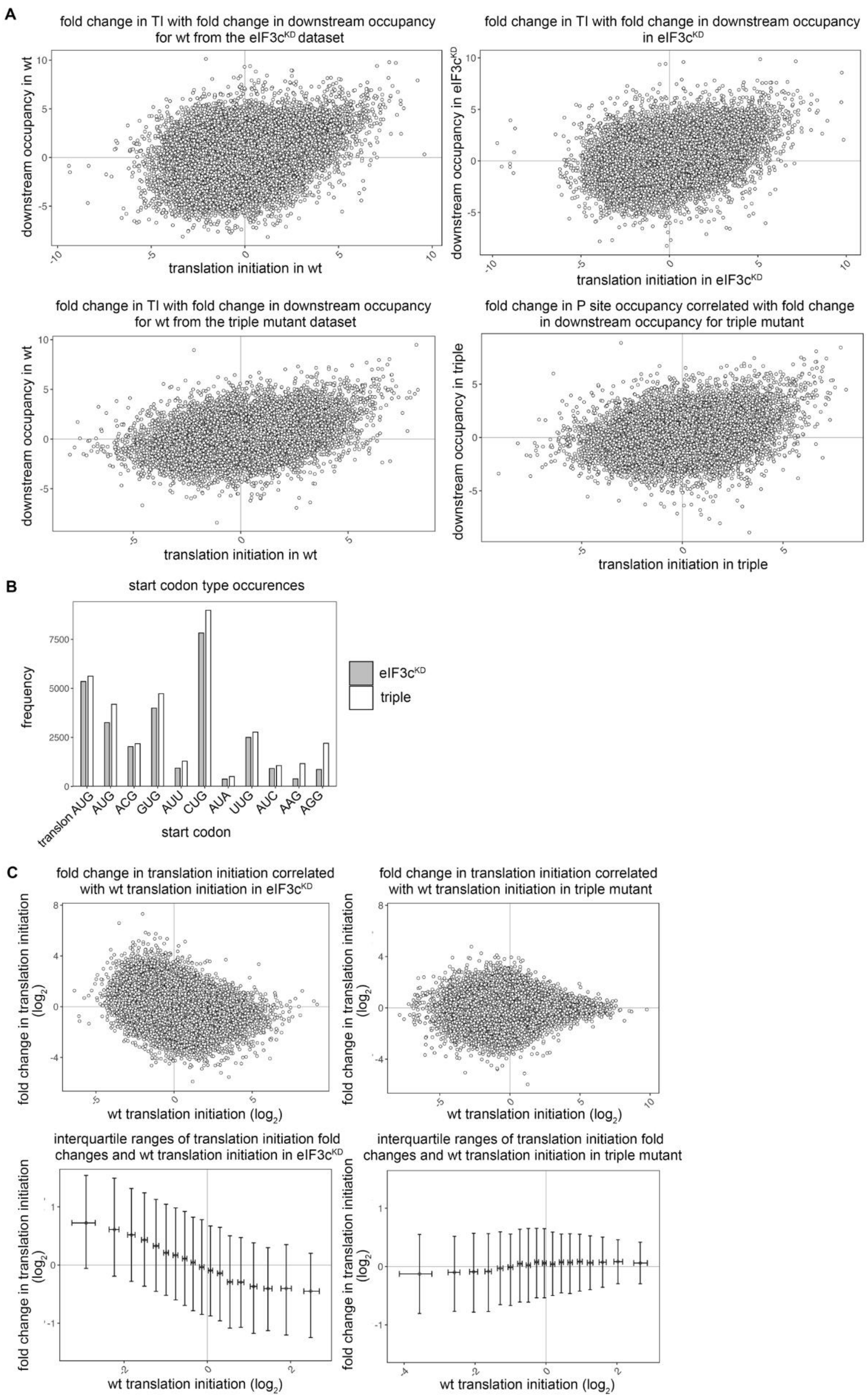
A
fold change in TI with fold change in downstream occupancy for wt from the eIF3c^KD dataset
downstream occupancy in wt
translation initiation in wt
fold change in TI with fold change in downstream occupancy in eIF3c^KD
downstream occupancy in eIF3c^KD
translation initiation in eIF3c^KD
fold change in TI with fold change in downstream occupancy for wt from the triple mutant dataset
downstream occupancy in wt
translation initiation in wt
fold change in P site occupancy correlated with fold change in downstream occupancy for triple mutant
downstream occupancy in triple
translation initiation in triple
B
start codon type occurences
frequency
eIF3c^KD
triple
translon AUG
AUG
ACG
GUG
AUU
CUG
AUA
UUG
AUC
AAG
AGG
start codon
C
fold change in translation initiation correlated with wt translation initiation in eIF3c^KD
fold change in translation initiation correlated with wt translation initiation in triple mutant
fold change in translation initiation (log2)
wt translation initiation (log2)
interquartile ranges of translation initiation fold changes and wt translation initiation in eIF3c^KD
interquartile ranges of translation initiation fold changes and wt translation initiation in triple mutant
fold change in translation initiation (log2)
wt translation initiation (log2)

**Figure S7. Wild-type initiation strength predicts TIS response to the eIF3c-NTD impairment**

(**A**) Translation initiation correlates with downstream occupancy indicating true initiation. The scatter plots on the left show the correlation between translation initiation (TI) (x-axis, wt translation initiation) and the average ribosome occupancy of the next/downstream 2 codons (y-axis, wt) normalized to RNA levels. The correlation was 0.32 and 0.36 (Pearson) for the wt for $eIF3c^{KD}$ and the triple, respectively. The scatter plots on the right show the correlation between translation initiation (TI) in knockdown (x-axis, top) or mutant (x-axis, bottom) conditions against the average occupancy of the next/downstream 2 codons (y- axis, knockdown in top graph, mutant below). The correlation (Pearson) was 0.36 and 0.31 for the $eIF3c^{KD}$ and the triple respectively, suggesting that initiation (reads on start codon) leads to elongation (downstream reads). For all datasets, the 3-nt periodicity of reads downstream of the start codon remained the same (not shown) further supporting elongation and therefore true initiation at the TIS.

(**B**) Frequency of occupied start codons in $eIF3c^{KD}$ and the triple mutant datasets. The bar plots show the number of occupied start codons, in both wt and the impaired condition, for each 5' UTR start codon type and main translon's start codon for each dataset, with $eIF3c^{KD}$ data in grey and triple mutant data in white.

(**C**) Correlation between fold change in translation initiation upon the eIF3c-NTD impairment and the wt translation initiation. In each plot, the x-axis denotes $log_2$ wt translation initiation. On the left, y-axis denotes $log_2$ ratio of $eIF3c^{KD}$ translation initiation / wt translation initiation for $eIF3c^{KD}$. On the right, y-axis denotes $log_2$ ratio of the triple mutant translation initiation / wt translation initiation for the triple mutant. Plots on top show scatter plots of wt translation initiation and the fold-change in translation initiation upon the eIF3c-NTD impairment. Plots on the bottom show median and interquartile ranges of equal sections of the data.

Fig.S8

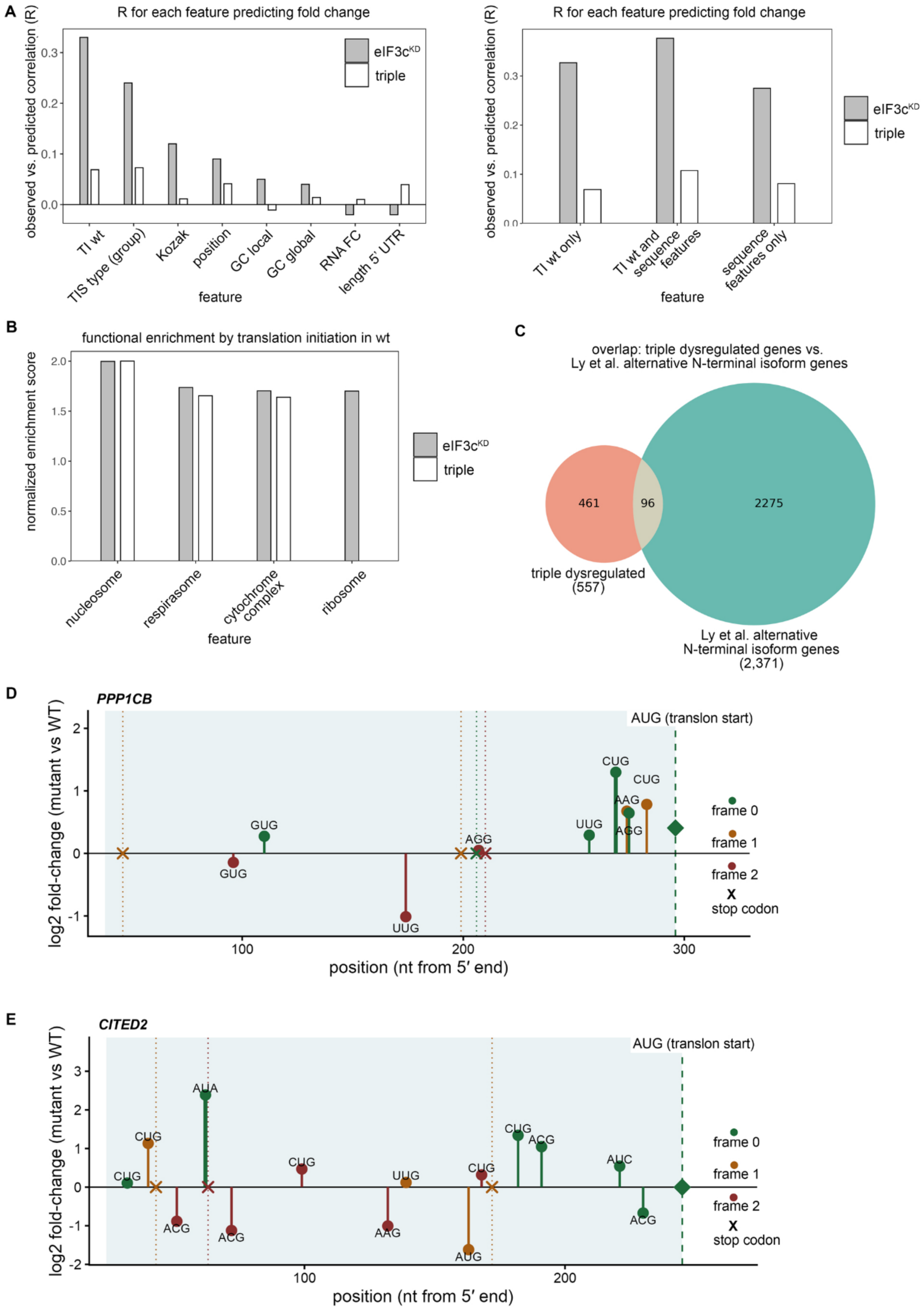

A
R for each feature predicting fold change
observed vs. predicted correlation (R)
0.0
0.1
0.2
0.3
eIF3c^KD
triple
TI wt
TIS type (group)
Kozak
position
GC local
GC global
RNA FC
length 5' UTR
feature
R for each feature predicting fold change
observed vs. predicted correlation (R)
TI wt only
TI wt and sequence features
sequence features only
feature
eIF3c^KD
triple
B
functional enrichment by translation initiation in wt
normalized enrichment score
0.0
0.5
1.0
1.5
2.0
nucleosome
respirasome
cytochrome complex
ribosome
feature
eIF3c^KD
triple
C
overlap: triple dysregulated genes vs.
Ly et al. alternative N-terminal isoform genes
461
96
2275
triple dysregulated
(557)
Ly et al. alternative
N-terminal isoform genes
(2,371)
D
PPP1CB
log2 fold-change (mutant vs WT)
AUG (translon start)
CUG
CUG
AAG
AGG
UUG
GUG
GUG
AGG
UUG
frame 0
frame 1
frame 2
X
stop codon
100
200
300
position (nt from 5′ end)
E
CITED2
log2 fold-change (mutant vs WT)
AUG (translon start)
AUA
CUG
CUG
CUG
UUG
CUG
CUG
ACG
AUC
ACG
ACG
ACG
AAG
AUG
ACG
frame 0
frame 1
frame 2
X
stop codon
100
200
position (nt from 5′ end)

**Figure S8. Correlation between observed and predicted fold change in TI for individual mRNA features and functional enrichment amongst genes rank-ordered by wt TI**.

(**A**) Correlation R between observed and predicted fold change in translation initiation for individual mRNA features. Each feature was tested as the sole predictor of the fold change in translation initiation using the M5Rules model in Weka with 10-fold cross validation. eIF3c$^{KD}$ is shown in grey and the triple mutant is shown in white. TI wt - translation initiation in wild-type.

(**B**) Functional enrichment amongst genes rank-ordered by wt translation initiation. For each test, we used Gene Set Enrichment Analysis (www.webgestalt.org) using Gene Ontology and either the Biological Process or Cellular Component branch. Plots show categories selected from all significant categories (false discovery rate<0.05). Y-axis shows normalized enrichment score, i.e. the direction of the function bias (enrichment or depletion. eIF3c$^{KD}$ is shown in grey and the triple mutant in white.

(**C**) Enrichment in genes with alternative N-terminal isoforms. Overlap between 461 triple mutant genes that are significantly dysregulated in red (p-value < 0.05), and 2,275 genes with alternative N-terminal isoforms (blue) taken from[11]. The overlap shows 96 genes.

(**D**) Diagram of 5' UTR of PPP1CB with in-frame CUG, 27 nt upstream of the main translon' AUG with increased TI in the triple mutant. The graph shows the nucleotide (nt) position from the 5' end on the x-axis and the stem height gives the change in initiation, log2 (TI), at each alternate start codon (circles) and the change in translation efficiency, $\log_2$ (TE) at the main translon's AUG (diamond). Stop codons are denoted by a cross and color denotes reading frames, with frame 0 as the frame of the main translon's AUG.

(**E**) Diagram of 5' UTR of CITED2 with an in-frame AUA, 183 nt upstream of main translon's AUG. The graph shows the nucleotide (nt) position from the 5' end on the x-axis and the stem height gives the change in initiation, $\log_2$ (TI), at each alternate start codon (circles) and the change in translation efficiency, $\log_2$ (TE) at the main translon's AUG (diamond). Stop codons are denoted by a cross and color denotes reading frames, with frame 0 as the frame of the main translon's AUG.

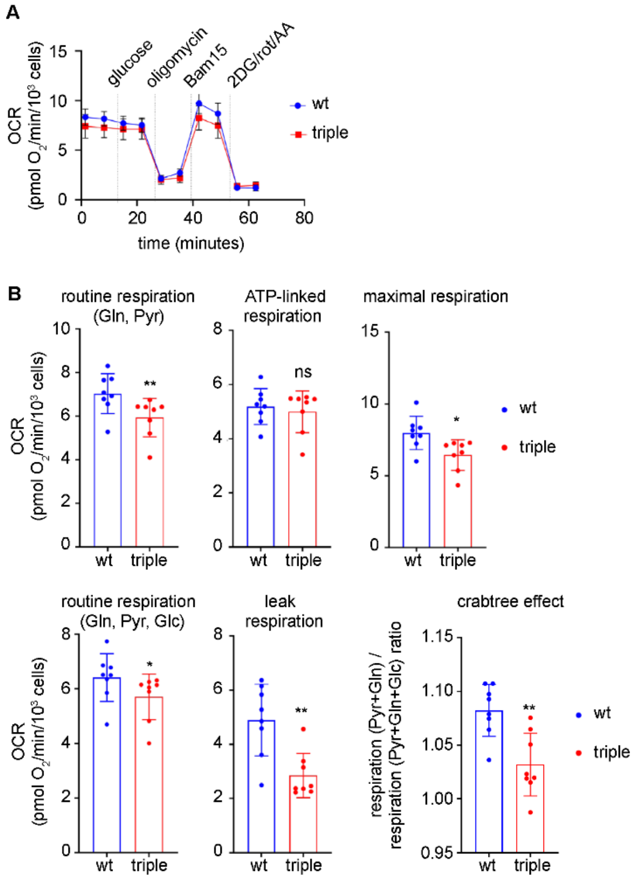


**Figure S9. Impaired eIF3c function compromises mitochondrial activity in adherent cells.**

(**A**) Seahorse XF analysis of oxygen consumption rate in wt and triple mutant cells. Mean OCR (oxygen consumption rate) traces ± S.D. in adherent wt and triple mutant cells. Curves show time-resolved changes in OCR under sequential substrate/inhibitor conditions; biological replicates: wt (n=8), triple mutant (n=8).

(**B**) Quantification of respiratory parameters derived from OCR measurements. Routine respiration was assessed under glutamine + pyruvate (Gln, Pyr), and glutamine + pyruvate + glucose (Gln, Pyr, Glc) conditions. ATP-linked respiration was calculated as the oligomycin-sensitive fraction. Leak respiration was determined after oligomycin addition, and maximal respiration was determined after Bam15 uncoupler addition. The Crabtree effect was calculated as the ratio of respiration before (Glc,Pyr) and after the addition of glucose (Glc, Pyr, Glc). Data are expressed as OCR per $10^3$ cells. Values represent mean ± S.D.; biological replicates: wt (n=8), triple mutant (n=8). Statistical significance was tested by unpaired two-tailed *t*-test or nonparametric Mann Whitney U test (* $p \le 0.05$; ** $p \le 0.01$; ns, not significant).

#### SUPPLEMENTARY TABLES

**Supplementary Table S1.** Please see Materials and Methods, chapter “CRISPR/Cas9 stable cell line generation”, where it was placed for the sake of clarity.

**Supplementary Table S2.** Recombinant DNA plasmids.

| plasmid | description | source |
|---|---|---|
| pGL4-SV40-firefly luc2 | Reporter plasmid, Firefly Luciferase encoded by ATG start codon | gift from Petr Svoboda’s lab |
| pGL4-SV40-CTG-firefly luc2 | Reporter plasmid, Firefly Luciferase with start codon mutated to CTG | [18] |
| pGL4-SV40-GTG-firefly luc2 | Reporter plasmid, Firefly Luciferase with start codon mutated to GTG | [18] |
| pGL4-SV40-TTG-firefly luc 2 | Reporter plasmid, Firefly Luciferase with start codon mutated to TTG | [18] |
| pGL4-SV40-NAT1-GTG | Reporter plasmid, Firefly Luciferase with NAT1/DAP5/eIF4G2 5UTR sequence initiating from GTG | This study |
| pGL4-SV40-NAT1-ATG | Reporter plasmid, Firefly Luciferase with NAT1/DAP5/eIF4G2 5UTR sequence initiating from ATG | This study |
| pGL4-SV40-AAG- firefly luc2 | Reporter plasmid, Firefly Luciferase with start codon mutated to AAG | This study |
| pGL4-SV40-5UTR_eIF1_native_context | Reporter plasmid, Firefly Luciferase with eIF1 native 5UTR sequence | [1] |
| pGL4-SV40-5UTR_eIF1_strong_context | Reporter plasmid, Firefly Luciferase with eIF1 strong 5UTR sequence | [1] |
| pGL4-SV40-5UTR_eIF5_native | Reporter plasmid, Firefly Luciferase with eIF5 native 5UTR sequence | [1] |

| | | |
|---|---|---|
| pGL4-SV40-5UTR_eIF5_uORF-mutated | Reporter plasmid, Firefly Luciferase with eIF5 ORF-less 5UTR sequence | [1] |
| pGL4-CMV-h3c-1-325 | *H.sapiens* eIF3c 1-325 under T7 promotor; for *in vitro* expression | [20] |
| pGL4-CMV-h3c | *H.sapiens* eIF3c under T7 promotor; for *in vitro* expression | [20] |
| pGL4-CMV-h3c-7-325 | *H.sapiens* eIF3c 7-325 under T7 promotor; for *in vitro* expression | This study |
| pGL4-CMV-h3c-1-325 R3N | *H.sapiens* eIF3c 1-325 with R3N substitution under T7 promotor; for *in vitro* expression | This study |
| pGL4-CMV-h3c-1-325 F4A | *H.sapiens* eIF3c 1-325 with F4A substitution under T7 promotor; for *in vitro* expression | This study |
| pGL4-CMV-h3c-1-325 F5A | *H.sapiens* eIF3c 1-325 with F5A substitution under T7 promotor; for *in vitro* expression | This study |
| pGL4-CMV-h3c-1-325 F4A F5A | *H.sapiens* eIF3c 1-325 with F4AF5A substitution under T7 promotor; for *in vitro* expression | This study |
| pGL4-CMV-h3c-1-325 R3N F4A F5A | *H.sapiens* eIF3c 1-325 with R3NF4AF5A substitution under T7 promotor; for *in vitro* expression | This study |
| pGL4-CMV-h3c R3N F4A F5A | *H.sapiens* eIF3c (whole protein) with R3NF4AF5A substitution under T7 promotor; for *in vitro* expression | This study |
| pGL4-CMV-eIF3c-1-180-WT-c-FLAG | *H.sapiens* eIF3c 1-180; for *in vivo* expression, FLAG-tagged | This study |
| pGL4-CMV-eIF3c-1-180-R3N-c-FLAG | *H.sapiens* eIF3c 1-180; for *in vivo* expression, FLAG-tagged | This study |
| pGL4-CMV-eIF3c-1-180-F4A-c-FLAG | *H.sapiens* eIF3c 1-180; for *in vivo* expression, FLAG-tagged | This study |
| pGL4-CMV-eIF3c-1-180-F5A-c-FLAG | *H.sapiens* eIF3c 1-180; for *in vivo* expression, FLAG-tagged | This study |
| pGL4-CMV-eIF3c-1-180-R3N-F4A-F5A-c-FLAG | *H.sapiens* eIF3c 1-180; for *in vivo* expression, FLAG-tagged | This study |

| | | |
|---|---|---|
| pGL4-CMV-eIF3c-1-325-WT-c-FLAG | *H.sapiens* eIF3c 1-325; for *in vivo* expression, FLAG-tagged | This study |
| pGL4-CMV-eIF3c-1-325-R3N-F4A-F5A-c-FLAG | *H.sapiens* eIF3c 1-325; for *in vivo* expression, FLAG-tagged | This study |
| pGL4-CMV-eIF3c-1-913-R3N-F4A-F5A-c-FLAG | *H.sapiens* eIF3c 1-913 (FL); for *in vivo* expression, FLAG-tagged | This study |
| pGL4-CMV-eIF3c-1-913-WT-c-FLAG | *H.sapiens* eIF3c 1-913 (FL); for *in vivo* expression, FLAG-tagged | This study |
| pGL4-CMV-h eIF1 | *H.sapiens* eIF1, for *in vivo* production | This study |
| pGL4-CMV h-eIF5 | *H.sapiens* eIF5, for *in vivo* production | This study |
| pGEX-heIF5 | *H.sapiens* GST-eIF5 under Tac promotor and Lac operator control; for production in *E.Coli* | [20] |
| pX330-U6-Chimeric_BB-CBh-hSpCas9 | vector for Crispr/Cas9 system | Addgene #42230, gift from David Staněk's lab |
| pX330-h3c- gRNA_1 | encodes spCas9 and gRNA1 | This study |
| pX330-h3c-gRNA_2 | encodes spCas9 and gRNA2 | This study |
| pX330-h3c-gRNA_3 | encodes spCas9 and gRNA3 | This study |
| pX330-h3c-gRNA_5 | encodes spCas9 and gRNA5 | This study |
| pX330-h3c-gRNA_6 | encodes spCas9 and gRNA6 | This study |
| pARV-GFP-RFP | Reporter plasmid for Crispr/Cas9 system | gift from David Staněk's lab |
| pARV-GFP-RFP-gRNA_1-ON_target | Reporter plasmid, GFP, RFP interrupted by gRNA1 target sequence | This study |
| pARV-GFP-RFP-gRNA_1-off_target_1 | Reporter plasmid, GFP, RFP interrupted by gRNA1 off-target 1 sequence | This study |
| pARV-GFP-RFP-gRNA_1-off_target_2 | Reporter plasmid, GFP, RFP interrupted by gRNA1 off-target 2 sequence | This study |
| pARV-GFP-RFP-gRNA_1-off_target_3 | Reporter plasmid, GFP, RFP interrupted by gRNA1 off-target 3 sequence | This study |

| | | |
|---|---|---|
| pARV-GFP-RFP-gRNA_2-ON_target | Reporter plasmid, GFP, RFP interrupted by gRNA2 target sequence | This study |
| pARV-GFP-RFP-gRNA_2-off_target_1 | Reporter plasmid, GFP, RFP interrupted by gRNA2 off-target 1 sequence | This study |
| pARV-GFP-RFP-gRNA_2-off_target_2 | Reporter plasmid, GFP, RFP interrupted by gRNA2 off-target 2 sequence | This study |
| pARV-GFP-RFP-gRNA_2-off_target_3 | Reporter plasmid, GFP, RFP interrupted by gRNA2 off-target 3 sequence | This study |
| pARV-GFP-RFP-gRNA_2-off_target_4 | Reporter plasmid, GFP, RFP interrupted by gRNA2 off-target 4 sequence | This study |
| pARV-GFP-RFP-gRNA_2-off_target_5 | Reporter plasmid, GFP, RFP interrupted by gRNA2 off-target 5 sequence | This study |
| pARV-GFP-RFP-gRNA_3-ON_target | Reporter plasmid, GFP, RFP interrupted by gRNA3 target sequence | This study |
| pARV-GFP-RFP-gRNA_3-off_target_1 | Reporter plasmid, GFP, RFP interrupted by gRNA3 off-target 1 sequence | This study |
| pARV-GFP-RFP-gRNA_3-off_target_2 | Reporter plasmid, GFP, RFP interrupted by gRNA3 off-target 2 sequence | This study |
| pARV-GFP-RFP-gRNA_3-off_target_3 | Reporter plasmid, GFP, RFP interrupted by gRNA3 off-target 3 sequence | This study |
| pARV-GFP-RFP-gRNA_5-ON_target | Reporter plasmid, GFP, RFP interrupted by gRNA5 target sequence | This study |
| pARV-GFP-RFP-gRNA_5-off_target_1 | Reporter plasmid, GFP, RFP interrupted by gRNA5 off-target 1 sequence | This study |
| pARV-GFP-RFP-gRNA_5-off_target_2 | Reporter plasmid, GFP, RFP interrupted by gRNA5 off-target 2 sequence | This study |

| | | |
|---|---|---|
| pARV-GFP-RFP-gRNA_5-off_target_3 | Reporter plasmid, GFP, RFP interrupted by gRNA5 off-target 3 sequence | This study |
| pARV-GFP-RFP-gRNA_6-ON_target | Reporter plasmid, GFP, RFP interrupted by gRNA6 target sequence | This study |
| pARV-GFP-RFP-gRNA_6-off_target_1 | Reporter plasmid, GFP, RFP interrupted by gRNA6 off-target 1 sequence | This study |
| pARV-GFP-RFP-gRNA_6-off_target_2 | Reporter plasmid, GFP, RFP interrupted by gRNA6 off-target 2 sequence | This study |
| pARV-GFP-RFP-gRNA_6-off_target_3 | Reporter plasmid, GFP, RFP interrupted by gRNA6 off-target 3 sequence | This study |
| pGL4-CMV-h3c-exon1-gRNA1-WT | genomic sequence surrounding gRNA1 target site in eIF3c cloned into pGL4-CMV vector; control for genotyping | This study |
| pGL4-CMV-h3c-exon1-gRNA1-R3N | genomic sequence surrounding gRNA1 target site in eIF3c with R3N substitution cloned into pGL4-CMV vector; control for genotyping | This study |
| pGL4-CMV-h3c-exon1-gRNA1-F5A | genomic sequence surrounding gRNA1 target site in eIF3c with F5A substitution cloned into pGL4-CMV vector; control for genotyping | This study |
| pGL4-CMV-h3c-exon1-gRNA1-R3N-F4A-F5A | genomic sequence surrounding gRNA1 target site in eIF3c with R3N-F4A-F5A substitution cloned into pGL4-CMV vector; control for genotyping | This study |

**Supplementary Table S3.** List of oligonucleotides and genestrings

| | |
|---|---|
| pGL4_AAG_LUC | GGCCTAGGCTTTTGCAAAAAGCTTGGC<br>AATCCGGTACTGTTGGTAAAGCCACCA<br>AGGAAGATGCCAAAAACATTAAGAAGG<br>GCCCAGCGCCATTCTACCCACTCGAAG<br>ACGGGACCGCCGGCGAGCAGCTGCAC<br>AAAGCCATGAAGCGCTACGCCCTGGTG<br>CCCGGCACCATCGCCTTTACCGACGCA<br>CATATCGAGGTGGACATTACCTACGCC<br>GAGTACTTCGAGATGAGCGTTCGGCTG<br>GCAGAAGCTATGAAGCGCTATGGGCTG<br>AATACAAACCATCGGATCGTGGTGTGC<br>AGCGAGAATAGCTTGCAGTTCTTCATG<br>CCCGTGTTGGGTGCCCTGTTCATCGGT<br>GTGGCTGTGGCCCCAGCTAACGACATC<br>TACAACGAGCGCGAGCTGCTGAACAGC<br>ATGGGCATCAGCCAGCCCACCGTCGTA<br>TTCGTGAGCAAGAAAGGGCTGCAAAAG<br>ATCCTCAACGTGCAAAAGAAGCTACCG<br>ATCATACAAAAGATCATCATCATGGATA<br>GCAAGACCGACTACCAGGGCTTCCAAA<br>GCATGTACACCTTCGTGACTTCCCATTT<br>GCCACCCGGCTTCAACGAGTACGACTT<br>CGTG |
| NAT1_AUG_GTG | CAAGATAAGCTTGATTCTTCGTTGTCAA<br>GCCGCCAAAGTGGAGGATGCCAAAAAC<br>ATTAAGAAGGGCCCAGCGCCATTCTAC<br>CCACTCGAAGACGGGACCGCCGGCGA<br>GCAGCTGCACAAAGCCATGAAGCGCTA<br>CGCCCTGGTGCCCGGCACCATCGCCT<br>TTACCGACGCACATATCGAGGTGGACA<br>TTACCTACGCCGAGTACTTCGAGATGA<br>GCGTTCGGCTGGCAGAAGCTATGAAGC<br>GCTATGGGCTGAATACAAACCATCGGA<br>TCGTGGTGTGCAGCGAGAATAGCTTGC<br>AGTTCTTCATGCCCGTGTTGGGTGCCC<br>TGTTCATCGGTGTGGCTGTGGCCCCAG<br>CTAACGACATCTACAACGAGCGCGAGC<br>TGCTGAACAGCATGGGCATCAGCCAGC<br>CCACCGTCGTATTCGTGAGCAAGAAAG |

| | |
|---|---|
| | GGCTGCAAAAGATCCTCAACGTGCAAA AGAAGCTACCGATCATACAAAAGATCAT CATCATGGATAGCAAGACCGACTACCA GGGCTTCCAAAGCATGTACACCTTCGT GACTTCCCATTTGCCACCCG |
| NAT1_AUG_ATG | CAAGATAAGCTTGATTCTTCGTTGTCAA GCCGCCAAAATGGAGGATGCCAAAAAC ATTAAGAAGGGCCCAGCGCCATTCTAC CCACTCGAAGACGGGACCGCCGGCGA GCAGCTGCACAAAGCCATGAAGCGCTA CGCCCTGGTGCCCGGCACCATCGCCT TTACCGACGCACATATCGAGGTGGACA TTACCTACGCCGAGTACTTCGAGATGA GCGTTCGGCTGGCAGAAGCTATGAAGC GCTATGGGCTGAATACAAACCATCGGA TCGTGGTGTGCAGCGAGAATAGCTTGC AGTTCTTCATGCCCGTGTTGGGTGCCC TGTTCATCGGTGTGGCTGTGGCCCCAG CTAACGACATCTACAACGAGCGCGAGC TGCTGAACAGCATGGGCATCAGCCAGC CCACCGTCGTATTCGTGAGCAAGAAAG GGCTGCAAAAGATCCTCAACGTGCAAA AGAAGCTACCGATCATACAAAAGATCAT CATCATGGATAGCAAGACCGACTACCA GGGCTTCCAAAGCATGTACACCTTCGT GACTTCCCATTTGCCACCCG |
| TP-h3c-7-325-PmeI | ATTAAGTTTAAACGCCATGACCGGTTC GGACAGCGAGTCCGAGTCG |
| AH-h3c-325-FseI | ATATACGGCCGGCCTCAGGTGATCTCA GTTCCCTTGGC |
| TP-h3c-1-325-R3N-PmeI-FW | CTAGCGTTTAAACGCCATGTCGAACTTT TTCACCACCGGTTCGGACAGC |
| KP_pGL4-CMV_h3c_1-325_F4A_F | TTATAGTTTAAACGCCATGTCGCGGGC TTTCACCACCGGTTCGGACAGCGAG |
| KP_pGL4-CMV_h3c_1-325_F5A_F | CTAATGTTTAAACGCCATGTCGCGGTTT GCCACCACCGGTTCGGACAGCGAGTC |

| | |
|---|---|
| TP-h3c-1-325-F4A-F5A | ATATAGTTTAAACGCCATGTCGCGGGC TGCCACCACCGGTTCGGACAGCGAG |
| TP-h3c-1-325-R3N-F4A-F5A | TTATAGTTTAAACGCCATGTCGAACGCT GCCACCACCGGTTCGGACAGCGAGTC C |
| AH-h3c-FseI-R | ATATAGGCCGGCCTCAGTAGGCCGTCT GAGACTG |
| AH-h3c-SphI | ACAACATTCAGCATGCAGACC |
| AH-h3c-FLAG2-R | GAAGCGGCCGGCCTCACTTGTCATCGT CGTCCTTGTAGTCGTAGGCCGTCTGAG ACTGCTGCTGGCGGT |
| eIF3c_HindIII_kATG_WT_F | CAAGATAAGCTTGCCACCATGTCGCGG TTTTTC |
| eIF3c_HindIII_kATG_R3N_F | CAAGATAAGCTTGCCACCATGTCGAAC TTTTTCACCAC |
| eIF3c_HindIII_kATG_F4A_F | CAAGATAAGCTTGCCACCATGTCGCGG GCTTTCACCAC |
| eIF3c_HindIII_kATG_F5A_F | CAAGATAAGCTTGCCACCATGTCGCGG TTTGCTACCACC |
| eIF3c_HindIII_R3NF4AF5A_F | CAAGATAAGCTTGCCACCATGTCGAAC GCTGCTACCACCG |
| eIF3c_FseI_FLAG_180a_R | GTCGTCATCGTCTTTGTAGTCGCCTTCT GAATCCTC |
| eIF3c_FseI_FLAG_180a_R | GTCGTCATCGTCTTTGTAGTCGCCTTCT GAATCCTC |
| eIF3c_325_NAA_T95_FWD | AGACGGATCCATGTCGAACGCTGCCAC |
| eIF3c_FseI_FLAG_325_R | GTCGTCATCGTCTTTGTAGTCGGTGAT CTCAGTTCC |
| FseI_FLAG_STOP_shrt_R | TATATGGCCGGCCCTACTTGTCGTCAT CGTC |
| pX330-h3c-qRNA_1_sense | CACCGTCCGCGCCGTCGCCATGTCG |
| pX330-h3c-gRNA_1_antisense | AAACCGACATGGCGACGGCGCGGAC |
| pX330-h3c-gRNA_2_sense | CACCGGTGGTGAAAAACCGCGACA |
| pX330-h3c-gRNA_2_antisense | AAACTGTCGCGGTTTTTCACCACC |
| pX330-h3c-gRNA_3_sense | CACCGGACTCGCTGTCCGAACCGG |
| pX330-h3c-gRNA_3_antisense | AAACCCGGTTCGGACAGCGAGTCC |
| pX330-gRNA_5-sense | CACCGATGTCGCGGTTTTTCACCAC |
| pX330-h3c-gRNA_5-antisense | AAACGTGGTGAAAAACCGCGACATC |
| pX330-h3c-gRNA_6-sense | CACCGCGGTTTTTCACCACCGGTT |
| pX330-h3c-gRNA_6-antisense | AAACAACCGGTGGTGAAAAACCGC |

| | |
|---|---|
| pARV-h3c-gRNA_1-on_target-sense | GGATCCACCTCCGCGCCGTCGCCATGTCGCGGTTTCGAT |
| pARV-h3c-gRNA_1-on_target-antisense | CGAAACCGCGACATGGCGACGGCGCGGAGGTGGATCC |
| pARV-h3c-gRNA_1-off_target1-sense | GGATCCCTCCCACGACCTGGAGACGGCGCTGAGGCCGAT |
| pARV-h3c-gRNA_1-off_target1-antisense | CGGCCTCAGCGCCGTCTCCAGGTCGTGGGAGGGATCC |
| pARV-h3c-gRNA_1-off_target_2-sense | GGATCCCCGTCCGCGCCGCCGCCATCCTGGGGTCCCGAT |
| pARV-h3c-gRNA_1-off_taget_2-antisense | CGGGACCCCAGGATGGCGGCGGCGCGGACGGGGATCC |
| pARV-h3c-gRNA_1-off_target_3-sense | GGATCCGAGCCCCGACGCGGCGGCGGCGCGGGGCTCGAT |
| pARV-h3c-gRNA_1-off_target_3-antisense | CGAGCCCCGCGCCGCCGCCGCGTCGGGGCTCGGATCC |
| pARV-h3c-gRNA_2-on_target-sense | GGATCCTCGCCATGTCGCGGTTTTTCACCACCGGTCGAT |
| pARV-h3c-gRNA_2-on-target-antisense | CGACCGGTGGTGAAAAACCGCGACATGGCGAGGATCC |
| pARV-h3c-gRNA_2-off_target_1-sense | GGATCCCAAGGTGGTGAAAAACCTCTACAAGGAAACGAT |
| pARV-h3c-gRNA_2-off_target_1-antisense | CGTTTCCTTGTAGAGGTTTTTCACCACCTTGGGATCC |
| pARV-h3c-gRNA_2-off_target_2-sense | GGATCCTTTCCTTGTAGCGGTCTTTCACCTCCTTGCGAT |
| pARV-h3c-gRNA_2-off_target_2-antisense | CGCAAGGAGGTGAAAGACCGCTACAAGGAAAGGATCC |
| pARV-h3c-gRNA_2-off_target_3-sense | GGATCCCAAGGAGGTGAAAGACCTCGACAAGGAGACGAT |
| pARV-h3c-gRNA_2-off_target_3-antisense | CGTCTCCTTGTCGAGGTCTTTCACCTCCTTGGGATCC |
| pARV-h3c-gRNA_2-off_target_4-sense | GGATCCTTTCCTTGTAGAGGTTTTTCACCGCCTTGCGAT |
| pARV-h3c-gRNA_2-off_target_4-antisense | CGCAAGGCGGTGAAAAACCTCTACAAGGAAAGGATCC |
| pARV-h3c-gRNA_2-off_target_5-sense | GGATCCTGAGAAGGTGAACAGCCGCGACACGGCGGCGAT |
| pARV-h3c-gRNA_2-off_target-5-antisense | CGCCGCCGTGTCGCGGCTGTTCACCTTCTCAGGATCC |

| | |
|---|---|
| pARV-h3c-gRNA_3-on_target-sense | GGATCCTCACCACCGGTTCGGACAGCGAGTCCGAGCGAT |
| pARV-h3c-gRNA_3-on_target-antisense | CGCTCGGACTCGCTGTCCGAACCGGTGGTGAGGATCC |
| pARV-h3c-gRNA_3-off_target_1-sense | GGATCCACGGGACTCACTGTCCGAACAGGAGGAGACGAT |
| pARV-h3c-gRNA_3-off_target_1-antisense | CGTCTCCTCCTGTTCGGACAGTGAGTCCCGTGGATCC |
| pARV-h3c-gRNA_3-off_target_2-sense | GGATCCGCGCCCCCGGTTCGGGCAGCGGCTGCGGCCGAT |
| pARV-h3c-gRNA_3-off_target_2-antisense | CGGCCGCAGCCGCTGCCCGAACCGGGGGCGCGGATCC |
| pARV-h3c-gRNA_3-off_target_3-sense | GGATCCGGGGGAGCCGCTGTCCGAGCGGGAGGTGACGAT |
| pARV-h3c-gRNA_3-off_target_3-antisense | CGTCACCTCCCGCTCGGACAGCGGCTCCCCCGGATCC |
| pARV-h3c-gRNA_5-on_target-sense | GGATCCGCCATGTCGCGGTTTTTCACCACCGGTTCCGAT |
| pARV-h3c-gRNA_5-on_target-antisense | CGGAACCGGTGGTGAAAAACCGCGACATGGCGGATCC |
| pARV-h3c-gRNA_5-off_target_1-sense | GGATCCATTTCTATGGTGAAAAACAGTGACATCTTCGAT |
| pARV-h3c-gRNA_5-off_target_1-antisense | CGAAGATGTCACTGTTTTTCACCATAGAAATGGATCC |
| pARV-h3c-gRNA_5-off_target_2-sense | GGATCCTGGATATGCGGTTTTTCACCACAGGCCTCGAT |
| pARV-h3c-gRNA_5-off_target_2-antisense | CGAGGCCTGTGGTGAAAAACCGCATATCCAGGATCC |
| pARV-h3c-gRNA_5-off_target_3-sense | GGATCCTCTATGTCGTGGGTTCCCACCACAGGCTGCGAT |
| pARV-h3c-gRNA_5-off_target_3-antisense | CGCAGCCTGTGGTGGGAACCCACGACATAGAGGATCC |
| pARV-h3c-gRNA_6-on_target_sense | GGATCCGTCGCGGTTTTTCACCACCGGTTCGGACACGAT |
| pARV-h3c-gRNA_6-on_target-antisense | CGTGTCCGAACCGGTGGTGAAAAACCGCGACGGATCC |
| pARV-h3c-gRNA_6-off_target_1-sense | GGATCCGTGCCCAACTGGTGGTGAACAACCGCTGGCGAT |
| pARV-h3c-gRNA_6-off_target_1-antisense | CGCCAGCGGTTGTTCACCACCAGTTGGGCACGGATCC |

| | |
|---|---|
| pARV-h3c-gRNA_6-off_target_2-sense | GGATCCGATGCGGTTTTGCACCTCTGGTTAGAGGCCGAT |
| pARV-h3c-gRNA_6-off_target_2-antisense | CGGCCTCTAACCAGAGGTGCAAAACCGCATCGGATCC |
| pARV-h3v-gRNA_6-off_target_3-sense | GGATCCTCGCCAAACTGGTGAAAAACCGGACCCGAT |
| pARV-h3c-gRNA_6-off_target_3-antisense | CGGGTCCGGTTTTTCACCAGTTTGGCGAGGATCC |
| TP-h3c-5UTR-EcoRI-FW | TACAGGAATTCGCGTGAGTGAGCCACCGCAC |
| TP-h3c-intron-FseI-R | TATATGGCCGGCCTGAGATCTGAAGGTCGAGGCAGC |
| TP-h3c-fushion-pair2_FW | ACCACCGGTTCGGACAGCGAGTC |
| TP-gen_h3c-R3N-gRNA1-fushion_pair1_R | CTCGCTGTCCGAACCGGTGGTGAAAAAGTTCGACATGGCGACGGCGCGGAG |
| TP-h3c-F5A_gRNA1-fushion_pair1-R | CTCGCTGTCCGAACCGGTGGTCGCGAAGCGCGACATGGCGACGGCGCGG |
| eIF1-native_LUC | GGAGGCTTTTTTGGAGGCCTAGGCTTTTGCAAAAAGCTTGCCCCAGTCACTGAGCCGCCGCCGAGGATTCAGCAGCCTCCCCCTTGAGCCCCCTCGCTTCCCGACGTTCCGTTCCCCCCTGCCCGCCTTCTCCCGCCACCGCCGCCGCCGCCTTCCGCAGGCCGTTTCCACCGAGGAAAAGGAATCGTATCGTATGTCCGAAGATGCCAAAAACATTAAGAAGGGCCCAGCGCCATTCTACCCACTCGAAGACGGGACCGC |
| eIF1-strong_LUC | GGAGGCTTTTTTGGAGGCCTAGGCTTTTGCAAAAAGCTTGCCCCAGTCACTGAGCCGCCGCCGAGGATTCAGCAGCCTCCCCCTTGAGCCCCCTCGCTTCCCGACGTTCCGTTCCCCCCTGCCCGCCTTCTCCCGCCACCGCCGCCGCCGCCTTCCGCAGGCCGTTTCCACCGAGGAAAAGGAATCGGCCACCATGGCCGAAGATGCCAAAAACATTAAGAAGGGCCCAGCGCCATTCTACCCACTCGAAGACGGGACCGC |
| eIF5-native_LUC | TGGCCTAACTGGCCGGTACCTGAGCTCGCTAGCCTCGAGGATATCAAGATCTGCGCAGCACCATGGCCTGAAATAACCTCT |

| | |
|---|---|
| | GAAAGAGGAACTTGGTTAGGTACCTTC TGAGGCGGAAAGAACCAGCTGTGGAAT GTGTGTCAGTTAGGGTGTGGAAAGTCC CCAGGCTCCCCAGCAGGCAGAAGTAT GCAAAGCATGCATCTCAATTAGTCAGC AACCAGGTGTGGAAAGTCCCCAGGCTC CCCAGCAGGCAGAAGTATGCAAAGCAT GCATCTCAATTAGTCAGCAACCATAGTC CCGCCCCTAACTCCGCCCATCCCGCCC CTAACTCCGCCCAGTTCCGCCCATTCT CCGCCCCATGGCTGACTAATTTTTTTTA TTTATGCAGAGGCCGAGGCCGCCTCG GCCTCTGAGCTATTCCAGAAGTAGTGA GGAGGCTTTTTTGGAGGCCTAGGCTTT TGCAAAAAGCTTAGTCAGAGCGAGAAC ATTCCAGAGGTCGCCCAGCTCCGGCG CTGACGGGTGTGGACCGCGGACGTCG CTGGGACAGCCCCTCCCCGCTGCTCG GCGGCGGCACCTGGCCCGGCCGCTCC TCGCTGCGCTTCGCCTCCGCCTCCTCG GACTCGGACTCGGGTTTATATCGCGCC TCACTTCATCCCAGTCCCGGGCGAGCA GCGTTGGGTTTATGTCTTTATTTGACGA AAACGAGCTGTTGCGCAGCCATTGGTA CCTGTATTGGGGAAACATAGCATACAA GCAAGAAGCTTACAGCCTCAGTGGCGA AAATTTTTTCATGTCAGAGACCGAGAAC TCTTGCAGTCGTTTATGTCATCCCTTCT TCTCCAGACAGAAGATACCAAAAAGTT GCAATCAAAGATCTCTTCATCTTATTGA TAAAGCCACTAATAAGCCAAAATGTCTG TCAATGTCAACCGCAGCGTGTCAGACC AGTTCTATCGCTACAAGATGCCCCGTC TGATTGAAGATGCCAAAAACATTAAGAA GGGCCCAGCGCCATTCTACCCACTCGA AGACGGGACCGC |
| eIF5-uORF-less_LUC | TGGCCTAACTGGCCGGTACCTGAGCTC GCTAGCCTCGAGGATATCAAGATCTGC GCAGCACCATGGCCTGAAATAACCTCT GAAAGAGGAACTTGGTTAGGTACCTTC |

| | |
|---|---|
| | TGAGGCGGAAAGAACCAGCTGTGGAAT GTGTGTCAGTTAGGGTGTGGAAAGTCC CCAGGCTCCCCAGCAGGCAGAAGTAT GCAAAGCATGCATCTCAATTAGTCAGC AACCAGGTGTGGAAAGTCCCCAGGCTC CCCAGCAGGCAGAAGTATGCAAAGCAT GCATCTCAATTAGTCAGCAACCATAGTC CCGCCCCTAACTCCGCCCATCCCGCCC CTAACTCCGCCCAGTTCCGCCCATTCT CCGCCCCATGGCTGACTAATTTTTTTTA TTTATGCAGAGGCCGAGGCCGCCTCG GCCTCTGAGCTATTCCAGAAGTAGTGA GGAGGCTTTTTTGGAGGCCTAGGCTTT TGCAAAAAGCTTAGTCAGAGCGAGAAC ATTCCAGAGGTCGCCCAGCTCCGGCG CTGACGGGTGTGGACCGCGGACGTCG CTGGGACAGCCCCTCCCCGCTGCTCG GCGGCGGCACCTGGCCCGGCCGCTCC TCGCTGCGCTTCGCCTCCGCCTCCTCG GACTCGGACTCGGGTTTATATCGCGCC TCACTTCATCCCAGTCCCGGGCGAGCA GCGTTGGGTTTAAATCTTTATTTGACGA AAACGAGCTGTTGCGCAGCCATTGGTA CCTGTATTGGGGAAACATAGCATACAA GCAAGAAGCTTACAGCCTCAGTGGCGA AAATTTTTTCAAATCAGAGACCGAGAAC TCTTGCAGTCGTTTAAATCATCCCTTCT TCTCCAGACAGAAGATACCAAAAAGTT GCAATCAAAGATCTCTTCATCTTATTGA TAAAGCCACTAATAAGCCAAAATGTCTG TCAATGTCAACCGCAGCGTGTCAGACC AGTTCTATCGCTACAAGATGCCCCGTC TGATTGAAGATGCCAAAAACATTAAGAA GGGCCCAGCGCCATTCTACCCACTCGA AGACGGGACCGC |

| | |
|---|---|
| **Primers used for genotyping** | |
| TP_eIF3c_FW genotyp pair1 | CACCGCACCCGGCAGGATTATCTAC |
| TP_eIF3c_R genotyp pair1 | TGGGTGGCTACCGGCTATCACAG |
| TP_eIF3c_FW genotyp pair2 | TCCCAGTGTAGTTAGTCGGCGCTTC |
| TP_eIF3c_R genotyp pair2 | CACAGCAGGTGACCAGACTTCAGAG |
| TP_eIF3c_FW genotyp pair 3 | AGACCTGCAAGATCTAGC |
| TP_eIF3c_R genotyp pair 3 | CGTTGGTGTCGGGATTTA |
| TP_eIF3c_R_genotyping pair 4 | GCCTCAGCCTCCTAAGTTG |
| **Crispr/Cas9 donor DNA** | |
| ssODN gRNA1 R3N | AGGACGACTCGGACTCGCTGTCCGAACCGGTGGTGAAAAAGTTCGACATGGCGACGGCGCGGAGGTGCTACGGCCGGACCTGCGGG |
| ssODN gRNA1 F5A | CGGACAAGGACGACTCGGACTCGCTGTCCGAACCGGTGGTCGCGAAGCGCGACATGGCGACGGCGCGGAGGTGCTACGGCCGGACCTGCGGG |
| ssODN gRNA1 R3N-F4A-F5A | CGGACAAGGACGACTCGGACTCGCTGTCCGAACCGGTGGTGGCAGCGTTCGACATGGCGACGGCGCGGAGGTGCTACGGCCGGACCTGCGGG |
| **qPCR primers** | |
| SW-18S-human | CCGCAGCTAGGAATAATG |
| SW-18S-human-R | CCGGTCCAAGAATTTCAC |
| RPL41a_fwd (SPIKE) | CGAAATGAGAGCCAAGTGG |
| RPL41a_rev (SPIKE) | ATGCAATTTAGATCCATTATGAGG |
| AH-huALAS_Fwd | CCACTGGAAGAGCTGTGTGATGTG |
| AH-huALAS_Rev | GCGATGTACCCTCCAACACAACC |
| SW pGL4-FF | ACGAGGTGCCTAAAGGACTG |
| SW pGL4-FF R | CGCCCTTCTTGGCCTTAATG |
| SW h3c 267 | GAGTCAGTGCTGCAACTTTC |
| SW h3c 267 R | TAGCTTTGTCCTCCCGTTTC |
| SW h3k 253 | GTTCAACCCAGCCTTCTTTC |

| | |
|---|---|
| SW h3k 253-R | TCGGACAGAGTCTTCAAAGC |
| SW h3l 269 | ATTGCCCTCACGATGTACCC |
| SW h3l 269-R | CTGCGGATGGTTGAAAGCTG |
| SW h3d 267 | CCAACCCAAACCCGTTTGTG |
| SW h3d 267-R | TCTTCAGCTCCGTGGCAATG |
| SW h3e 253 | CTGGTTCCAGCAACAGATAG |
| SW h3e 253-R | GTGGCTGATAAAGGAAGAGG |
| TP_eIF5_qPCR_FW | CTGATTGCCAAGGTTGAGG |
| TP_eIF5_qPCR_R | ATCAAACTGGGTCTGTGCT |
| **siRNA** | |
| ON-TARGETplus Human EIF3C (8663) siRNA - SMARTpool (Dermacon # L-009036-00) | |
| ON-TARGETplus Human EIF3K (27335) siRNA - SMARTpool (Dermacon #L-020216-02) | |
| ON-TARGETplus Human EIF3L (51386) siRNA - SMARTpool (Dermacon #L-020949-01) | |
| ON-TARGETplus Human EIF3E (3646) siRNA - SMARTpool (Dermacon #L-010518-00) | |
| ON-TARGETplus Non-targeting siRNA #3 (Dermacon #D-001810-03) | |

**Supplementary Table S4.** Antibodies used in the study.

| antibody | source |
|---|---|
| eIF3a for Co-IP | Cell Signalling # 3411 |
| eIF5 for Co-IP | Santa cruz #sc-28309 |
| eIF3a | Sigma #HPA038315 |
| eIF3b | Thermo Scientific # PA5-23278 |
| eIF3c | Santa Cruz # sc-28858 |
| eIF3d | Sigma # HPA066216 |
| eIF3e | Thermo Scientific # PA5-29487 |
| eIF3f | Sigma # HPA049250 |
| eIF3g | Sigma # HPA041997 |
| eIF3h | Cell Signalling # 3413S |
| eIF3i | Sigma # HPA029939 |
| eIF3j | Santa Cruz # sc-50356 |
| eIF3k | Abcam # 85968 |
| eIF3l | Bethyl # A304-753A-M |
| eIF3m | Sigma # HPA031063 |
| eIF1A | Abcam # ab177939 |
| eIF2α | Cell Signalling # 5324 |
| eIF5 | Cell Signalling # 13894 |
| eIF1 | Cell Signalling # 12496S |
| FLAG M2 | Sigma # F1804 |
| GAPDH | Thermo Scientific # PA1987 |
| Rps14 | Abcam # ab174661 |
| RPS3 | Baria # 15198-1-AP |

**Supplementary Table S5. 3' Biotinylated (BITEG) rRNA depletion oligos used in the study.**

| Name | 5ˈto 3ˈ sequence | Percentage in the pool |
|---|---|---|
| AH-h18S-2 | GGTTTTGATCTGATAAATGCACGCATCCCCCCCG | 26 |
| AH-h18S-3 | TGCGATCGGCCCGAGGTTATCTAGAGTCACCAAAGCC | 7 |
| AH-h28S-7 | AGACGGGCCGGTGGTGCGCCCTCGGCGGACTGGAGAGGCCTCGGGATCCCACCTCGGCCG | 7 |
| AH-h28S-12 | CAAACCCTTGTGTCGAGGGCTGACTTTCAATAGATCGCAGCGAGG | 18 |
| AH-h5.8S-13 | AAGCGACGCTCAGACAGGCGTAGCCCCGGGAGGA | 7 |
| AH-h-sub-3-R | ATTCCATTATTCCTAGCTGCGGTATCCAGGCGGCTC | 7 |
| AH-h-sub-4-R | GGGCCTCGATCAGAAGGACTTGGGCCCCCCACGA | 7 |
| AH-h-sub-5-R | TCGCTCCCTCGGCCCCGGGATTCGGCGAGTGC | 7 |
| AH-h-sub-7-R | CGCGCCGTGGGAGGGGTGGCCCGGCCCC | 7 |
| AH-h-sub-12-R | GCGGGGGACCGGCTATCCGAGGCCAACCGAGGCTC | 7 |
